# Two liverworts from same habitats developed many similar but few distinct seasonal adaptive strategies: Insights from a transcriptomic approach

**DOI:** 10.1101/2024.07.12.603233

**Authors:** Suvajit Basu, Sandhya Yadav, Vishal Kumar Jha, Subhankar Biswas, Akanksha Srivastava, Kritika Tripathi, Raju Mondal, Neha Chaurasia, Sushil Kumar Singh, Yogesh Mishra

## Abstract

Liverworts are among the first land plants to experience seasonal changes after terrestrialization Therefore, we conducted a seasonal transcriptome analysis of two representative liverwort species from India, *Dumortiera hirsuta* and *Plagiochasma appendiculatum*, during their distinct growing seasons: the pre-monsoon, monsoon, post-monsoon, and fruiting season to understand the seasonal adaptations. Phylogenetic trees and evolutionary timescale analyses showed that *D. hirsuta* is primitive compared to *P. appendiculatum*. The RNA-seq analysis showed that *D. hirsuta* primarily modifies its transcriptome in the post-monsoon season but mainly induces specific genes in the fruiting season, most likely to develop reproductive organs and to adapt strategically by conserving energy in the fruiting season to deal with the harsh environmental conditions of both seasons. Conversely, *P. appendiculatum* exhibited significant transcriptome variability during both the fruiting and post-monsoon seasons, albeit to a lesser degree than *D. hirsuta*. This suggests that, to survive in the harsh conditions of both seasons, it strategically modulated its necessary gene expression levels over an extended period of time while taking energy conservation into consideration. This study offers the first comprehensive view of seasonal adaptive strategies employed by two liverworts that coexist in the same habitat but diverged at different stages in their evolutionary history.

## 1 INTRODUCTION

Seasonality represents the strongest and most ubiquitous source of external variations influencing the behavior of organisms on the Earth. Seasonal fluctuating environments impose selection on life-history traits that can elicit various adaptive responses in organisms (Varpe, 2017). Depending on the geographical locations, plants endure remarkably different growing seasons where they are influenced by several key seasonal factors such as photoperiod, temperature, precipitation, humidity, and nutrient availability (Walker et al., 2019). Stresses caused by seasonal changes in these key factors are accentuated in plants due to their sessile nature. Consequently, plants developed several tolerance mechanisms to combat environmental pressures imposed by seasonality. To gain insight into these adaptive mechanisms, several research at the ecological, morphological, physiological, biochemical, and molecular levels have been conducted in a wide variety of plants (Singh and Yadava, 1974; Chen et al., 2006; Shao et al., 2008; Vuleta et al., 2010; Angelcheva et al., 2014; Leelahawong et al., 2016; Sheikh et al., 2017; Aoussar et al., 2020). The molecular aspects of seasonality have been deeply investigated through the use of some “omics” technologies such as transcriptomics, metabolomics, and proteomics (Angelcheva et al., 2014; Budzinski et al., 2016; Jokipii-Lukkari et al., 2018; Nagano et al., 2019; Lu et al., 2020; Bag et al., 2020; Grebe et al., 2020; Sobreiro et al., 2021).

A group of non-vascular, cryptogamic, seedless plants named as bryophytes, have contributed significantly to the plant diversity with *c*. 20,000 species worldwide, making them the second most diverse land plant group after angiosperms. Bryophytes are a monophyletic plant lineage that is sister to all other land plants (tracheophytes) (Bowman et al., 2017). They have three evolutionary lineages: mosses, liverworts, and hornworts (Shaw and Renzaglia, 2004; Puttick et al., 2018). Although these bryophytes lineages have long faced environmental challenges related to seasonality, research on seasonality is not as extensive as that on flowering plants. From the 1940s to 2000s, there have been some studies on seasonality in bryophytes; but these studies have mainly focused on the morphological and/or anatomical changes (Lackner, 1939; Tamm, 1953; Pitkin, 1975; Rincon and Grime, 1989; Hanslin, 1999). In the past few years, some research has been done on analyzing the effect of seasonality on metabolomics, flavonoid and phenolic content, polyphenol oxidase levels, and antioxidant contents (Thakur and Kapila, 2016; Thakur and Kapila, 2017; Peters et al., 2018; Yadav et al., 2022). However, these studies do not seem to offer sufficient details regarding the seasonal adaptation mechanisms of bryophytes.

Liverworts (Marchantiophyta) are a prominent and important part of many terrestrial ecosystems all over the world. The number of different species of liverworts is extremely large in India (von Konrat et al., 2008), where they have endured seasonal environmental challenges over millions of years. Since liverworts were among the first land colonizers, they are crucial to our understanding of how early terrestrial plants evolved to survive and adapt to the many abiotic stresses brought on by seasonal changes (Bowman et al., 2017). To early colonize on the land and to adapt to the varying seasons, they might have developed a number of defense mechanisms at molecular scale. But little is known about it. To provide insight into these molecular mechanisms, it is crucial to conduct seasonality-based investigations at the gene expression level. Elucidation of the seasonality-driven gene expression modulations may be possible if RNA-sequencing is employed on naturally-flourished plants.

Although large scale RNA sequencing have not been applied in bryophytes in relation to seasonality but it has been widely employed in two model bryophyte systems (*Physcomitrella patens*, and *Marchantia polymorpha*) as well as some non-model species to understand their adaptation to diverse abiotic stresses (Oliver et al., 2004; Xiao et al., 2011; Sharma et al., 2014; Khraiwesh et al., 2015; Singh et al., 2015; Szövényi et al., 2015; Kumar et al., 2016; Gao et al., 2017; Kubo et al., 2019; Elzanati et al., 2020).

Given the lack of transcriptome-based studies with respect to seasonality in liverworts, we have employed RNA-sequencing (RNA-seq) in two non-model liverwort species: *Dumortiera hirsuta* and *Plagiochasma appendiculatum* during their different growing seasons. These liverwort species were selected because of their different evolutionary histories. *D. hirsuta* emerged approximately 145 million years ago (MYA) during the Cretaceous period of the Mesozoic era, whereas *P. appendiculatum* originated about 25 MYA during the Paleogene period of the Cenozoic era (Kumar et al., 2017) [Fig.1(a)]. Moreover, these two species are widely distributed throughout India and flourish in various regions (both Eastern and Western Himalayan, and Western Ghats), where they experience four distinct growing seasons: pre-monsoon (summer), monsoon, post-monsoon (autumn), and fruiting (winter/fertile) period We then analysed and compared the seasonal transcriptome data from the RNA-seq analysis of these two liverworts in order to understand the molecular basis of liverwort seasonal adaptation and to identify any similarities or differences in seasonality-driven gene-expressions changes at the species level.

**Figure 1.**
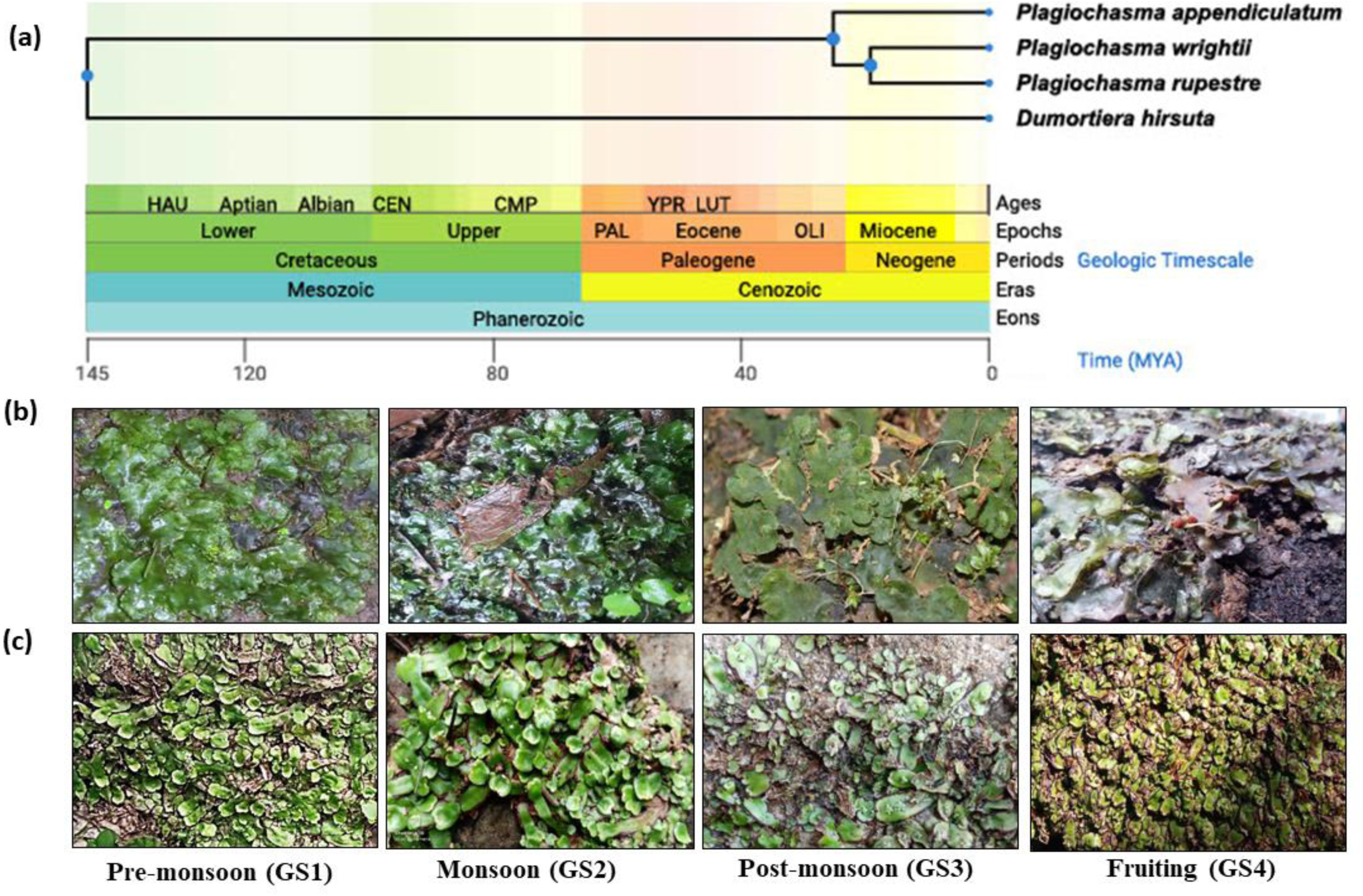
**(a)** The phylogenetic tree and evolutionary timescale representing the divergence time of *D. hirsuta* (145 MYA) and *P. appendiculatum* (25 MYA). **(b)** and **(c)** Images of the gametophytes of **(b)** *D. hirsuta,* **(c)** *P. appendiculatum* in their natural habitat during the course of four different growing seasons (GS), namely pre-monsoon as GS1, monsoon as GS2, post-monsoon as GS3, and fruiting season as GS4. MYA; Million years ago.

## 2 MATERIALS AND METHODS

### 2.1 Phylogenetic tree and evolutionary timescale

The phylogenetic tree and evolutionary timescale (geological time and astronomical history) was analyzed using TIMETREE v5 (http://www.timetree.org; Kumar et al., 2017) resources.

### 2.2 Plant Materials and collection site

The thalli of *D. hirsuta* and *P. appendiculatum* were collected from four different seasons of the year 2019 and 2020 as per Yadav et al. (2022) from the Pynursla town (located at latitude 25.5788°N and longitude 91.8933°E) of the East Khasi Hill district, Shillong, Meghalaya, India and from Botanical Survey of India (BSI) campus in Shillong, respectively. Shillong has an average annual temperature of 17.1 °C and an average annual precipitation of 3385 mm. This region experiences mostly humid weather throughout the year, except for the relatively dry spell usually between December and March. The maximum temperature in this region seldom rises above 28 °C. In relation to the life histories of liverworts, this region has four contrasting growing seasons (GS): pre-monsoon (March–May) or GS1, monsoon (June–August) or GS2, post-monsoon (September–November) or GS3, and sporophyte or fruiting-body producing/fertile season (December–February) or GS4 (henceforth referred to as the fruiting season throughout the manuscript). Figs. 1(b) and (c), respectively, show the thalli of *D*. *hirsuta* and *P*. *appendiculatum* from all four seasons.

### 2.3 Experimental designing, RNA extraction, library preparation and sequencing

Total RNA was extracted from four contrasting growing seasons of *D*. *hirsuta* and *P*. *appendiculatum* in three biological replicates (a total of 12 samples per organism - 4 growing seasons and 3 replicates), as detailed in Supplementary file 1: Figure S1. Total RNA was isolated using TRIzol reagent (Invitrogen, USA) following the manufacturer’s instructions. The RNA-seq libraries were created according to the manufacturer’s instructions using the Tru-Seq RNA sample prep kit (Illumina, Inc.). In brief, the Tru-Seq RNA sample prep kit uses poly-T oligo-attached magnetic beads to convert poly-A containing mRNA in total RNA into a cDNA library. After mRNA purification, the RNA is chemically fragmented before reverse transcription and cDNA synthesis. The fragmentation process results in an RNA-seq collection with inserts ranging in size from 100 to 400 bp. In an Illumina Tru-Seq RNA sequencing library, the typical insert size is around 200 bp. The cDNA fragments are then repaired at the ends by adding a single ’A’ base to the 3’ end, followed by adapter ligation. The products are then purified and amplified with PCR to produce the final double-stranded cDNA libraries. Finally, libraries were quality controlled and quantified using an Agilent Bio-analyzer Chip DNA 1000 series II before being sequenced directly utilizing the high-throughput Illumina HiSeq sequencing equipment (Illumina, Inc.). The raw data files were submitted to the NCBI SRA database under the Bio Project PRJNA1117754 (http://www.ncbi.nlm.nih.gov/bioproject/1117754).

### 2.4 *De novo* transcriptome assembly and annotation of peptide sequences

The data from all the samples were digitally normalized using BBNorm (https://github.com/BioInfoTools/BBMap/blob/master/sh/bbnorm.sh) at K=25, to achieve a kmer-depth of 100. Before data processing, reads with a low-quality score and fewer than 50 bp in length were deleted using QTrim (https://bioinformaticshome.com/tools/rna-seq/descriptions/QTrim.html). The transcriptome was assembled using Trinity with *k-mer* K=25. The assemblies were concatenated and were taken for filtering and identifying the true transcripts. Filtering the false transcripts from the true transcripts was done using the Evidential gene packages: tr2aacds.pl, and retained if the minimal CDS was 90 bp in length. The designed package followed the pipeline in which first redundancy was removed from all the reads by “Perfect redundant removal: fastanrdb (from exonerate package)”. Then, the fragments less than 90 bp in length were removed by “Perfect fragment removal: cd-hit-est -c 1.0”. Ultimately, the *de novo* assembly was completed by Blast searches and annotation of fragments with the following method; “Blastn: identifying the highly identical transcripts” followed by, “Classify main/alternate cds, okay & drop subsets, using evigene/rnaseq/asmrna_dupfilter2.pl”. The above-mentioned pipeline provided an “okay-set” of transcripts in the end, which passed the filters of the pipeline. These peptide sequences were annotated to know their function. For the purpose of differential expression and annotation, we have merged “okay” transcripts and their alternative isoforms. BLASTx v2.2.25 was used to blast transcriptome assemblies against the Uniprot database (The UniProt Consortium, 2014) (Altschul et al., 1997). BLAST outputs were filtered to eliminate transcripts with top hits to non-land plants in order to remove transcripts assembled from contaminated species. The Trinity package’s ’transcripts to best scoring ORFs’ perl script was used to make ORF predictions for all transcripts that passed these filtering stages. The longest, highest-scoring projected ORF with at least 150 amino acids was chosen for further downstream analysis for each transcript.

### 2.5 Annotations with different standard genome databases

All transcripts (contigs and singletons) were compared to the NCBI non-redundant protein database (nr; ftp://ftp.ncbi.nih.gov/blast/db/FASTA/) using BLASTx with rigorous settings (e-value 1010), and the best hits were chosen for further investigation. These transcripts were also searched using BLASTx against the TAIR database (ftp://ftp.arabidopsis.org /home /tair/ Sequences/blastdatasets/ TAIR10 blastsets/). TAIR locus IDs were allocated to all transcripts that shared similarities with the TAIR10 protein database. Similarly, the unigenes were searched using BLASTx against *Physcomitrium patens* genome database (https://www.plantgdb.org/PpGDB/) and *Marchantia polymorpha* genome database (https://www.marchantia.org/marchantia-genome).

### 2.6 Differential gene expression analysis

The *de novo* transcriptome analysis of the *D. hirsuta* and *P. appendiculatum* led to the differential expression and comparison among the samples. The filtered transcript set was indexed using Salmon in order to map the reads from the samples and to derive the respective transcript counts. Salmon uses a quasi-alignment approach where in the read-kmers are matched against the reference, deriving the counts transcript-wise. The counts are rounded off to the closest integers and are used for further analyses in DESeq2 (Love et al., 2014). Thresholds used in case of DESeq2 are |log2FC|>1 and *p*-adjusted value <0.05. The analyses have been conducted by clustering the transcripts to genes and a differentially expressed gene (DEG) analysis was then performed. To represent the differentially regulated transcripts for both liverworts from a wholesome perspective, heatmaps were generated from their RNA-seq data of all three replicates of each of the four seasons. The raw expression values were segregated accordingly and heatmap package from R was used for generating the heatmaps. To observe the uniqueness of the transcriptome data among different growing seasons, we also carried out the Principal Component Analysis (PCA) of all three replicates of the four seasons as well as their average.

### 2.7 Functional annotation, gene-ontology (GO), and pathway enrichment analysis

This was achieved by retrieving the gene sequences from the *de novo* reference using a Perl module within the trinity pipeline. Subsequently, BLASTX was performed against the non-redundant (nr) database, and InterProScan was utilized for domain analysis. Furthermore, GO annotation was conducted using BLAST2GO (https://www.blast2go.org/) software (version 5 Basic).

Following annotation, we selected the GO IDs and employed an in-house Python program to obtain the GO descriptions. This allowed us to count the frequency of occurrence of each GO term, facilitating a comprehensive understanding of the functional annotations. Subsequently, a combined GO plot was generated to visualize the enriched biological processes, molecular functions, and cellular components associated with the DEGs. Cumulative bar graphs were created and the GO terms were divided into up and downregulated transcript pools between the comparative growing seasons for both the organisms.

To further explore the functional significance of the DEGs, pathway enrichment analysis was carried out using KOBAS 3.0. Utilizing the gene FASTA sequences as input, KOBAS identified significantly enriched pathways, providing insights into the biological pathways potentially impacted by the observed differential expression patterns. This comprehensive annotation and pathway analysis pipeline enabled a deeper understanding of the biological processes underlying the observed transcriptional changes.

### 2.8 Prediction of putative transcription factor families

The *de novo* assembly of transcriptome was subjected for the identification of putative transcription factor families present in both the liverworts. The sequence data were subjected to nucleotide BLAST analysis with the PlnTFDB (https://planttfdb.gao-lab.org/prediction.php) database (Jin et al., 2016). Transcription factor families obtained for *D. hirsuta* and *P. appendiculatum* transcripts were depicted in the form of bar graphs. The analysis was done on the complete annotated data along with all the season vs season comparisons.

### 2.9 Biological network analysis

The Biological Networks Gene Ontology tool (BiNGO) (https://apps.cytoscape.org/apps/bingo) was used to identify the over-representation of GO keywords in a group of genes (Maere et al., 2005). BiNGO extracted the appropriate GO annotation before testing for significance with the Hypergeometric test and correcting multiple testing with the Benjamini and Hochberg false discovery rate (FDR) correction. To assess the similarity of enrichment sets A and B, the Jaccard coefficient was utilized. It was defined as the intersection of groups A and B split by their union, and the BiNGO findings were then shown using the enrichment map. The Gene Set Enrichment Analysis (GSEA) findings from the BiNGO plug-in were shown using Cytoscape (https://cytoscape.org/) and an enrichment map. These enrichment maps or networks were only created for season vs season comparison datasets for D. *hirsuta* and *P. appendiculatum* As seasonality imposes several abiotic stresses on plants, the clusters concerning abiotic stress response/ abiotic stimuli response/ stress or stimuli response were isolated to identify specific putative transcripts present on those nodes. These specific transcripts were selected from the DEG list of several season-to-season comparisons, and Morpheus was used to create heatmaps based on the transcripts’ normalized expression levels (https://software.broadinstitute.org/morpheus/). Blastx was further run on the selected transcripts to identify the putative names of the genes or their relevant functions.

### 2.10 Validation of RNA-Seq data

The RNA-Seq data was validated by semi-quantitative PCR (semi-qRT-PCR) and quantitative Real-Time PCR (qRT-PCR). To do this, a total of ten transcripts were randomly picked from the DEGs among all the season vs season comparisons. Actin and Glyceraldehyde 3-Phosphate Dehydrogenase (GAPDH) sequences were fished from the de novo assembly data by Blastx analysis and were used as controls (reference genes) for the PCR reactions. Primers of these transcripts were created using Primer3Plus software (https://www.bioinformatics.nl/cgi-bin/primer3plus/primer3plus.cgi). Total RNA was extracted and subsequently subjected to semi-qRT-PCR and qRT-PCR, as was previously described in Mondal et al., (2021). The details of all the primers used in this study are provided in Supplementary file 1: Table S1 and S2.

## 3 RESULTS

3.1 *D. hirsuta* is primitive compared to *P. appendiculatum*

According to phylogenetic trees and evolutionary timescale analyses, *D*. *hirsuta* is roughly 25 million years ago significantly more primitive than *P*. *appendiculatum*, having evolved during the Cretaceous period of the Mesozoic era, or about 145 million years ago, whereas *P*. *appendiculatum* originated during the Cenozoic era, or roughly 25 million years ago, during the Paleogene period (Fig. 1a).

### 3.2 The overview of transcriptome sequencing and *de novo* assembly

RNA-seq analysis was carried out on *D*. *hirsuta* and *P*. *appendiculatum* from their four distinct growing seasons (GS1-GS4) that were collected in the year 2019-2020 in three biological replicates (12 samples per organism; Supplementary file 1: Fig. S1).

The three *D*. *hirsuta* replicates from GS1 yielded total reads of 51118586, 47886094, 54094010 and the total generated data was of 7.12, 7.231, 6.17 Gb, respectively. The GS2 triplicates generated reads 63025429, 48343141, and 62240067, while the corresponding genomic data was 6.99, 6.82, and 6.26 Gb. The GS3 triplicates generated total reads of 61019576, 49887897, 56064089 and the total generated genomic data of 7.02, 7.18, and 6.264 Gb. Finally, in GS4 triplicates, the obtained reads were 64521456, 47652491, 61274563 and comprised of 6.56, 6.56, 6.27 Gb sequence data, respectively. Furthermore, in the case of *P*. *appendiculatum*, three GS1 replicates generated total reads of 66026428, 58353042, and 61240064 as well as total data of 6.97, 6.81, and 6.25 Gb, respectively. The GS2 triplicates provided 6.2555486, 46581331, and 62642080 reads, along with 6.96, 6.56, and 6.86 Gb sequence data. The GS3 triplicates contained 66119728, 48987865, and 55874259 total reads with 7.02, 7.25, and 6.66 Gb data. Finally, the GS4 included 64589325, 56278942, 55789423 reads with 7.10, 7.56, 6.55 Gb sequence data, respectively. The Trinity assembly provided 1648079 number of sequences in total, which had a summed-up length of 1391990105 bp. Minimum length of assembled sequences came to 177 bp, whereas the maximum length of the sequence was 48514 bp. Average length of the sequence came to 844.6 bp.

### 3.3 Major transcriptome remodeling was observed in both liverworts during the post-monsoon (GS3) and fruiting seasons (GS4)

To determine the common and specific expressed transcripts in the four different growing seasons of *D*. *hirsuta* and *P*. *appendiculatum*, Venn diagrams (Figs. 2a and d) were generated. In *D*. *hirsuta*, 27466 transcripts out of 55232 were present during all growing seasons, with GS4 having the greatest number of unique transcripts (7390) and GS2 having the lowest number (2781) (Fig. 2a). Similarly, for *P*. *appendiculatum*, of the 107177 transcripts, GS4 had the highest number of unique transcripts (29541), GS2 the lowest number of specific transcripts (4690) and 4058 transcripts expressed during all growing seasons (Fig. 2 d).

**Figure 2.**
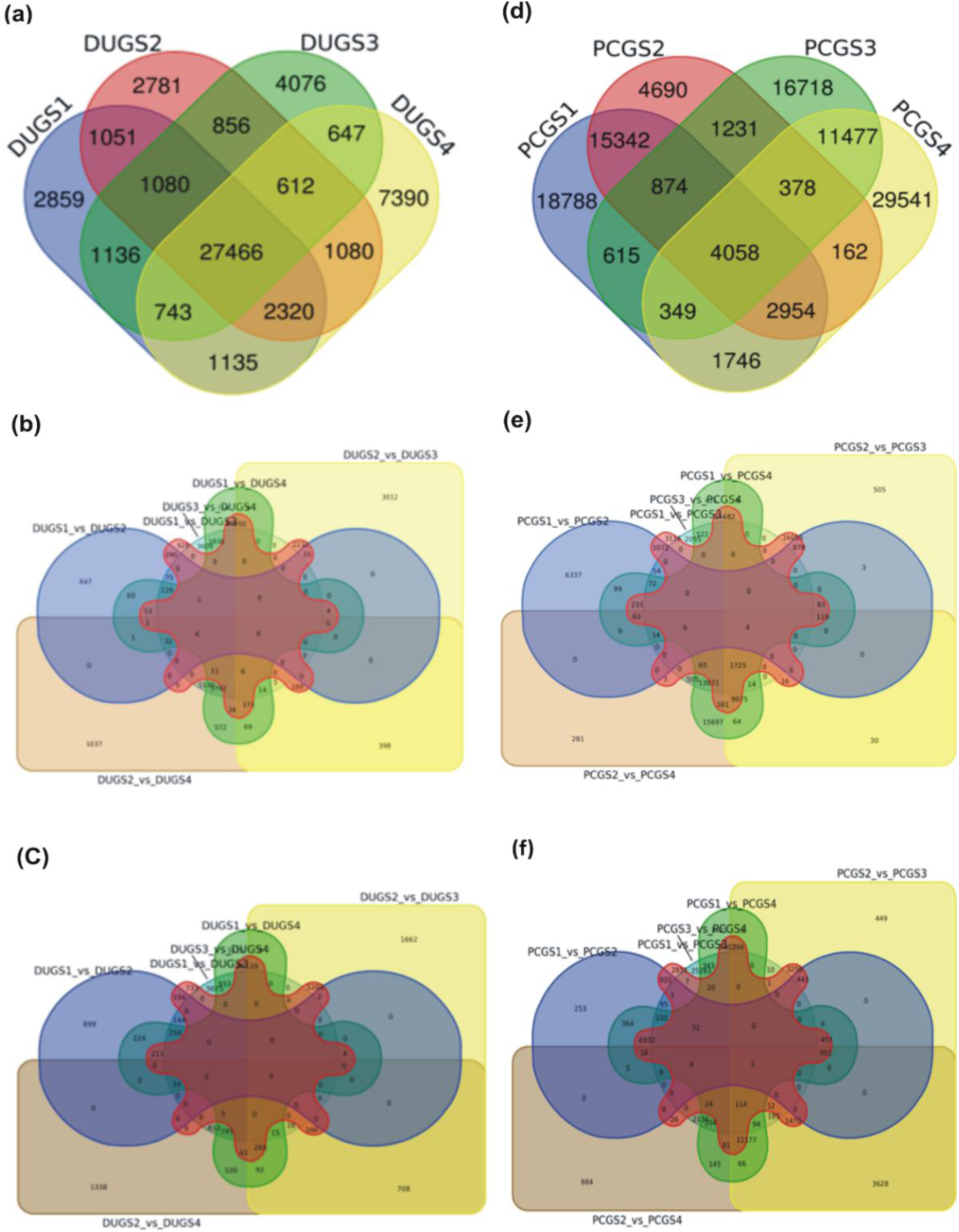
**(a)** and **(d)** Venn diagrams depicting the number of commonly and uniquely expressing transcripts in the four different growing seasons of **(a)** *D. hirsuta* and **(d)** *P. appendiculatum*. Here **(a)** and **(d)** depicts, the dispersion of all obtained transcripts in the complete transcriptome dataset. **(b-f)** Venn diagrams depicting the number of commonly and uniquely expressing **(b,e)** up- and **(c,f)** downregulated transcripts in each of the six paired-wise comparisons of all four seasons in **(b,c)** *D. hirsuta* and **(e,f)** *P. appendiculatum*. DU indicates *D. hirsuta,* while PC indicates *P. appendiculatum*. GS1 is pre-monsoon, GS2 is monsoon, GS3 is post-monsoon, and GS4 is fruiting seasons.

In order to determine the total transcripts that were up- and down-regulated in both liverwort species during the four seasons, six pairwise comparisons were conducted (Supplementary file 1: Table S3). In *D*. *hirsuta*, the GS1 vs GS2 comparison group had 1531 upregulated and 1978 downregulated transcripts. GS1 vs GS3 group had 4110 up and 3138 downregulated transcripts. GS1 vs GS4 had 7721 up and 4778 down-regulated transcripts, GS2 vs GS3 had 6185 up and 4192 downregulated transcripts and GS3 vs GS4 depicted 10638 up and 8653 downregulated transcripts. Lastly, the GS2 vs GS4 had 7469 up and 4783 downregulated transcripts. Among all the comparisons, the maximum variability in the expression of transcripts was observed in the GS3 vs GS4 and GS1 vs GS4 comparison sets, whereas, the GS1 vs GS2 dataset had the least variability in terms of expression levels of transcripts (Supplementary file 1: Table S3). Venn diagrams of up- and downregulated transcripts in each of the six paired-wise comparisons of all four seasons in *D*. *hirsuta* showed that GS3 vs GS4 had the greatest number of unique up- and downregulated transcripts, while GS1 vs GS3 had the lowest number (Figs. 2b and c). In case of *P*. *appendiculatum*, the GS1 vs GS2 comparison group had 9047 upregulated and 10649 downregulated transcripts. GS1 vs GS3 group had 33805 up and 36479 downregulated transcripts. GS1 vs GS4 had 43617 up and 30617 down-regulated transcripts, GS2 vs GS3 had 29561 up and 22583 downregulated transcripts, GS3 vs GS4 depicted 18870 up and 29081 downregulated transcripts. Finally, the GS2 vs GS4 had 42589 up and 21590 downregulated. Among all the comparisons, the GS1 vs GS4 and GS1 vs GS3 groups had the most variability in terms of differentially expressed transcripts, whereas, the GS1 vs GS2 were least variable in this aspect (Supplementary file 1: Table S3). Venn diagrams of up- and downregulated transcripts in each of the six paired-wise comparisons of four seasons in *P*. *appendiculatum* showed that GS3 vs GS4 had the greatest number of unique up- and downregulated transcripts, while GS2 vs GS3 had the lowest number (Figs. 2e and f).

Figs. 3a and c show heatmaps with hierarchical clustering of all differentially expressed transcripts in three biological replicates of each of the four seasons in both organisms. To make the visualization of the transcript expression levels simpler and to identify the seasons that cluster together (due to similar gene expression), the expression datasets of the three replicates were further averaged and additional heatmaps were created (Figs. 3b and d). The *D*. *hirsuta* averaged heatmap revealed that, of the four seasons, GS3 differed significantly from GS1, GS2, and GS4, which all three descended from the same parent and were further split into GS1-GS2 and GS4 (Fig. 3b). The averaged heatmap of *P*. *appendiculatum* revealed that, after deriving from the same parent, GS1 and GS2 formed one group whereas GS3 and GS4 formed another (Fig. 3d). PCA analysis further supported and clarified the clustering data of all four seasons based on the differential gene expression in both liverworts (Figs. 4a-d). Heatmaps of the transcripts that were differentially expressed in each of the six pairwise comparisons of the four seasons were provided in Supplementary file 1: Figs. S2 and S3. These again showed the maximum variation in transcripts expression in GS3 vs GS4 comparisons for *D*. *hirsuta* and in GS1 vs GS4 comparisons for *P*. *appendiculatum*. Volcano plots of significantly up- and downregulated transcripts in six paired-wise comparisons of all four seasons for both species are shown in Supplementary file 1: Figs. S4 and S5.

**Figure 3.**
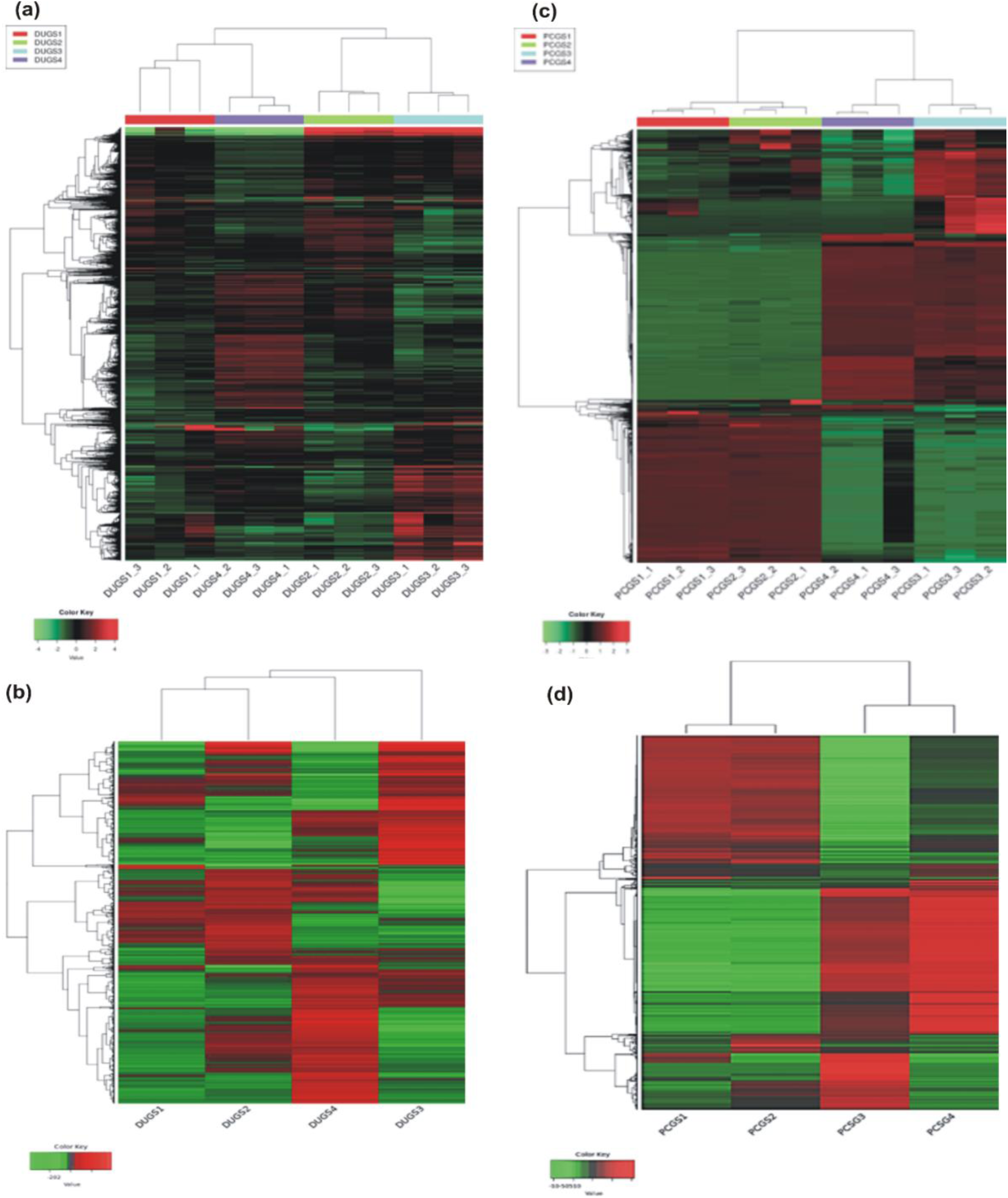
**(a)** and **(c)** showing the heatmaps with hierarchical clustering of all differentially expressed transcripts in three biological replicates of each of the four growing seasons (GS) in **(a)** *D. hirsuta* and **(c)** *P. appendiculatum*. **(b)** and **(d)** depicting the heatmaps with hierarchical clustering of all differentially expressed transcripts, which are the average of three biological replicates, identified during all four growing seasons of **(b)** *D. hirsuta* and **(d)** *P. appendiculatum*. DU indicates *D. hirsuta,* while PC indicates *P. appendiculatum*. GS1 is pre-monsoon, GS2 is monsoon, GS3 is post-monsoon, and GS4 is fruiting seasons. The color green depicts the downregulated transcripts and red depicts the upregulated transcripts.

**Figure 4.**
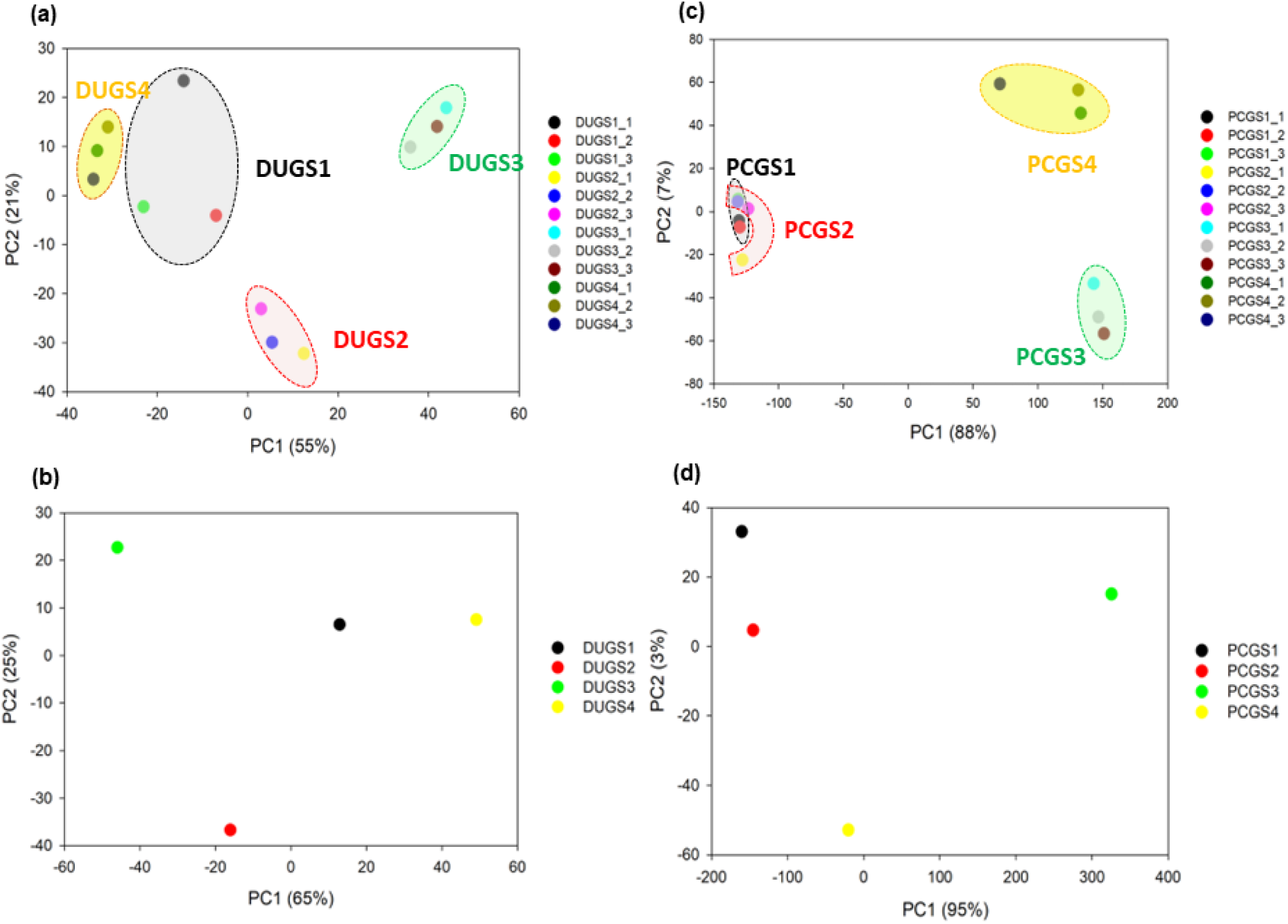
Principal Component Analysis of the all four growing seasons (GS) of **(a,b)** *D. hirsuta* and **(c,d)** *P*. *appendiculatum* in each of their three biological replicates **(a,c)** as well as in the average of those three biological replicates **(b,d)**. DU indicates *D. hirsuta,* while PC indicates *P. appendiculatum*. GS1 is pre-monsoon, GS2 is monsoon, GS3 is post-monsoon, and GS4 is fruiting seasons.

### 3.4 GO analysis of DEGs identified during six inter-season comparisons confirmed that *D*. *hirsuta* and *P*. *appendiculatum* primarily use similar seasonal adaptive strategies, with a few differences

To gain an understanding of the major biological events due to seasonal changes in both liverworts, we selected the top 50 GO terms from each of the three categories: biological process, cellular component, and molecular function, by analyzing the DEGs in six paired-wise comparison groups among all four growing seasons (Supplementary file 1: Figs. S6 and S7). But only the top 10 GO terms from each of the three categories (Figs. 5 and 6) are highlighted below.

**Figure 5.**
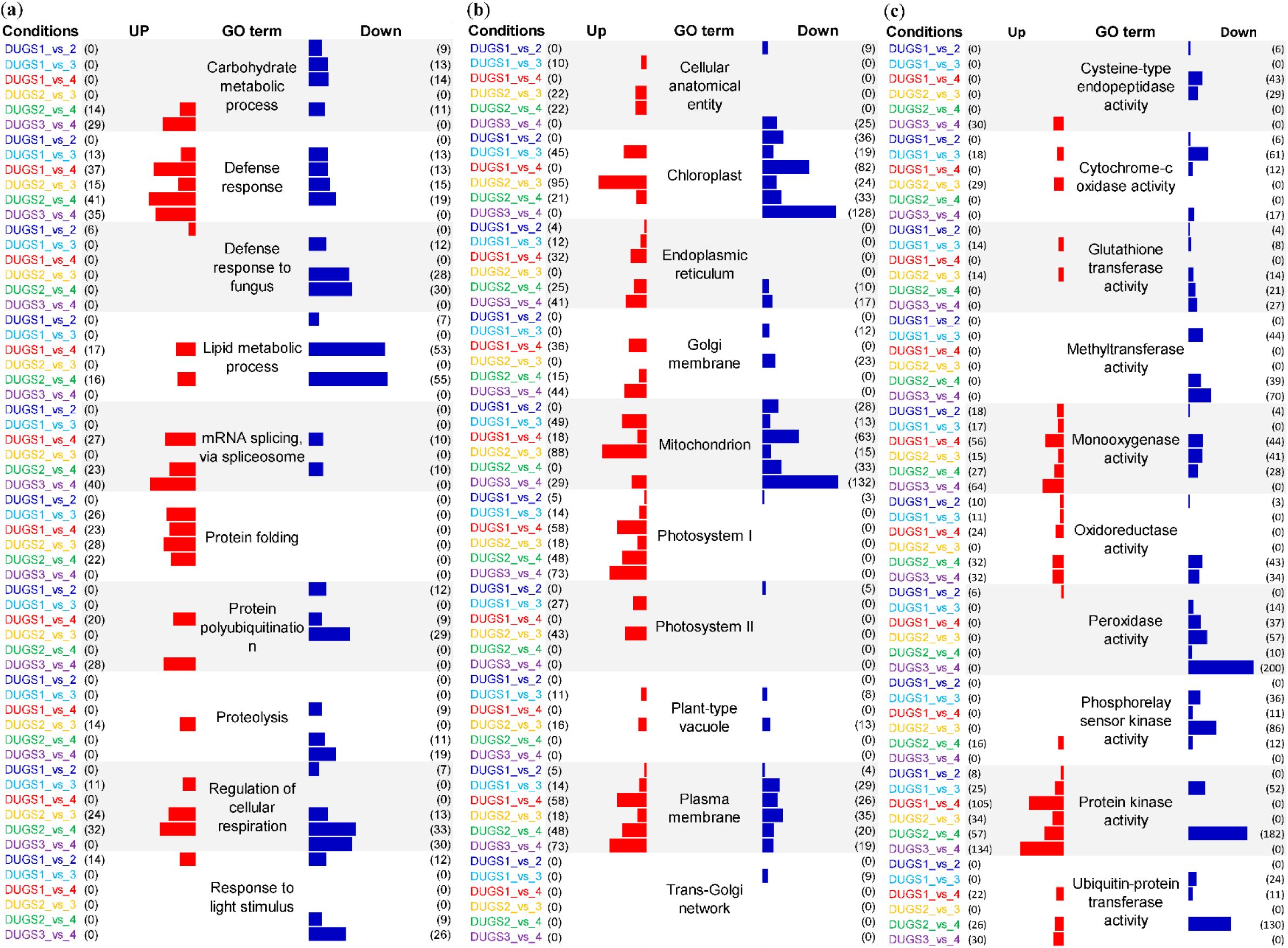
Gene Ontology (GO) enrichment analysis of top ten pathways of **(a)** biological processes (BP), **(b)** cellular components (CC), and **(c)** molecular functions (MF) for the differentially regulated genes of *D. hirsuta* in each of the six paired-wise comparisons of all four seasons. Values inside the parenthesis in front of each bars represents the enrichment counts. DU indicates *D. hirsuta*. GS1 is pre-monsoon, GS2 is monsoon, GS3 is post-monsoon, and GS4 is fruiting seasons.

**Figure 6.**
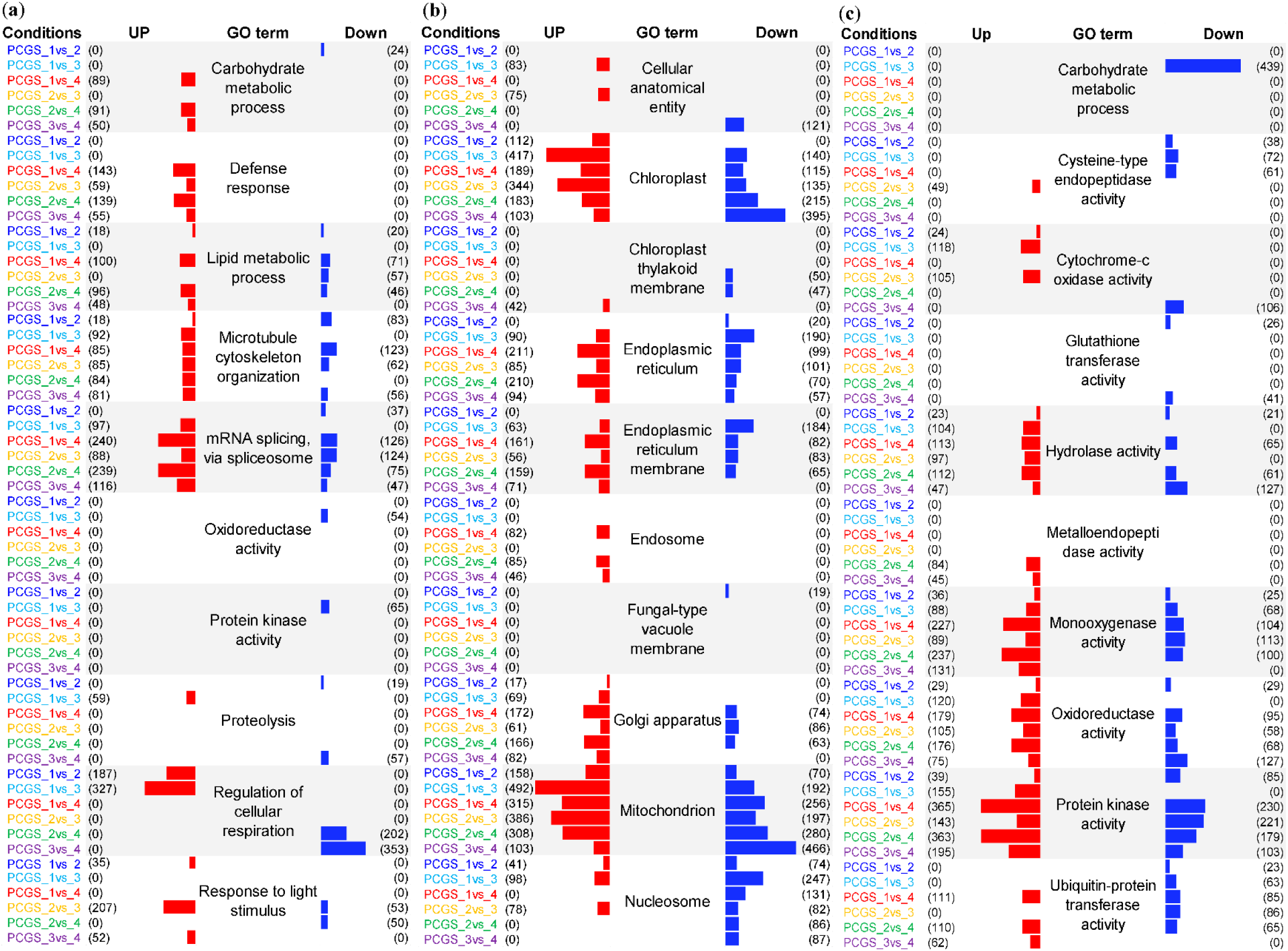
Gene Ontology (GO) enrichment analysis of top ten pathways of **(a)** biological processes (BP), **(b)** cellular components (CC), and **(c)** molecular functions (MF) for the differentially regulated genes of *P. appendiculatum* in each of the six paired-wise comparisons of all four seasons. Values inside the parenthesis in front of each bars represents the enrichment counts. PC indicates *P. appendiculatum*. GS1 is pre-monsoon, GS2 is monsoon, GS3 is post-monsoon, and GS4 is fruiting seasons.

#### 3.4.1 Biological process (BP)

The top 10 GO terms in the BP for *D. hirsuta* included carbohydrate metabolic process, defense response, defense response to fungus, lipid metabolic process, RNA splicing, protein folding, protein polyubiquitination, proteolysis, regulations of cellular perspiration, and response to stimulus (Fig. 5a). However, in the *P. appendiculatum* the top ten GO categories included carbohydrate metabolic process, defense response, lipid metabolic process, microtubule cytoskeleton organization, RNA splicing, oxidoreductase activity, protein kinase activity, proteolysis, regulations of cellular perspiration, response to light stimulations (Fig. 6a). The comparative analysis of BP GO categories of both the organism suggested that two GO categories were unique for *D. hirsuta* i.e. protein folding, protein polyubiquitination while for *P. appendiculatum*, microtubule cytoskeleton organization, oxidoreductase activity, and protein kinase activity were unique.

In the inter-season comparisons of *D. hirsuta*, carbohydrate metabolic-responsive genes were downregulated in GS1 vs GS2, GS1 vs GS3, and GS1 vs GS4 but strongly upregulated in GS2 vs GS4 and GS3 vs GS4 (Fig. 5a). In the case of *P. appendiculatum,* carbohydrate metabolic-responsive genes were downregulated in GS1 vs GS2, mostly upregulated in GS1 vs GS4 and GS2 vs GS4, and only slightly elevated in GS3 vs GS4 (Fig. 6a).

The genes related to defense response in *D*. *hirsuta* showed the most upregulation in GS1 vs GS4, GS2 vs GS4, and GS3 vs GS4 (Fig. 5a) In *P*. *appendiculatum*, the genes related to defence response showed the most expression in GS1 vs GS4 and GS2 vs GS4, but slightly decreased in GS3 vs GS4 (Fig. 6a). However, in *D*. *hirsuta*, certain defence-related genes were downregulated in GS1 vs GS3, GS1 vs GS4, GS2 vs GS3, and GS2 vs GS4; however, in the *P*. *appendiculatum*, these genes were not downregulated in any season vs season comparison group (Figs. 5a and 6a).

Genes related to lipid metabolism were maximally up- and downregulated in both liverworts in GS1 vs GS4 and GS2 vs GS4 (Figs. 5a and 6a). The mRNA splicing associated genes were substantially increased in GS1 vs GS4, GS2 vs GS4, and GS3 vs GS4 in *D*. *hirsuta*, but only maximally increased in GS1 vs GS4 and GS2 vs GS4 of *P*. *appendiculatum* (Figs. 5a and 6a). Genes associated with protein folding, a GO keyword unique to *D*. *hirsuta*, mostly increased across all season comparisons, with the exception of GS1 vs GS2 (Fig. 5a). Protein polyubiquitination genes, which were also specific to *D*. *hirsuta*, were primarily upregulated in GS1 vs GS4 and GS3 vs GS4, but primarily downregulated in GS2 vs GS3 (Fig. 5a). GO keywords unique to *P*. *appendiculatum*, such as genes related to oxidoreductase and protein kinase, were primarily downregulated in GS1 vs GS3 (Fig. 6a). A GO term that is also unique to *P*. *appendiculatum*, microtubule cytoskeleton organisation, was shown to have the highest levels of upregulated genes in all season vs season comparison groups, with the exception of GS1 vs GS2 (Fig. 6a). Genes associated with cellular respiration regulation were predominantly upregulated in GS2 vs GS3 and GS2 vs GS4, while they were predominantly downregulated in GS2 vs GS4 and GS3 vs GS4 of *D*. *hirsuta*; in *P*. *appendiculatum*, these genes were predominantly upregulated in GS1 vs GS3 and downregulated in GS3 vs GS4 (Figs. 5a and 6a). The last top 10 GO term of the BP, the response to light stimulation, mostly had downregulated genes in GS3 vs GS4 of *D*. *hirsuta* and upregulated genes in GS3 vs GS4 of *P*. *appendiculatum* (Figs. 5a and 6a).

#### 3.4.2 Cellular component (CC)

When comparing the top 10 GO categories for CC modified during the seasonal changes, both liverworts had some similarities and differences (Figs. 5b and 6b). For example, both species have four GO categories in common, such as endoplasmic reticulum, mitochondrion, chloroplast, and cellular anatomical entrance, but each species had six distinct GO categories. Among these top ten CC GO terms, the genes associated with the endoplasmic reticulum, Golgi membrane, PSI, and plasma membrane were found to be largely elevated in GS1 vs GS4 and GS3 vs GS4 of *D*. *hirsuta*, whereas genes linked to mitochondria, PSII, and plant type vacuole were primarily upregulated in GS1 vs GS3 and GS2 vs GS3 (Fig. 5b). Other genes related to chloroplast and mitochondria were mostly downregulated in GS1 vs GS4 and GS3 vs GS4 of *D*. *hirsuta* (Fig. 5b). Similar to this, in the case of *P*. *appendiculatum*, genes related to mitochondria and chloroplasts were primarily downregulated in GS1 vs GS4 and mostly elevated in GS1 vs GS3 and GS2 vs GS3 (Fig. 6b). Genes related to the endoplasmic reticulum and Golgi membrane were mostly increased in GS1 vs GS4 and GS2 vs GS4 of *P*. *appendiculatum* (Fig. 6b).

#### 3.4.3 Molecular function (MF)

The GO categories within the MF exhibited the highest degree of similarity, as both liverworts shared seven of the top ten GO categories related with the MF (Figs. 5c and 6c). These categories were cysteine type endopeptidase activity, cytochrome C oxidase activity, glutathione transferase activity, monooxygenase activity, oxidoreductase activity, protein kinase activity, and ubiquitin-protein transferase activity. Methyltransferase activity, peroxidase activity, and phosphorelay sensor kinase activity were unique in *D. hirsuta* while carbohydrate metabolic process, hydrolase activity, and metalloendopeptidase activity were unique in *P. appendiculatum*.

The majority of the genes associated with cysteine type endopeptidase activity were upregulated in GS3 vs GS4 of *D*. *hirsuta* and GS2 vs GS3 of *P. appendiculatum* (Figs. 5c and 6c). Most of the genes related to cytochrome C oxidase activity were increased in the GS1 vs GS3 and GS2 vs GS3 of both liverworts (Figs. 5c and 6c). The genes linked to glutathione transferase activity were mostly increased in *D*. *hirsuta* GS1 vs GS3 and GS2 vs GS3, while they were downregulated in *D*. *hirsuta* GS1 vs GS4 and did not appear to be significantly involved in *P*. *appendiculatum* seasonal adaptation (Figs. 5c and 6c). Methyltransferase activity and peroxidase activity genes, which are GO keywords unique to *D*. *hirsuta*, were shown to be downregulated in all six inter-season comparisons, with GS2 vs GS3 showing the greatest decrease (Fig. 5c). The GO term phosphorelay sensor kinase activity was also unique to *D*. *hirsuta*, and genes related to this activity were mainly upregulated in GS2 vs GS3 (Fig. 5c). In *D*. *hirsuta*, oxidoreductase and ubiquitin-protein transferase activity related genes were primarily upregulated in GS1 vs GS4, GS2 vs GS4, and GS3 vs GS4; in *P*. *appendiculatum*, oxidoreductase-related genes were specifically upregulated in GS1 vs GS4 and GS2 vs GS4 (Figs. 5c and 6c). The majority of the protein kinase related genes were upregulated GS1 vs GS4 and GS3 vs GS4 of *D*. *hirsuta*, and in GS1 vs GS4 and GS2 vs GS4 of *P*. *appendiculatum* (Figs. 5c and 6c). Genes related to hydrolase activity, a GO word unique to *P*. *appendiculatum*, were primarily upregulated in GS1 vs GS3, GS1 vs GS4, GS2 vs GS3, and GS2 vs GS4, and downregulated in GS3 vs GS4 (Fig. 6c). The genes related to metalloendopeptidase activity, a GO word also unique to *P*. *appendiculatum*, were mostly upregulated in GS2 vs GS4 and GS3 vs GS4 (Fig. 6c).

### 3.5 Significant expression changes were also noticed in the expressed transcription factors (TFs) of both liverworts

The TF families derived from the entire transcriptome data of *D*. *hirsuta* and *P*. *appendiculatum* over the course of four growing seasons were represented as bar graphs (Fig. 7), which indicate that *P*. *appendiculatum* has only 39 identified TF families in all inter-season data sets, while *D*. *hirsuta* has 54. A critical examination of the DEGs for these families indicated that *D*. *hirsuta* displayed a more pronounced degree of expression change than *P*. *appendiculatum*, indicating that the TFs in both liverworts function slight differently.

**Figure 7.**
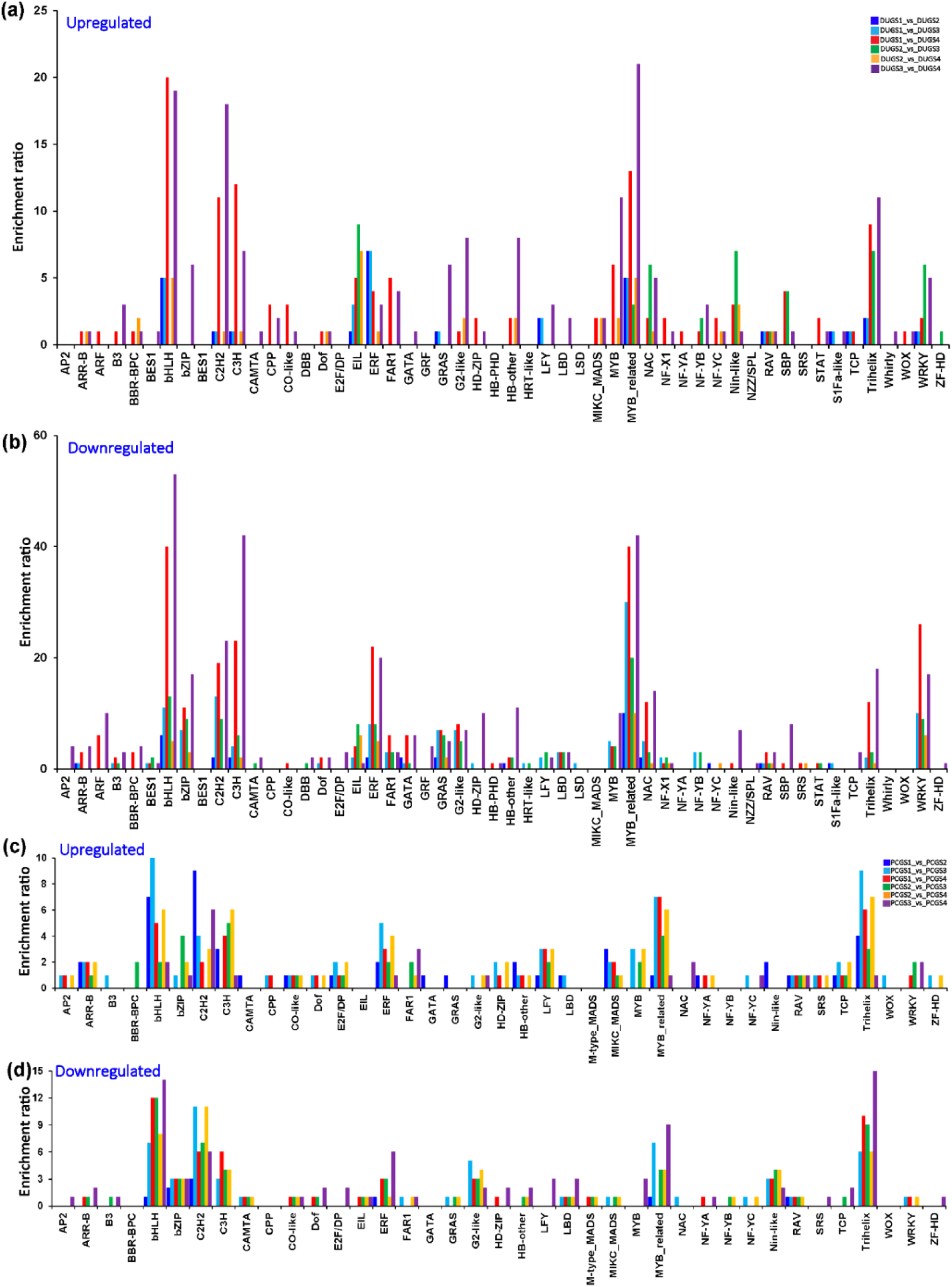
Putative transcription factor (TF) families obtained in the complete transcriptome data of *D. hirsuta* and *P. appendiculatum* during four growing seasons based on PlnTFDB database in total. Figures **(a)** and **(b)** depict the abundance of these TF families in each of the six paired-wise comparisons of all four seasons of **(c)** *D. hirsuta* and **(d)** *P. appendiculatum*. DU indicates *D. hirsuta,* while PC indicates *P. appendiculatum*. GS1 is pre-monsoon, GS2 is monsoon, GS3 is post-monsoon, and GS4 is fruiting seasons.

Overall, *D. hirsuta* TF-related transcript accumulation pattern indicates that the five most prevalent transcription factor families were bHLH, C3H (Cysteine3Histidine), ERF (Ethylene Response Factors), MYB-related (Myeloblastosis-related), and WRKY, which dominated the species throughout all growing seasons (Figs. 7a and b). The members of bHLH, C3H, and MYB-related TF families were highly upregulated in GS1 vs GS4 and GS3 vs GS4, while the ERF TF family members were largely elevated in GS1 vs GS2 and GS1 vs GS3 and downregulated in GS1 vs GS4 and GS3 vs GS4 of *D*. *hirsuta* (Figs. 7a and b). The members of WRKY TF family were mostly upregulated in GS2 vs GS3 and GS3 vs GS4 and downregulated in GS1 vs GS4 and GS3 vs GS4 of *D*. *hirsuta* (Figs.7a and b).

The five most prevalent TF families for *P*. *appendiculatum* were bHLH, C3H, MYB-related, TriHelix, and WRKY, but their patterns of gene expression differed slightly from those of *D*. *hirsuta* (Figs. 7c and d). The members of bHLH TF family were mostly upregulated in GS1 vs GS2 and GS1 vs GS3, and downregulated in GS3 vs GS4 of (Figs. 7c and d). The C3H TF family members were primarily upregulated in GS2 vs GS3 and GS2 vs GS4, and decreased in GS1 vs GS4 (Figs. 7c and d). The members of MYB-related and TriHelix family were mostly upregulated in GS1 vs GS3, GS1 vs GS4, and GS2 vs GS4, and downregulated in GS3 vs GS4 of (Fig. 7c and d). The members of the WRKy were mainly upregulated in GS2 vs GS3 and GS3 vs GS4 (Fig. 7c).

### 3.6 Season-to-season comparison of the gene regulatory networks showed that both liverworts have largely similar but slightly different stress-responsive gene expression pattern

We tried to find the genes in *D. hirsuta* and *P. appendiculatum* that are involved in the regulation of stress tolerance/ stress response/ stimulus response in order to get an elementary understanding of stress adaptation under seasonal fluctuation. In order to achieve this, Gene-Ontology Enrichment analysis (GESA), was applied to transcriptome datasets of both liverworts. Keeping our focus on the stress-specific nodes within these expression networks, we isolated the putative genes linked to those clusters (Supplementary file 1: Figs. S8 and S9).

A thorough examination of the six paired-wise season comparisons suggested that the two liverworts respond to abiotic stresses during seasonal changes in a slight different way.

The GS1 vs GS2 upregulated dataset included ten transcripts in *D. hirsuta* (Supplementary file 1: Fig. S8a). Of these, the genes that were shown to be highly abundant were Inosine triphosphate pyrophosphatase, Cytokinin riboside 5’-monophosphate phosphoribohydrolase LOG7, and Cyclin family protein. Nevertheless, *P. appendiculatum* was found to have 25 upregulated transcripts in GS1 vs GS2, along with a large protein family that included U1 small nuclear ribonucleoprotein, C2H2-like zinc finger protein, Protein kinase superfamily protein, and DEA(D/H)-box RNA helicase family protein (Supplementary file 1: Fig. S9a). Furthermore, compared to *P. appendiculatum*, in the GS1 vs GS2 dataset, there were more down-regulated transcripts in *D. hirsuta* (Supplementary file 1: Figs. S8b and S9b).

The stress-responsive genes in the GS1 vs GS3 showed similar up- and downregulation in both liverworts. In upregulated datasets of GS1 vs GS3, 57 transcripts were found in the case of *D. hirsuta* and 56 transcripts in the *P. appendiculatum* (Supplementary file 1: Figs. S8c and S9c). Out of these 57 transcripts, the genes of udp-glucuronic acid decarboxylase 5, photosystem I reaction center subunit xi, and brefeldin A-inhibited guanine nucleotide-exchange protein 3 were dominants in case of *D. hirsuta* (Supplementary file 1: Fig. S8c). Moreover, the genes of polyubiquitin 4, F-type h^+^-transporting atpase subunit a, probable receptor-like protein kinase, Ribosomal protein L1p/L10e family, 30S ribosomal protein S19, chloroplastic, and C2H2-like zinc finger protein were mainly dominant in *P. appendiculatum* (Supplementary file 1: Fig. S8c). When it came to downregulated datasets, there were 36 down-regulated transcripts for *D. hirsuta* and 38 transcripts for *P. appendiculatum* (Supplementary file 1: Figs. S8d and S9d). Out of these 36 transcripts of *D. hirsuta*, the notable downregulation was noticed for the genes of Chaperonin 60 subunit beta 1, Ethylene-dependent gravitropism-deficient, Molecular chaperone, S-adenosylmethionine synthetase 2, Chaperonin 60 subunit alpha 1, Atp-dependent Clp protease ATP-binding subunit, and methylcrotonoyl-CoA carboxylase beta chain categories (Supplementary file 1: Fig. S8d). For *P. appendiculatum*, the major genes that registered downregulation belongs to Ribosomal protein S27a / Ubiquitin family protein, Cytochrome c oxidase subunit 1, Strong similarity to initiation factor eIF-2, Glycosyltransferase of cazy family gt31 a, Zinc finger C-x8-C-x5-C-x3-H type family protein, DEA(D/H)-box RNA helicase family protein, and Lipolytic acyl hydrolase (LAH) (Supplementary file 1: Fig. S9d).

In GS1 vs GS4, 30 genes were upregulated in *D. hirsuta* and 40 genes were upregulated in *P. appendiculatum*. The upregulated datasets of *D. hirsuta* showed a very significant upregulation for the genes of Cytochrome c oxidase subunit 1, O-fructosyltransferase family protein, Ribosomal protein S8e family protein, and Translation elongation factor efg/ef2 protein categories (Supplementary file 1: Fig. S8e). However, such upregulation was observed for Polyubiquitin 4, Clathrin heavy chain 1, F-type h^+^-transporting atpase subunit a, TCP-1/cpn60 chaperonin family protein, U1 small nuclear ribonucleoprotein 70 kDa, Ubiquitin carboxyl-terminal hydrolase 13, and Galactose oxidase for *P. appendiculatum* (Supplementary file 1: Fig. S9e). In the aforementioned season vs season comparison group, 64 genes were downregulated in *D. hirsuta* and 31 genes in *P. appendiculatum* . The most downregulated genes were U5 small nuclear ribonucleoprotein helicase, ARM repeat protein interacting with ABF2, Histone acetyltransferase of the CBP family for *D. hirsuta* (Supplementary file 1: Fig. S8f). For *P. appendiculatum*, the major downregulated genes belonged to Double-stranded RNA-binding protein 2, Strong similarity to initiation factor eIF-2, DNA-directed RNA polymerase II subunit 1, Calcineurin-like metallo-phosphoesterase superfamily protein, and Glutamate dehydrogenase (Supplementary file 1: Fig. S9f).

In GS2 vs GS3, 37 and 55 genes were found to exhibit up- and downregulation in *D. hirsuta*, respectively. Of these37 upregulated genes, the expression of the following genes was highly significant: aldolase-type TIM barrel family protein, phospho-2-dehydro-3-deoxyheptonate aldolase, chlorophyll a-b binding protein CP29.1, ethylene-responsive transcription factor ERF087, and lactate/malate dehydrogenase family protein (Supplementary file 1: Fig. S8g). Of 55 downregulated genes, phospholipid-transporting ATPase 1, integral membrane protein, and glyceraldehyde-3-phosphate dehydrogenase showed a highly notable change (Supplementary file 1: Fig. S8h). In *P. appendiculatum*, out of 45 upregulated transcripts, nine genes belonged to Polyubiquitin 4, F-type h+-transporting atpase subunit a, Probable receptor-like protein kinase, Vacuolar protein-sorting-associated protein 4, 30S ribosomal protein S19, Adp-ribosylation factor-like protein 5b, Glucose-6-phosphate isomerase 1, chloroplastic, Double-stranded RNA-binding protein 2, and SNARE-like superfamily protein showed significant increase in their transcripts (Supplementary file 1: Fig. S9g). Of 39 dowregulated transcripts in *P. appendiculatum*, strong similarity to initiation factor eIF-2, DNA-directed RNA polymerase II subunit 1, Glycosyltransferase of the Casy family gt31 a, DEA(D/H)-box RNA helicase family protein, and Lipolytic acyl hydrolase (LAH) are the genes associated with the highest abundance (Supplementary file 1: Fig. S9h).

In GS2 vs GS4 of both liverworts, 36 transcripts in *P. appendiculatum* and only ten in *D. hirsuta* were significantly elevated. The downregulated transcripts for *D. hirsuta* and *P. appendiculatum* were 13 and 37, respectively. Aldolase-type TIM barrel family protein, Chaperone protein (htpG family protein), ferredoxin-NADP reductase, and leaf isozyme 1 in *D. hirsuta* are among the members that showed abundance in transcripts (Supplementary file 1: Fig. S8i). Tubulin/FtsZ family protein, F-type h+-transporting atpase subunit a, Probable receptor-like protein kinase, Vacuolar protein-sorting-associated protein 4, Ribosomal protein L1p/L10e family, 30S ribosomal protein S19, chloroplastic, ATP-dependent zinc metalloprotease FTSH 4, mitochondrial, Adp-ribosylation factor-like protein 5b, Cation efflux family protein, Double-stranded RNA-binding protein 2 in *P. appendiculatum* are among the members that showed abundance in transcripts (Supplementary file 1: Fig. S9i). The gene that is associated with the DUF21 protein in *D. hirsuta* showed a significant presence in the downregulated dataset (Supplementary file 1: Fig. S8j) However, in *P. appendiculatum*, the transcripts of seven genes, including eIF-2, 50S ribosomal protein L14, RNA polymerase II subunit 1, glycosyltransferase, and DEA(D/H)-box RNA helicase family protein, significantly decreased (Supplementary file 1: Fig. S9i).

Compared to previous season-to-season comparisons, the GS3 vs GS4 experienced the largest variations in transcript abundance in the *D. hirsuta*. In the upregulated dataset, 66 transcripts shown significant changes; in the downregulated dataset, this number was 94 (Supplementary file 1: Fig. S8k and S8l). The upregulated genes belonged to important vital processes, such as Photosystem II reaction center protein z, Glutamate receptor, ionotropic; Glutamate receptor 3.3, Protein kinase family protein, and Insulinase. The downregulated genes belonged to Tetratricopeptide repeat (TPR)-like superfamily protein ATPase 11, plasma membrane-type transcripts; NADP-binding rossmann-fold superfamily protein; Alpha/beta hydrolase fold-1 Phosphatidylinositol-3,4,5-trisphosphate 3-phosphatase and dual-specificity protein phosphatase PTEN, and Uncharacterized conserved protein UCP031088.

When compared to *D. hirsuta*, the *P. appendiculatum* GS3 vs GS4 comparison did not demonstrate as dramatic shifts in transcript quantity. In the upregulated dataset, 28 transcripts demonstrated differential expression; in the downregulated dataset, this number was 60 (Supplementary file 1: Fig. S9k and S9l). The upregulated genes belonged to Cytochrome c oxidase subunit 1, NADH-ubiquinone oxidoreductase chain 4L, and nucleoside triphosphate hydrolases.The downregulated genes belonged to Double-stranded RNA-binding protein 2, Strong similarity to initiation factor eIF-2, MIF4G domain-containing protein / MA3 domain-containing protein, Calcineurin-like metallo-phosphoesterase superfamily protein, C2H2-like zinc finger protein, Zinc finger C-x8-C-x5-C-x3-H type family protein, Protein kinase superfamily protein, Transcription factor bHLH66, Chorismate synthase, and SNARE-like superfamily protein.

To validate RNA-seq reflected transcript level, we performed qRT-PCR and determined the expression levels of ten selected DEGs from four growing seasons. Consistent with RNA-seq data, qRT-PCR result showed that the expression of most of the genes registered the same expression patterns during the tested growing seasons in both the liverworts (Supplementary file 1: Figs. S10-S12).

## 4 DISCUSSIONS

Since liverworts were among the first terrestrial colonists, they might have evolved a number of unique adaptive strategies to endure seasonality. To understand these seasonal adaptive strategies at the gene level, we conducted RNA-seq analysis of two liverworts, *D*. *hirsuta* and *P*. *appendiculatum*, which are found in the same habitat today but diverged at different times, in their four distinct growing seasons (GS), namely pre-monsoon (GS1), monsoon (GS2), post-monsoon (GS3), and fruiting (GS4) season. In pre-monsoon season, the life cycle of both liverworts starts and vegetative thallus begins to emerge from older decay or spores. In monsoon season, precipitation and humidity elevates that allows both liverworts to grow luxuriously. In post-monsoon season, precipitation decreases and winter begins, the thallus achieves reaches to the adult stage after reaching sexual maturity. In fruiting season, both liverworts complete their sexual reproduction and produce a sporophyte (fruiting body), the temperature decreases significantly, and the environment becomes extremely harsh with limited nutrition.

Using RNA-seq data, Venn diagrams, heatmaps with hierarchical clustering, and PCA of all differentially regulated transcripts in six pairwise comparisons of all four seasons suggested that, for *D*. *hirsuta*, the post-monsoon season had the highest transcriptome modulation, while fruiting season had the highest number of unique up- and downregulated transcripts (Figs. 2-4). In contrast, *P*. *appendiculatum* showed maximum transcriptome modulation throughout both the post-monsoon and fruiting seasons, with the fruiting season exhibiting the highest number of unique up- and downregulated transcripts (Figs. 2-4). In both liverwort species, the transition from pre-monsoon to monsoon season had the lowest transcriptome variability (Figs. 2-4). The greatest transcript modulation observed in the post-monsoon season of both liverworts can be due to the drop in temperature, precipitation, and the growth of their reproductive structures, which are the highly energy- and nutrient-demanding. Further, the reason behind the greatest up- and down-regulation of unique transcripts during the fruiting season could be to cope with the highest level of ROS production in comparison to the other three seasons (Yadav et al., 2022). This ROS production is caused by nutritional deficiencies due to sporophyte production as well as by decreasing temperature, precipitation, and daylength. These two liverworts most likely need specific proteins to cope with this extremely challenging season. Along with these similarities, lowest transcript modulation was seen in both liverworts during the pre-monsoon to monsoon transition. This could be because of the higher humidity and precipitation levels, which enable both liverworts to grow lavishly. Even with these similarities, there were a few differences between the two liverworts: *D*. *hirsuta* displayed maximum transcriptome alteration in the post-monsoon season, which was further reduced in the fruiting season, while *P*. *appendiculatum* displayed maximum transcriptome modulation during both the fruiting and post-monsoon seasons. Interestingly, however, *D*. *hirsuta* had a considerably greater extent of total transcriptome modulation than *P*. *appendiculatum*. These differences in seasonal adaption in both liverwort species could be attributed to their different evolutionary divergence times. The observation of a greater degree of transcriptome modulation in *D*. *hirsuta* during the post-monsoon season could potentially be attributed to two factors: either its comparatively more primitive nature than that of *P*. *appendiculatum* means that it requires a greater amount of differentially regulated proteins to adapt to post-monsoon specific seasonal changes, or it was unable to precisely regulate gene expression to conserve energy, unlike *P*. *appendiculatum*. But *D*. *hirsuta* probably managed to conserve energy during fruiting season by slightly decreasing overall transcriptome modulation and focusing mostly on greatly modifying the expression of fruiting season-specific genes. When compared to *D*. *hirsuta*, *P*. *appendiculatum* showed greater transcriptome variability in both the fruiting and post-monsoon seasons, but to a lesser extent than *D*. *hirsuta*. This suggests that the *P*. *appendiculatum* has been strategically modifying its gene expression levels for an extended period of time while taking energy conservation into account in order to withstand the challenging conditions of both seasons. The hypothesis of the slightly different seasonal adaptive strategy of both liverworts becomes obvious by analysing the seasonal dynamics of top ten GO keywords (Figs. 5 and 6), TFs (Fig. 7), and stress-responsive genes (Supplementary file 1: S8 and S9).

With respect to BP, the seasonal cycle significantly impacted seven major GO terms: carbohydrate metabolic process, defense response, lipid metabolic process, protein folding, proteolysis, regulation of cellular respiration, and response to stimulus (Figs. 5 and 6). The maximum upregulated genes related to these GO terms were found in fruiting season of *D. hirsuta* and in both post-monsoon and fruiting season of *P. appendiculatum*. This was shown by the higher expression of these genes in GS2 vs GS4 and GS3 vs GS4 of *D. hirsuta* (Fig. 5) as well as the higher gene expression in GS1 vs GS4 but lower in GS3 vs GS4 of *P. appendiculatum* (Fig. 6). The reason for the increased expression of these genes during the fruiting season and/or post-monsoon could be to cope with the stress caused by decreased precipitation and temperatures, as well as to supply energy and nutrients for the development of reproductive structures and sporophytes, which house haploid spores for the germination of fresh thallus. The differential expression pattern of these genes was further supported by the expression of associated TFs including bHLH, ERF, C2H2, EIL, and MYB-relate, which exhibit similar expression patterns in different seasons of both organisms (Fig.7). As most food resource are used on producing sporangia during the fruiting season, the thallus of both liverwort species started to dry, as seen by the change in thallus colour (Fig. 1).

The six GO keywords of the CC category, namely mitochondria, endoplasmic reticulum, Golgi membranes, cellular anatomical entity, chloroplast, and thylakoid apparatus were differentially regulated in both liverwort species over the course of the four growing seasons (Figs. 5 and 6). Of these, the majority of chloroplast and/or PSII related genes were increased in both liverworts during the post-monsoon season, possibly to optimise growth and energy production for the formation of reproductive structures or to prepare for migration during the challenging fruiting season. With the exception of PSI genes, which were upregulated (Figs. 5 and 6), these photosynthetic genes were further downregulated during the fruiting season, most likely as a result of the photosystem’s components being severely quenched in the winter (Bag et al., 2020). Since PSI is believed to be more durable than PSII, its component stays high over the winter to keep photosynthesis going at a minimal level through PSI in order to survive Shillong’s chilly winters, when the average temperature stays between 7 and 8°C. Golden2-like, or G2-like, is a transcription factor that is essential for regulating the expression of genes related to photosynthesis and chloroplast growth and development. The seasonal dynamics of G2-like TF expression also support the dynamics of genes related to chloroplasts and photosystem I and II (Fig. 7). The expression of genes related to mitochondria was primarily elevated in both liverworts during the post-monsoon season, while the expression of genes related to endoplasmic reticulum and Golgi membrane were mainly elevated in fruiting season of *D. hirsuta* and in both post-monsoon and fruiting season of *P. appendiculatum*, again indicating a rise in energy needs for the development of reproductive structures and adaptation to unfavourable environmental conditions (Figs. 5 and 6).

The GS3 vs GS4 comparison of *D. hirsuta* revealed an abundance of two significant GO terms that fall under the MF category: monooxygenase activity and protein kinase activity (Fig. 5). The former is known to be vital for plant growth, development, and adaptation, whereas the latter is recognized to be essential for signal transduction in plants. The abundance of these genes in the fruiting season of *D. hirsuta* induces their expression to help it adapt well to the fruiting season-specific changes in the environment. The GS1 vs GS4 comparison of *P. appendiculatum* revealed, maximum upregulation of some important GO terms associated with the MF, such as translation elongation activity, monooxygenase activity, oxidoreductase activity, and protein kinase activity (Fig. 6). As the aforementioned GO categories are associated with development, defense, and signaling, their GO enrichment pattern suggests a strategy for preparing *P. appendiculatum* to meet the challenges associated with fruiting season. The seasonal dynamics of the MF GO keywords described above are supported by the related TF families, which include EIL, NAC, Tri-Helix, Nin-Like, and WRKY (Fig. 7).

In addition to understanding the seasonal physiological and metabolic adjustments, we focused on understanding how these two liverworts respond to season-specific abiotic stresses by differentially regulating unique stress-responsive genes. Gene regulatory networks study of DEGs observed in each of the six paired-wise comparisons of all four seasons in both liverworts using GSEA allowed us to focus on the nodes in the networks that involved with stress regulation, tolerance, stimuli response, and stress or stimuli response. These results are shown in the Supplementary file 2, where the highlighted colours blue and olive green, respectively, represent stress-responsive genes that are commonly found in any one of the six-season vs season comparison datasets of *D*. *hirsuta* or *P*. *appendiculatum*. The stress-responsive genes highlighted in yellow mean that both liverworts have these genes in any one of the six comparative datasets (Supplementary file 2). The stress-responsive genes in each liverwort that are particular to the specific season are not highlighted (Supplementary file 2).

The cyclin family protein, the inosine triphosphate pyrophosphatase family protein, the aldolase-type TIM barrel family protein, and the histone acetyltransferase of the CBP family 12 are important examples of stress-responsive proteins that are only upregulated in *D*. *hirsuta* (Supplementary file 2). Members of the cyclin family of proteins, such as cyclins and cyclin-dependent kinases, have long been recognised as contributing to the regulation of stress responses (Kitsios and Doonan, 2011). Inosine triphosphate pyrophosphatase is a protein that works as a molecular defence mechanism in Arabidopsis by activating stress-related genes and preventing premature senescence (Straube et al., 2023) in Arabidopsis. Histone acetyltransferase controls how plants respond to their environment and interacts with genes, whereas glycolate oxidase, which is encoded by the aldolase-type TIM barrel family protein, controls ROS-mediated signal transduction. Therefore, upregulating the genes encoding these proteins in post-monsoon and/ or fruiting season seems rational since they will protect *D*. *hirsuta* from challenges brought on by seasonality (Supplementary file 2).

When compared to *D*. *hirsuta*, *P*. *appendiculatum* solely exhibited upregulation of several stress-responsive genes, suggesting probably a higher level of stress tolerance in *P*. *appendiculatum* (Supplementary file 2). A few of these genes include SNARE-like superfamily protein, Protein kinase superfamily protein, *Arabidopsis thaliana* responsive to abscisic acid 1b, F-type h+-transporting atpase subunit a, Adp-ribosylation factor-like protein 5b, Double-stranded RNA-binding protein 2, and Clathrin heavy chain 1 (Supplementary file 2). These genes exhibited a strong upregulation in the post-monsoon season, suggesting that *P*. *appendiculatum* had already prepared to confront the seasonality-driven challenges of the fruiting season.

Cytochrome c oxidase subunit 1, 50S ribosomal protein L14, TCP-1/cpn60 chaperonin family protein, Molecular chaperone htpg, Methylcrotonoyl-CoA carboxylase beta chain, Tubulin/FtsZ family protein, ARM repeat protein interacting with ABF2, Regulator of nonsense transcripts 1 homolog, and Methylcrotonoyl-CoA carboxylase beta chain proteins appeared as a common stress-responsive proteins of both organisms (Supplementary file 2). The majority of these proteins offer defence against various abiotic stresses (Kim et al., 2004; Nixon et al., 2005; Wagner et al., 2011; Liberatore et al., 2016; Analin et al., 2020). Therefore, the elevated expression of these genes, primarily during the post-monsoon and/or fruiting season, shows many commonalities in the stress response approaches of both liverwort species.

We also identified the stress-responsive genes in both liverworts that are linked to particular seasons. *D*. *hirsuta* was shown to have a higher number of season-specific genes than *P*. *appendiculatum*; this suggests that *D*. *hirsuta* requires more specialised reprogramming during specific seasons (Supplementary file 2). Ten distinct genes, for instance, are included in the GS1 vs GS3 data and are either directly or indirectly linked to reducing stress (Supplementary file 2). The genes encoding the SecY protein transport family protein, putative ABC transport system permease protein, and photosystem I reaction centre subunit xi, however, appear to be the most significant of these ten genes. SecY protein transport family protein and putative ABC transport system permease protein are known to change membrane transport rates in response to abiotic stressors, which could be the cause of their upregulation (Ma and Browse, 2006; Kang et al, 2011). Since PSII activity declines in cold situations, the photosystem I reaction centre subunit xi upregulation shows that *D. hirsuta* has an acclimatory response in which it tends to maintain its ATP demand in cold conditions. Six distinct genes, namely O-fucosyltransferase family protein, Ribosomal protein S19 family protein, Ribosomal protein S8e family protein, Translation elongation factor efg/ef2 protein, U5 small nuclear ribonucleoprotein helicase, and *Arabidopsis thaliana* exportin 1are present in the GS1 vs GS4 dataset, since they are mostly involved in stress responses, their substantial change is justified (Supplementary file 2). Approaching GS2 vs GS3 revealed the notable alterations in fourteen distinct genes (Supplementary file 2). Glutamine synthetase cytosolic isozyme 1-4, chlorophyll a-b binding protein CP29.1, and lactate/malate dehydrogenase family protein appear to be the main contributors to *D. hirsuta* adaptation during the change from the monsoon to the post-monsoon season. Cytosolic isozyme 1-4 of glutamine synthetase is very advantageous in the assimilation of ammonia in amino acids and provides protection against various abiotic stimuli by preserving cellular homeostasis and this helps to explain the upregulation of this enzyme in post-monsoon season (Jain et al., 2020). Thus, increased expression of it will support thallus development, maintenance, and maturity. Additionally, an increase in the gene expression of the protein CP29.1, which binds chlorophyll a–b, suggests an adaptive response to sustain photosynthesis during post-monsoon seasons when light durations and temperature declines. The greatest number of distinct genes (24) was identified in GS3 vs GS4 dataset (Supplementary file 2). Six out of 24 genes exhibited notable upregulation and the remaining 18 showed downregulation. Given the abundance of down-regulating genes, it is possible that *D*. *hirsuta* tends to preserve energy by only upregulating genes that are necessary to defend it from the harsh conditions of the fruiting season and downregulating genes that are not highly essential.

The number of season-specific unique genes in *P. appendiculatum* was significantly lower than in *D*. *hirsuta* (Supplementary file 2). For example, no unique genes were found in GS2 vs GS3 and GS2 vs GS4, whereas 5, 3, 10, and 2 unique genes were found in the GS1 vs GS2, GS1 vs GS3, GS1 vs GS4, and GS3 vs GS4 datasets, respectively. Significant changes in unique stress-responsive genes were observed during the transition from pre-monsoon to fruiting season, suggesting a strong reprogramming of these genes to enable *P. appendiculatum* to cope with stress, which are particularly prevalent during fruiting season. Eukaryotic translation initiation factor 4b1, nuclear-encoded mitochondrial ribosomal protein S14, transcription factor-like protein DPB, survival protein SurE-like phosphatase/nucleotidase, and galactose oxidase/kelch repeat superfamily protein are among the significant genes included in the aforementioned datasets. These genes are all linked to defence against biotic and abiotic stressors, therefore it makes sense for them to be upregulated.

## 5 CONCLUSIONS

In this study, we investigated the transcriptomes of two liverworts, *D*. *hirsuta* (which evolved approximately 145 million years ago) and *P*. *appendiculatum* (which evolved approximately 25 million years ago), that coexist in the same environment today, during each of their four distinct growing seasons (pre-monsoon, monsoon, post-monsoon, and fruiting season). For both liverworts, the fruiting season and the post-monsoon season are the most stressful of these four seasons because of factors like decreased temperature, precipitation, availability of nutrients, and day length. The RNA-seq analysis showed that both liverwort species, with a few significant differences largely use the same seasonal adaptive mechanisms by altering the expression of genes mainly involved in the same functions. *D*. *hirsuta* primarily modifies its transcriptome in the post-monsoon season by differentially regulating growth, metabolism, and stress-responsive genes and related TFs, most likely to develop reproductive organs, to cope with the harsh environmental conditions there and to get ready for the even harsher fruiting season. During fruiting seasons, it primarily induces stress-responsive genes specific to that season by downregulating other genes, which is likely a strategic way to deal with this season by conserving energy. *P*. *appendiculatum*, on the other hand, showed major transcriptome variability in both the fruiting and post-monsoon seasons by differentially regulating essentially same growth, metabolic, and stress-responsive genes and TFs as *D*. *hirsuta*, but to a lesser extent. It suggests from this that *P*. *appendiculatum* has been strategically modulating its necessary gene expression levels over an extended period of time while considering energy conservation to survive the harsh conditions of both seasons. Like *D*. *hirsuta*, it also induces the greatest number of unique genes during the fruiting season in order to withstand demanding environments. To the best of our knowledge, this is the first work to examine the seasonal transcriptome of early land plants, such as liverworts, and it offers valuable information about how two different species of liverworts that coexist in the same habitat but have distinct evolutionary divergence times endure seasonal changes. These findings are also shown in a hypothetical model for the ease of the readers (Fig. 8).

**Figure 8.**
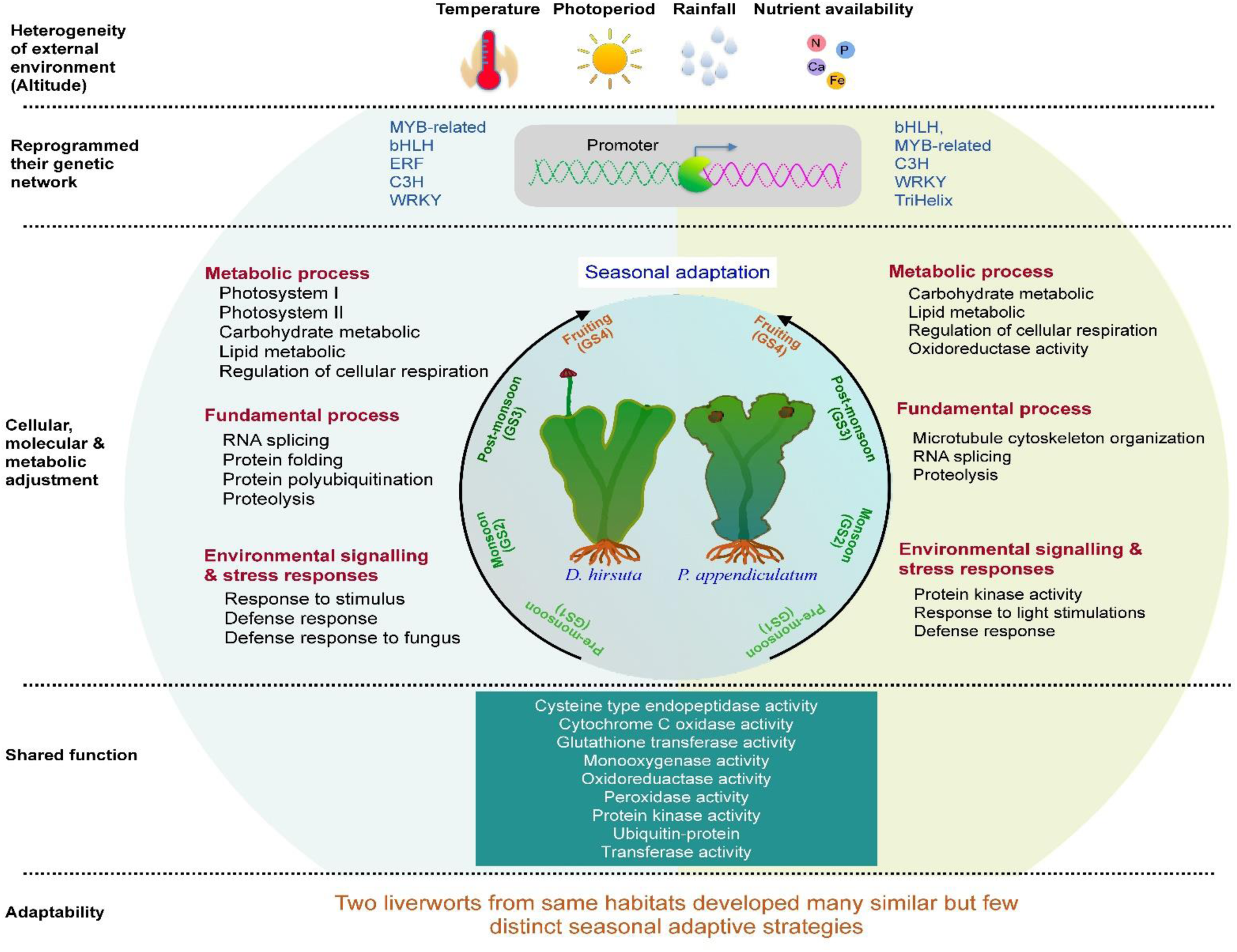
A hypothetical model based on RNA-Seq data illustrates how two liverworts, *D*. *hirsuta* and *P*. *appendiculatum*, which diverged at different times but coexisted in the same habitat, modulate their metabolic, cellular, and molecular processes to adapt to seasonal changes.

## DATA AVAILABILITY STATEMENT

The dataset generated during the current study have been deposited in NCBI’s BioProject Accession Number PRJNA1117754 (http://www.ncbi.nlm.nih.gov/bioproject/1117754).

## DECLARATION OF COMPETING INTEREST

The authors declare that they have no known competing financial interests or personal relationships that could have appeared to influence the work reported in this paper.

## ACKNOWLEDGMENTS

Dr. Sandhya Yadav is thankful to University Grant Commission (UGC), New Delhi, India, for the Senior Research Fellowship (SRF). Dr. Akanksha Srivastava is thankful to the DST– INSPIRE Fellowship, New Delhi, India. Mr. Subhankar Biswas is thankful to Council of Scientific and Industrial Research (CSIR), New Delhi, India for SRF. Vishal Kumar Jha and Kritika Tripathi is thankful to UGC for JRF. We appreciate that Nucleome Informatics Pvt Ltd, Hyderabad, provided the raw data and performed the RNA sequencing work. We extend our sincere gratitude to Mr. Naveen Kumar Pandey of Novelgene Technologies Pvt Ltd, Hyderabad, for his invaluable assistance with data analysis and manuscript preparation for publication. Dr. Yogesh Mishra acknowledges the financial support provided by the Department of Biotechnology (DBT), Government of India (Grant number: BT/PR32376/AGIII/103/1145/2019) and Institute of Eminence (IOE) Incentive Grant, Banaras Hindu University, Varanasi, India (Grant number: R/Dev/D/IOE/Incentive/2021-22/32401). We are highly thankful to Dr. N. Odyuo, Scientist E & H.o.O, BSI-Shillong for the help extended during the collection of bryophytes. Further, we also acknowledge Mr. David W Lamare for the assistance during sample collections.

## AUTHORS’ CONTRIBUTIONS

YM, NC, and SKS conceived the idea and designed the experiments. S Basu, SY, SB, and VKJ conducted the experiments. S Basu, SY, SB, VKJ, AS, KT, and RM analyzed the data. YM and AS wrote the manuscript with help from the other co-authors.

## Notes

### Competing Interest Statement

The authors have declared no competing interest.

