## Supplementary material for "Two liverworts from same habitats developed many similar but few distinct seasonal adaptive strategies: Insights from a transcriptomic approach": Suplementary file 1

**Table S1** List of primers used to validate RNA-Seq data of *D. hirsuta* using RT-PCR

| **Transcript name/id** | **length** | **%GC** | **Tm (°C)** | **Sequence** |
|---|---|---|---|---|
| Actin_FP (Reference gene) | 20 | 50 | 58 | TCACCCACGACTACCACAAA |
| Actin_RP (Reference gene) | 20 | 50 | 58 | CTCCAGCGTCTTCTCCTGTC |
| GAPDH_FP (Reference gene) | 20 | 50 | 53 | CGCGTCTGGGTTTAATATGC |
| GAPDH_RP (Reference gene) | 20 | 50 | 55.4 | TCGAATACGTGGTTGAGTCG |
| TRINITY_DN1638_FP | 21 | 50 | 58 | CCAGGGAATTCTCGTGGTAA |
| TRINITY_DN1638_RP | 21 | 50 | 58 | CCAGAAAAGTCGAAGCGAAC |
| TRINITY_DN256219_FP | 20 | 50 | 56.5 | CCTCTTGGCCACCCATACTA |
| TRINITY_DN256219_RP | 20 | 50 | 55.4 | ATGTGTGCAACGTCTGCTTC |
| TRINITY_DN200_FP | 20 | 50 | 57.3 | CACTGGCGCTAGCATATTCA |
| TRINITY_DN200_RP | 20 | 55 | 59.35 | CCTCCAAGAGCAGGATCAAG |
| TRINITY_DN27350 FP | 20 | 50 | 60.1 | ACCACTATCGCAGGATTTCG |
| TRINITY_DN27350 RP | 20 | 50 | 60.9 | TCACTGCAAGCAATGGTGTC |
| TRINITY_DN26369 FP | 20 | 50 | 60.3 | AACACTTCCAAGCTCCAACG |
| TRINITY_DN26369 RP | 20 | 50 | 60.9 | ATGCAGATATCCGTCCATCG |
| TRINITY_DN975_FP | 20 | 50 | 60.2 | CTGGGTTTGCTTGTGTGATG |
| TRINITY_DN975_RP | 20 | 50 | 60.5 | TGTGGCCGAAGAAGAAGAAG |
| TRINITY_DN413 FP | 20 | 50 | 57.3 | TCTGGTGGCTGTTTGTTGAC |
| TRINITY_DN413 RP | 20 | 50 | 57.3 | GCCATGGTTGCTGTTGTATG |
| TRINITY_DN527_FP | 20 | 45 | 55.25 | TGCAACAACCCATACCTGAA |
| TRINITY_DN527_RP | 20 | 50 | 57.3 | TTCCCTATCGACGTTCCTTG |
| TRINITY_DN8767_FP | 20 | 50 | 57.3 | CGATTCGAGCCTTATCGAAG |
| TRINITY_DN8767_RP | 20 | 50 | 57.3 | CCGGAGTACCAAATTGCTGT |
| TRINITY_DN2320_FP | 20 | 50 | 56.3 | CCTTTCACCGACCAAAGTGT |
| TRINITY_DN2320_FP | 20 | 50 | 54.8 | GTGCTTTTCGGAACATCCAT |

**Table S2** List of primers used to validate RNA-Seq data of *P. appendiculatum* using RT-PCR

| **Transcript name/id** | **length** | **%GC** | **Tm (℃)** | **Sequence** |
|---|---|---|---|---|
| TRINITY_DN1365_Beta tubulin_FP (Reference gene) | 20 | 50 | 60.1 | AGATGTGGGACGCAAAGAAC |
| TRINITY_DN1365_Beta tubulin_RP (Reference gene) | 20 | 50 | 60 | GATCATCTGCTCGTCGACTTC |
| TRINITY_DN82941_Histone H3_FP (Reference gene) | 22 | 50 | 59.6 | AAGAAGGCCCACAGATACAGAC |
| TRINITY_DN82941_Histone H3_RP (Reference gene) | 20 | 50 | 60.6 | GGCGATGTTCTTGACGATTC |
| TRINITY_DN12862 FP | 20 | 50 | 60.5 | CGAAGGGTTCACAGAAATGG |
| TRINITY_DN12862 RP | 20 | 50 | 59.8 | CCGAATGGCAAACTTCTCTC |
| TRINITY_DN835_FP | 20 | 50 | 57.8 | CATGAATTGTCCGTGGAGAG |
| TRINITY_DN835_RP | 20 | 50 | 59.2 | TGGAGACCTGAACCCAAATC |
| TRINITY_DN18949_FP | 20 | 50 | 60.1 | AGCAAACCACCAGAATGGAG |
| TRINITY_DN18949_RP | 20 | 50 | 60 | TCAGTCCTTGTGCGTTGAAG |
| TRINITY_DN14333_FP | 20 | 50 | 61.1 | CAGGATGTTTGTCCCCTTTC |
| TRINITY_DN14333_RP | 20 | 50 | 59 | TGGTTCATTGCCAGGTGTAG |
| TRINITY_DN204553_FP | 20 | 50 | 58.3 | GAACCCAACATTGTCACCAG |
| TRINITY_DN204553_RP | 20 | 50 | 59 | TCAAGCCTGGTATGGTTGTG |
| TRINITY_DN3740_FP | 20 | 50 | 59.3 | GTGTTTCCGATGCAGCTATG |
| TRINITY_DN3740_RP | 20 | 50 | 59 | ACCGAGGAACTTCCAACATC |
| TRINITY_DN3958_FP | 20 | 50 | 58.6 | AAAGGAGCAGCGAGAGATTG |
| TRINITY_DN3958_RP | 20 | 50 | 58.3 | GAACTCGGCTTCAAATCTGG |
| TRINITY_DN13268_FP | 20 | 50 | 59.7 | CGTACTCATGCAAGGAATGC |
| TRINITY_DN13268_RP | 20 | 50 | 59 | ATTCTGACTGAGTGTGCGTTG |
| TRINITY_DN5330_FP | 20 | 50 | 58.2 | TCGAGCTTCTGTTGATGCAC |
| TRINITY_DN5330_RP | 20 | 50 | 57.3 | GTCATGGCGATGTTCAACTG |
| TRINITY_DN14232_FP | 20 | 50 | 60.4 | AAGCAACAGAAGCACCAAGG |
| TRINITY_DN14232_RP | 20 | 50 | 60 | TTAATGGGCTGGGCAGTTAC |

**Table S3** DEG analysis summary of *D. hirsuta* and *P. appendiculatum*

| Comparison | Condition 1 (Group name) | Condition 2 (Group name) | Up-Regulation (genes) | Down-Regulation (genes) |
|---|---|---|---|---|
| 1 | DUGS1 | DUGS2 | 1531 | 1978 |
| 2 | DUGS1 | DUGS3 | 4110 | 3138 |
| 3 | DUGS1 | DUGS4 | 7721 | 4778 |
| 4 | DUGS2 | DUGS3 | 6185 | 4192 |
| 5 | DUGS3 | DUGS4 | 10638 | 8653 |
| 6 | DUGS2 | DUGS4 | 7469 | 4783 |
| 7 | PCGS1 | PCGS2 | 9047 | 10649 |
| 8 | PCGS1 | PCGS3 | 33805 | 36479 |
| 9 | PCGS1 | PCGS4 | 43617 | 30617 |
| 10 | PCGS2 | PCGS3 | 29561 | 22583 |
| 11 | PCGS3 | PCGS4 | 18870 | 29081 |
| 12 | PCGS2 | PCGS4 | 42589 | 21590 |

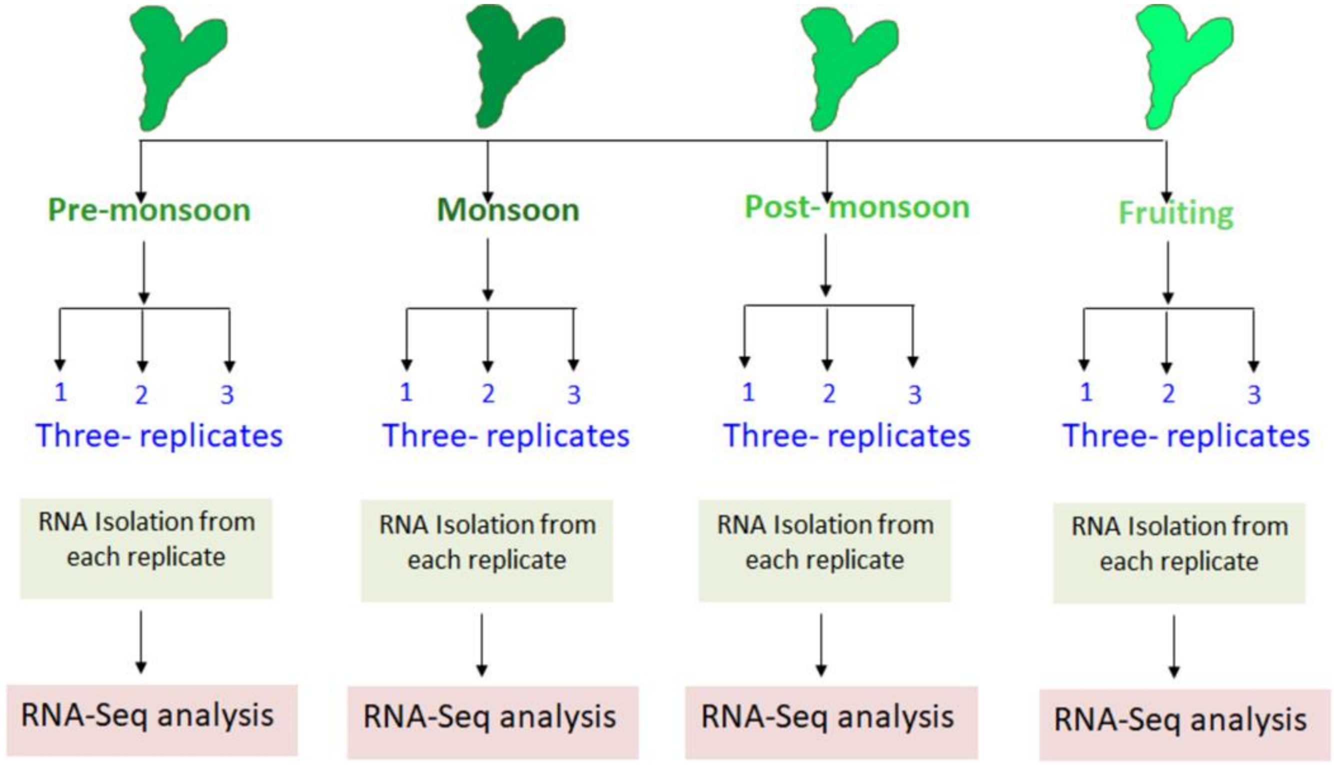


**Figure S1** Schematic representation of experimental setup used to perform the RNA-seq analysis of D. *hirsuta* and *P. appendiculatum* during four contrasting growing seasons of year 2019-2020.

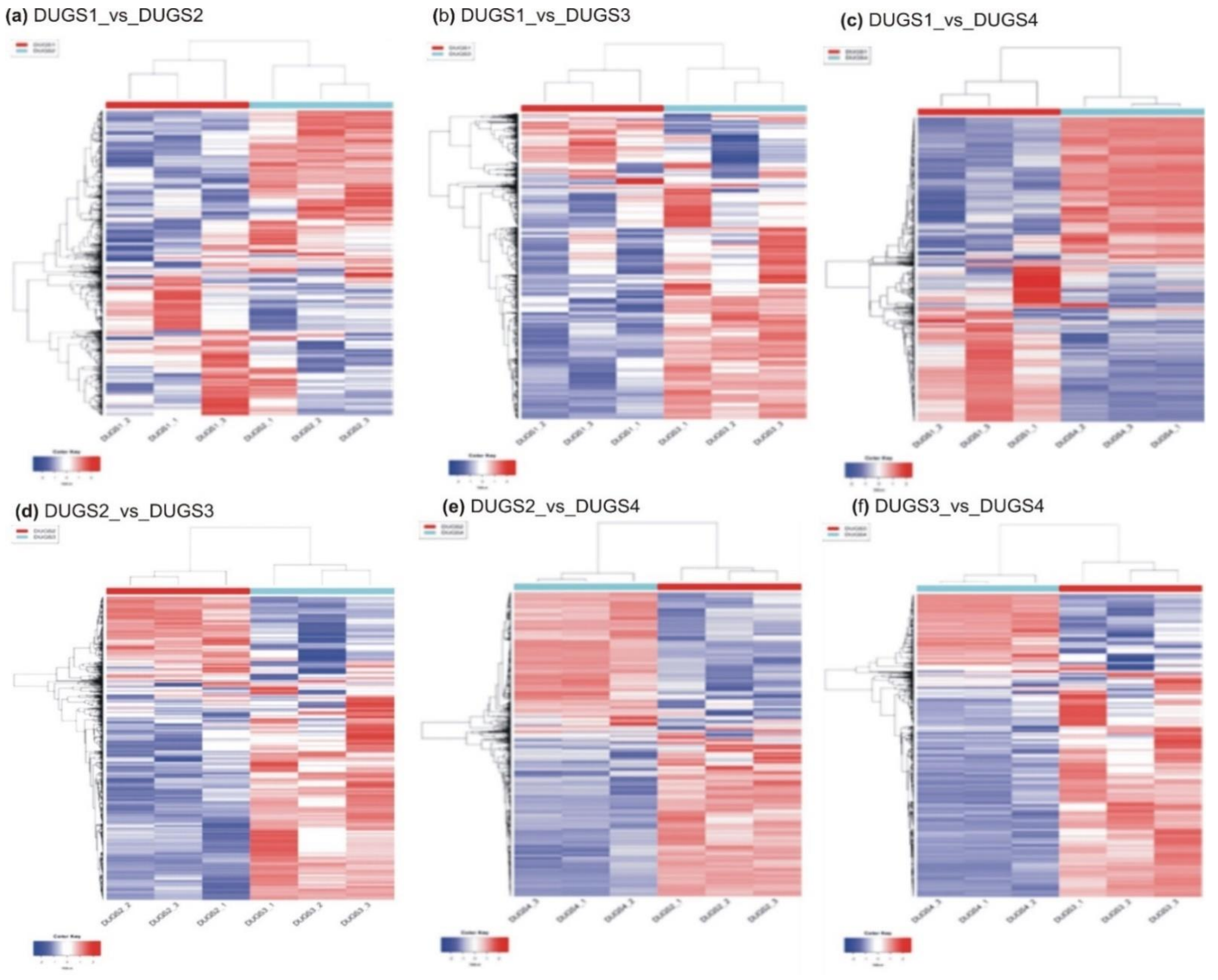


**Figure S2** Heatmaps with hierarchical clustering of all differentially expressed transcripts identified in each of the six paired-wise comparisons of all four seasons (DUGS1_vs _DUGS2, DUGS1_vs_DUGS3, DUGS1 _vs _DUGS4, DUGS2_vs _DUGS3, DUGS2_vs _DUGS4 and DUGS3_vs_DUGS4) of *D. hirsuta* in three biological replicates.

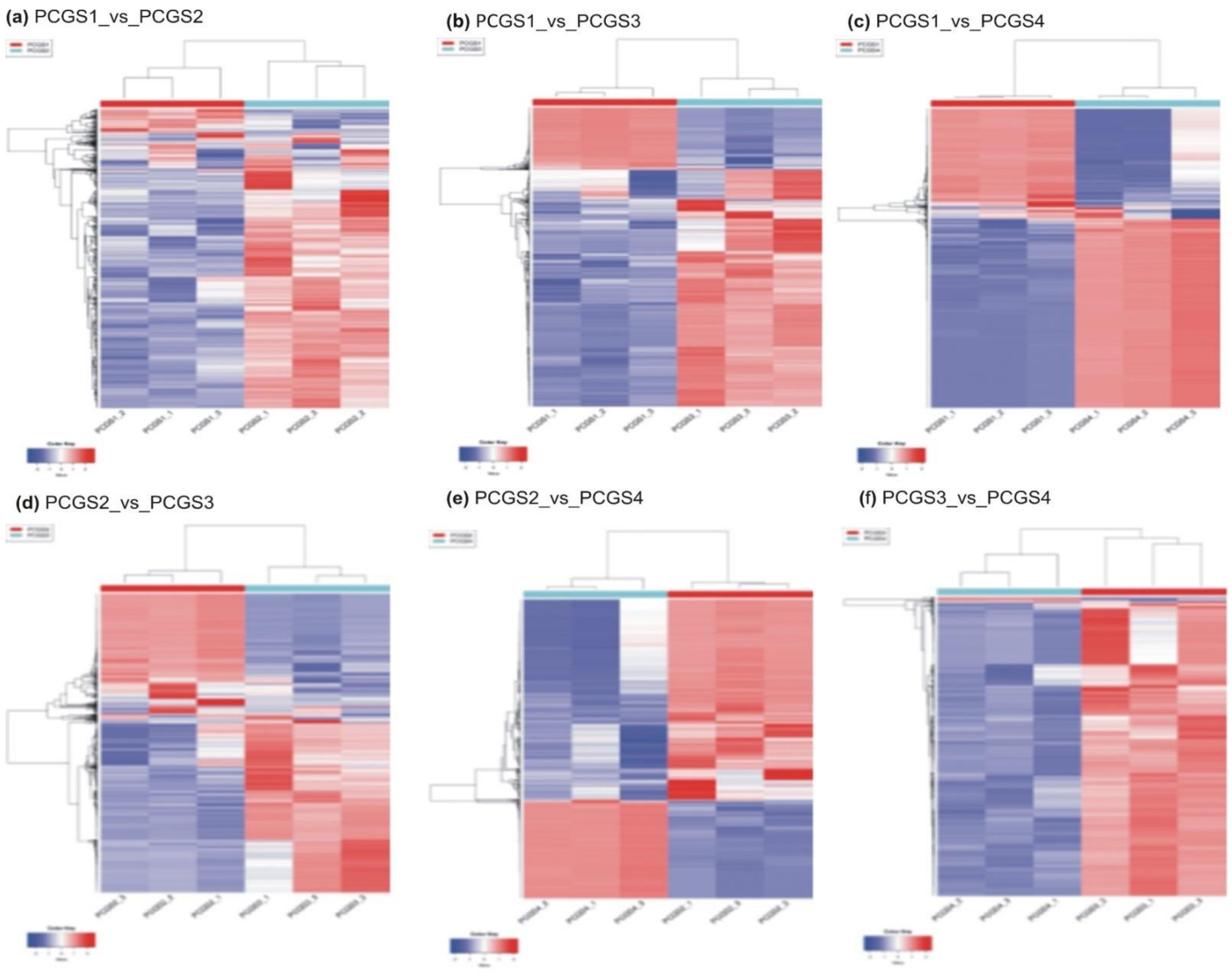


**Figure S3** Heatmaps with hierarchical clustering of all differentially expressed transcripts identified in each of the six paired-wise comparisons of all four seasons (DUGS1_vs _DUGS2, DUGS1_vs_DUGS3, DUGS1 _vs _DUGS4, DUGS2_vs _DUGS3, DUGS2_vs _DUGS4 and DUGS3_vs_DUGS4) of *P. appendiculatum* in three biological replicates.

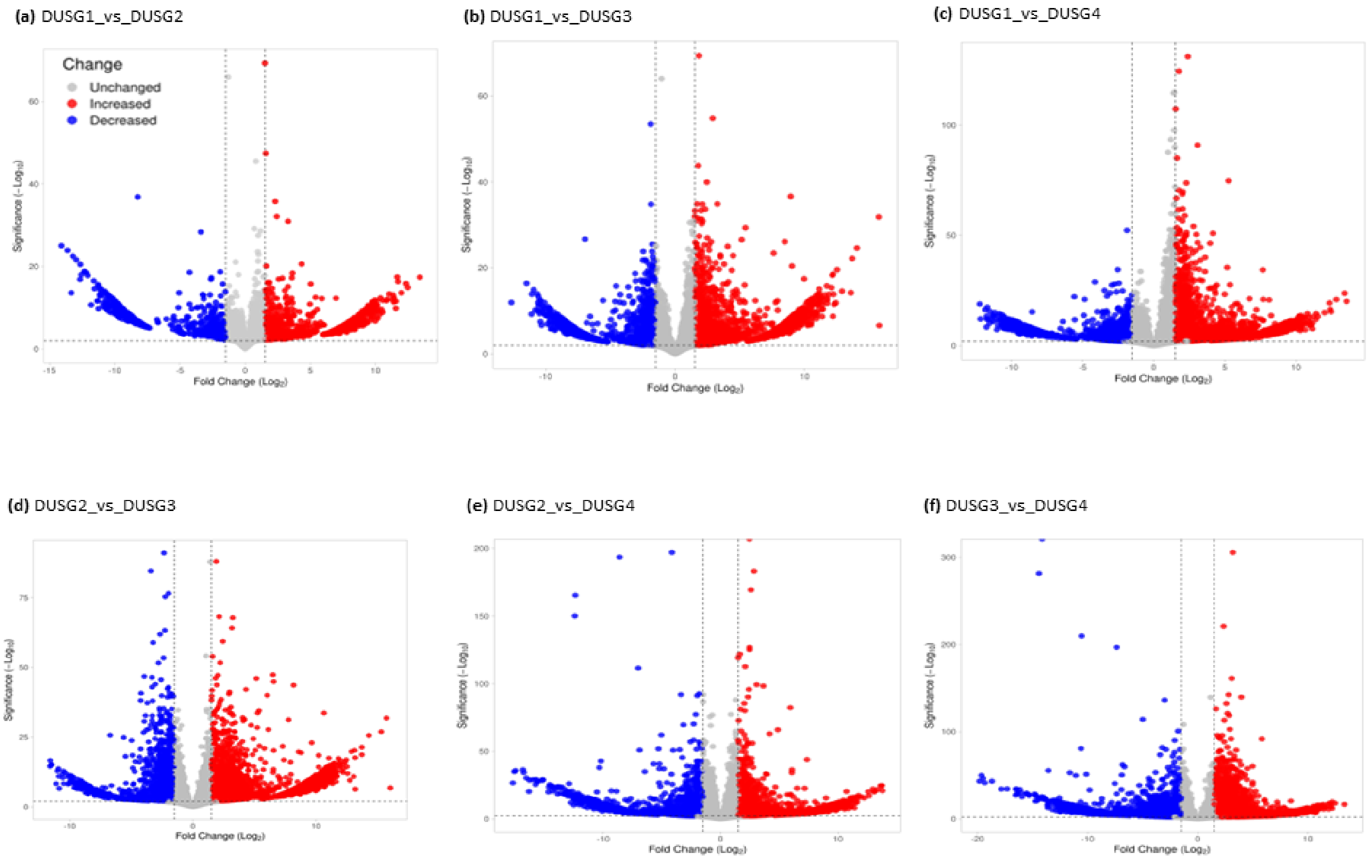


**Figure S4** Volcano plot depicting all differentially expressed transcripts identified in each of the six paired-wise comparisons of all four seasons (DUGS1_vs _DUGS2, DUGS1_vs_DUGS3, DUGS1 _vs _DUGS4, DUGS2_vs _DUGS3, DUGS2_vs _DUGS4 and DUGS3_vs_DUGS4) of *D. hirsuta*.

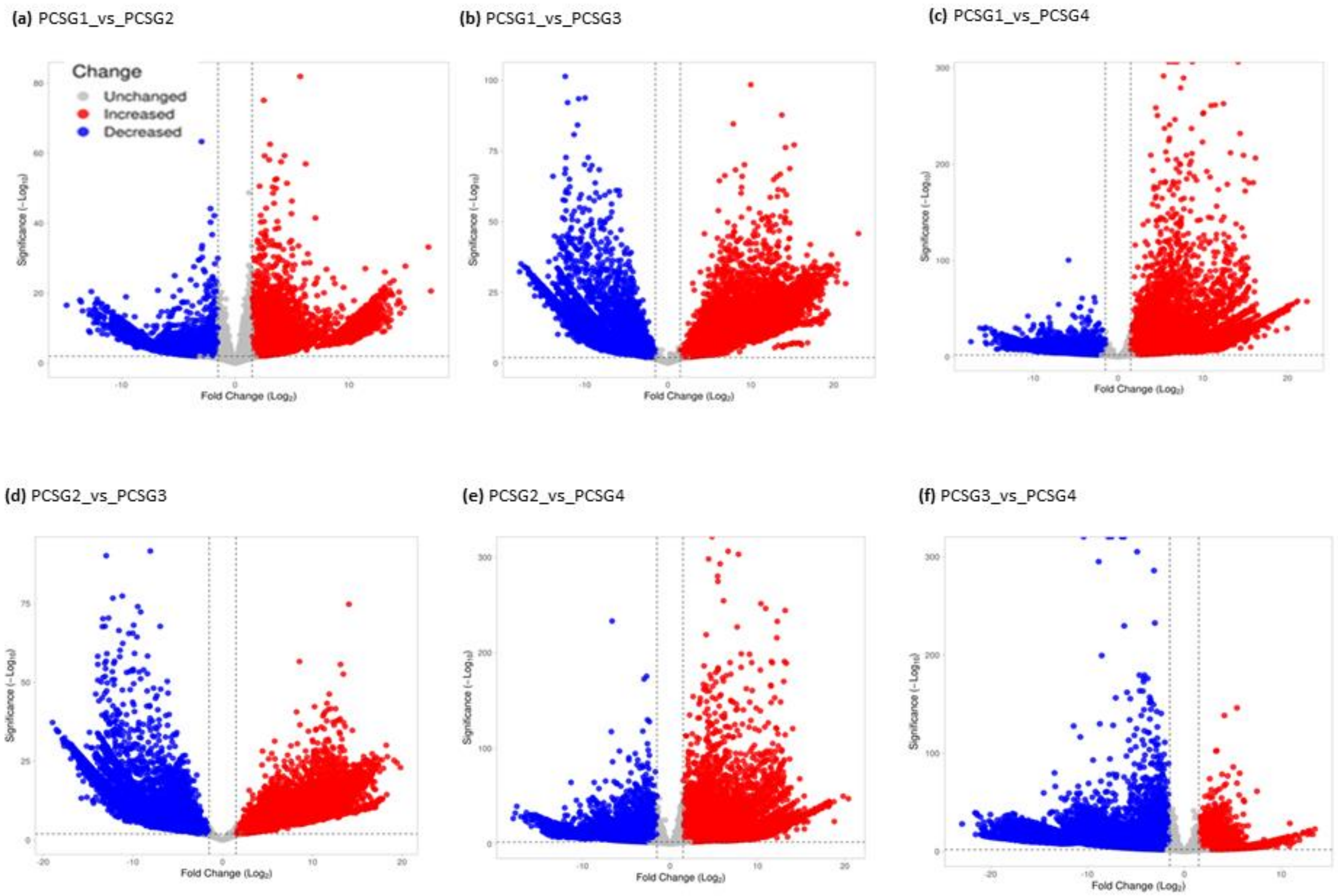


**Figure S5** Volcano plot depicting all differentially expressed transcripts identified in each of the six paired-wise comparisons of all four seasons (DUGS1_vs _DUGS2, DUGS1_vs_DUGS3, DUGS1 _vs _DUGS4, DUGS2_vs _DUGS3, DUGS2_vs _DUGS4 and DUGS3_vs_DUGS4) of *P. appendiculatum*.

## (a) BP

| Conditions | UP | GO term | Down |
|---|---|---|---|
| DUGS1_vs_2 | 0 | Carbohydrate metabolic process | 9 |
| DUGS1_vs_3 | 0 | | 13 |
| DUGS1_vs_4 | 0 | | 14 |
| DUGS2_vs_3 | 0 | | 0 |
| DUGS2_vs_4 | 14 | | 11 |
| DUGS3_vs_4 | 29 | | 0 |
| DUGS1_vs_2 | 0 | Cellular process | 0 |
| DUGS1_vs_3 | 0 | | 20 |
| DUGS1_vs_4 | 0 | | 0 |
| DUGS2_vs_3 | 0 | | 14 |
| DUGS2_vs_4 | 16 | | 11 |
| DUGS3_vs_4 | 0 | | 0 |
| DUGS1_vs_2 | 5 | Cytoplasmic translation | 0 |
| DUGS1_vs_3 | 43 | | 28 |
| DUGS1_vs_4 | 19 | | 0 |
| DUGS2_vs_3 | 63 | | 15 |
| DUGS2_vs_4 | 0 | | 26 |
| DUGS3_vs_4 | 0 | | 123 |
| DUGS1_vs_2 | 0 | Cytoplasmic translational elongation | 9 |
| DUGS1_vs_3 | 0 | | 0 |
| DUGS1_vs_4 | 0 | | 15 |
| DUGS2_vs_3 | 12 | | 0 |
| DUGS2_vs_4 | 0 | | 14 |
| DUGS3_vs_4 | 0 | | 19 |
| DUGS1_vs_2 | 0 | Defense response | 0 |
| DUGS1_vs_3 | 13 | | 13 |
| DUGS1_vs_4 | 37 | | 13 |
| DUGS2_vs_3 | 15 | | 15 |
| DUGS2_vs_4 | 41 | | 19 |
| DUGS3_vs_4 | 35 | | 0 |
| DUGS1_vs_2 | 6 | Defense response to fungus | 0 |
| DUGS1_vs_3 | 0 | | 12 |
| DUGS1_vs_4 | 0 | | 0 |
| DUGS2_vs_3 | 0 | | 28 |
| DUGS2_vs_4 | 0 | | 30 |
| DUGS3_vs_4 | 0 | | 0 |
| DUGS1_vs_2 | 4 | Fatty acid metabolic process | 0 |
| DUGS1_vs_3 | 0 | | 0 |
| DUGS1_vs_4 | 0 | | 0 |
| DUGS2_vs_3 | 0 | | 0 |
| DUGS2_vs_4 | 0 | | 0 |
| DUGS3_vs_4 | 0 | | 0 |
| DUGS1_vs_2 | 0 | Lipid metabolic process | 7 |
| DUGS1_vs_3 | 0 | | 0 |
| DUGS1_vs_4 | 17 | | 53 |
| DUGS2_vs_3 | 0 | | 0 |
| DUGS2_vs_4 | 16 | | 55 |
| DUGS3_vs_4 | 0 | | 0 |
| DUGS1_vs_2 | 0 | Maturation of LSU-rRNA from tricistronic rRNA transcript (SSU-rRNA, 5.8S rRNA, LSU-rRNA) | 0 |
| DUGS1_vs_3 | 10 | | 0 |
| DUGS1_vs_4 | 0 | | 9 |
| DUGS2_vs_3 | 26 | | 15 |
| DUGS2_vs_4 | 0 | | 77 |
| DUGS3_vs_4 | 0 | | 0 |
| DUGS1_vs_2 | 6 | Microtubule cytoskeleton organization | 0 |
| DUGS1_vs_3 | 11 | | 18 |
| DUGS1_vs_4 | 24 | | 10 |
| DUGS2_vs_3 | 22 | | 21 |
| DUGS2_vs_4 | 0 | | 17 |
| DUGS3_vs_4 | 38 | | 0 |

| Conditions | UP | GO term | Down |
|---|---|---|---|
| DUGS1_vs_2 | 0 | mRNA splicing, via spliceosome | 0 |
| DUGS1_vs_3 | 0 | | 0 |
| DUGS1_vs_4 | 27 | | 10 |
| DUGS2_vs_3 | 0 | | 0 |
| DUGS2_vs_4 | 23 | | 10 |
| DUGS3_vs_4 | 40 | | 0 |
| DUGS1_vs_2 | 0 | Nuclear-transcribed mRNA catabolic process, nonsense-mediated decay | 0 |
| DUGS1_vs_3 | 0 | | 0 |
| DUGS1_vs_4 | 0 | | 31 |
| DUGS2_vs_3 | 0 | | 0 |
| DUGS2_vs_4 | 0 | | 65 |
| DUGS3_vs_4 | 24 | | 0 |
| DUGS1_vs_2 | 0 | Protein folding | 0 |
| DUGS1_vs_3 | 26 | | 0 |
| DUGS1_vs_4 | 23 | | 0 |
| DUGS2_vs_3 | 28 | | 0 |
| DUGS2_vs_4 | 22 | | 0 |
| DUGS3_vs_4 | 0 | | 0 |
| DUGS1_vs_2 | 0 | Protein polyubiquitination | 12 |
| DUGS1_vs_3 | 0 | | 0 |
| DUGS1_vs_4 | 20 | | 9 |
| DUGS2_vs_3 | 0 | | 29 |
| DUGS2_vs_4 | 0 | | 0 |
| DUGS3_vs_4 | 28 | | 0 |
| DUGS1_vs_2 | 0 | Proteolysis | 0 |
| DUGS1_vs_3 | 0 | | 0 |
| DUGS1_vs_4 | 0 | | 9 |
| DUGS2_vs_3 | 14 | | 0 |
| DUGS2_vs_4 | 0 | | 11 |
| DUGS3_vs_4 | 0 | | 19 |
| DUGS1_vs_2 | 0 | Regulation of cellular respiration | 7 |
| DUGS1_vs_3 | 11 | | 0 |
| DUGS1_vs_4 | 0 | | 0 |
| DUGS2_vs_3 | 24 | | 13 |
| DUGS2_vs_4 | 32 | | 33 |
| DUGS3_vs_4 | 0 | | 30 |
| DUGS1_vs_2 | 14 | Response to light stimulus | 12 |
| DUGS1_vs_3 | 0 | | 0 |
| DUGS1_vs_4 | 0 | | 0 |
| DUGS2_vs_3 | 0 | | 0 |
| DUGS2_vs_4 | 0 | | 9 |
| DUGS3_vs_4 | 0 | | 26 |
| DUGS1_vs_2 | 9 | Ribosomal large subunit assembly | 5 |
| DUGS1_vs_3 | 16 | | 0 |
| DUGS1_vs_4 | 0 | | 0 |
| DUGS2_vs_3 | 23 | | 0 |
| DUGS2_vs_4 | 0 | | 0 |
| DUGS3_vs_4 | 23 | | 44 |
| DUGS1_vs_2 | 6 | Ribosomal small subunit assembly | 10 |
| DUGS1_vs_3 | 35 | | 12 |
| DUGS1_vs_4 | 17 | | 0 |
| DUGS2 _vs_3 | 31 | | 0 |
| DUGS2_vs_4 | 0 | | 0 |
| DUGS3_vs_4 | 0 | | 0 |
| DUGS1_vs_2 | 0 | rRNA processing | 5 |
| DUGS1_vs_3 | 0 | | 0 |
| DUGS1_vs_4 | 0 | | 0 |
| DUGS2_vs_3 | 13 | | 0 |
| DUGS2_vs_4 | 0 | | 0 |
| DUGS3_vs_4 | 0 | | 0 |

## (b) CC

| Conditions | Up | GO term | Down |
|---|---|---|---|
| DUGS1_vs_2 | 0 | Cellular anatomical entity | 9 |
| DUGS1_vs_3 | 10 | | 0 |
| DUGS1_vs_4 | 0 | | 0 |
| DUGS2_vs_3 | 22 | | 0 |
| DUGS2_vs_4 | 22 | | 0 |
| DUGS3_vs_4 | 0 | | 25 |
| DUGS1_vs_2 | 0 | Chloroplast | 36 |
| DUGS1_vs_3 | 45 | | 19 |
| DUGS1_vs_4 | 0 | | 82 |
| DUGS2_vs_3 | 95 | | 24 |
| DUGS2_vs_4 | 21 | | 33 |
| DUGS3_vs_4 | 0 | | 128 |
| DUGS1_vs_2 | 8 | Chloroplast thylakoid membrane | 4 |
| DUGS1_vs_3 | 0 | | 0 |
| DUGS1_vs_4 | 0 | | 15 |
| DUGS2_vs_3 | 0 | | 0 |
| DUGS2_vs_4 | 0 | | 12 |
| DUGS3_vs_4 | 0 | | 0 |
| DUGS1_vs_2 | 0 | Cytoplasm | 6 |
| DUGS1_vs_3 | 0 | | 41 |
| DUGS1_vs_4 | 0 | | 35 |
| DUGS2_vs_3 | 0 | | 65 |
| DUGS2_vs_4 | 15 | | 51 |
| DUGS3_vs_4 | 0 | | 56 |
| DUGS1_vs_2 | 20 | Cytosol | 0 |
| DUGS1_vs_3 | 29 | | 11 |
| DUGS1_vs_4 | 119 | | 13 |
| DUGS2_vs_3 | 35 | | 21 |
| DUGS2_vs_4 | 105 | | 19 |
| DUGS3_vs_4 | 146 | | 28 |
| DUGS1_vs_2 | 0 | DNA-directed RNA polymerase complex | 4 |
| DUGS1_vs_3 | 16 | | 0 |
| DUGS1_vs_4 | 43 | | 0 |
| DUGS2_vs_3 | 21 | | 0 |
| DUGS2_vs_4 | 15 | | 0 |
| DUGS3_vs_4 | 61 | | 0 |
| DUGS1_vs_2 | 4 | Endoplasmic reticulum | 0 |
| DUGS1_vs_3 | 12 | | 0 |
| DUGS1_vs_4 | 32 | | 0 |
| DUGS2_vs_3 | 0 | | 0 |
| DUGS2_vs_4 | 25 | | 10 |
| DUGS3_vs_4 | 41 | | 17 |
| DUGS1_vs_2 | 0 | Endoplasmic reticulum membrane | 0 |
| DUGS1_vs_3 | 0 | | 0 |
| DUGS1_vs_4 | 23 | | 0 |
| DUGS2_vs_3 | 0 | | 0 |
| DUGS2_vs_4 | 0 | | 12 |
| DUGS3_vs_4 | 0 | | 0 |
| DUGS1_vs_2 | 11 | Extracellular region | 4 |
| DUGS1_vs_3 | 23 | | 10 |
| DUGS1_vs_4 | 35 | | 11 |
| DUGS2_vs_3 | 14 | | 14 |
| DUGS2_vs_4 | 40 | | 19 |
| DUGS3_vs_4 | 50 | | 19 |
| DUGS1_vs_2 | 0 | Golgi apparatus | 0 |
| DUGS1_vs_3 | 0 | | 0 |
| DUGS1_vs_4 | 17 | | 0 |
| DUGS2_vs_3 | 0 | | 14 |
| DUGS2_vs_4 | 0 | | 0 |
| DUGS3_vs_4 | 0 | | 0 |
| DUGS1_vs_2 | 0 | Golgi membrane | 0 |
| DUGS1_vs_3 | 0 | | 12 |
| DUGS1_vs_4 | 36 | | 0 |
| DUGS2_vs_3 | 0 | | 23 |
| DUGS2_vs_4 | 15 | | 0 |
| DUGS3_vs_4 | 44 | | 0 |

| Conditions | Up | GO term | Down |
|---|---|---|---|
| DUGS1_vs_2 | 0 | Intracellular anatomical structure | 4 |
| DUGS1_vs_3 | 0 | | 0 |
| DUGS1_vs_4 | 45 | | 0 |
| DUGS2_vs_3 | 0 | | 0 |
| DUGS2_vs_4 | 30 | | 0 |
| DUGS3_vs_4 | 70 | | 0 |
| DUGS1_vs_2 | 39 | Membrane | 63 |
| DUGS1_vs_3 | 156 | | 113 |
| DUGS1_vs_4 | 287 | | 156 |
| DUGS2_vs_3 | 236 | | 185 |
| DUGS2_vs_4 | 283 | | 161 |
| DUGS3_vs_4 | 412 | | 281 |
| DUGS1_vs_2 | 0 | Mitochondrion | 28 |
| DUGS1_vs_3 | 49 | | 13 |
| DUGS1_vs_4 | 18 | | 63 |
| DUGS2_vs_3 | 88 | | 15 |
| DUGS2_vs_4 | 0 | | 33 |
| DUGS3_vs_4 | 29 | | 132 |
| DUGS1_vs_2 | 0 | Nucleosome | 12 |
| DUGS1_vs_3 | 16 | | 15 |
| DUGS1_vs_4 | 20 | | 42 |
| DUGS2_vs_3 | 23 | | 14 |
| DUGS2_vs_4 | 15 | | 29 |
| DUGS3_vs_4 | 24 | | 47 |
| DUGS1_vs_2 | 11 | Nucleus | 13 |
| DUGS1_vs_3 | 25 | | 42 |
| DUGS1_vs_4 | 126 | | 49 |
| DUGS2_vs_3 | 35 | | 58 |
| DUGS2_vs_4 | 87 | | 53 |
| DUGS3_vs_4 | 182 | | 73 |
| DUGS1_vs_2 | 5 | Photosystem I | 3 |
| DUGS1_vs_3 | 14 | | 0 |
| DUGS1_vs_4 | 58 | | 0 |
| DUGS2_vs_3 | 18 | | 0 |
| DUGS2_vs_4 | 48 | | 0 |
| DUGS3_vs_4 | 73 | | 0 |
| DUGS1_vs_2 | 0 | Photosystem II | 5 |
| DUGS1_vs_3 | 27 | | 0 |
| DUGS1_vs_4 | 0 | | 0 |
| DUGS2_vs_3 | 43 | | 0 |
| DUGS2_vs_4 | 0 | | 0 |
| DUGS3_vs_4 | 0 | | 0 |
| DUGS1_vs_2 | 0 | Plant-type vacuole | 0 |
| DUGS1_vs_3 | 11 | | 8 |
| DUGS1_vs_4 | 0 | | 0 |
| DUGS2_vs_3 | 16 | | 13 |
| DUGS2_vs_4 | 0 | | 0 |
| DUGS3_vs_4 | 0 | | 0 |
| DUGS1_vs_2 | 5 | Plasma membrane | 4 |
| DUGS1_vs_3 | 0 | | 29 |
| DUGS1_vs_4 | 0 | | 26 |
| DUGS2_vs_3 | 0 | | 35 |
| DUGS2_vs_4 | 0 | | 20 |
| DUGS3_vs_4 | 0 | | 19 |
| DUGS1_vs_2 | 0 | Plastid | 18 |
| DUGS1_vs_3 | 27 | | 0 |
| DUGS1_vs_4 | 0 | | 29 |
| DUGS2_vs_3 | 43 | | 0 |
| DUGS2_vs_4 | 0 | | 0 |
| DUGS3_vs_4 | 0 | | 31 |
| DUGS1_vs_2 | 0 | Ribosome | 7 |
| DUGS1_vs_3 | 11 | | 0 |
| DUGS1_vs_4 | 0 | | 13 |
| DUGS2_vs_3 | 16 | | 0 |
| DUGS2_vs_4 | 0 | | 0 |
| DUGS3_vs_4 | 0 | | 69 |

**(c) MF**

| Conditions | Up | GO terms | Down |
|---|---|---|---|
| DUGS1_vs_2 | 7 | 3-deoxy-7-phosphoheptulonate synthase activity | 0 |
| DUGS1_vs_3 | 0 | | 82 |
| DUGS1_vs_4 | 0 | | 11 |
| DUGS2_vs_3 | 0 | | 19 |
| DUGS2_vs_4 | 0 | | 22 |
| DUGS3_vs_4 | 0 | | 0 |
| DUGS1_vs_2 | 45 | ATP binding | 32 |
| DUGS1_vs_3 | 136 | | 0 |
| DUGS1_vs_4 | 324 | | 22 |
| DUGS2_vs_3 | 142 | | 21 |
| DUGS2_vs_4 | 260 | | 15 |
| DUGS3_vs_4 | 0 | | 206 |
| DUGS1_vs_2 | 6 | Calcium ion binding | 0 |
| DUGS1_vs_3 | 24 | | 15 |
| DUGS1_vs_4 | 38 | | 14 |
| DUGS2_vs_3 | 41 | | 0 |
| DUGS2_vs_4 | 26 | | 0 |
| DUGS3_vs_4 | 48 | | 39 |
| DUGS1_vs_2 | 5 | Catalytic activity | 5 |
| DUGS1_vs_3 | 9 | | 14 |
| DUGS1_vs_4 | 26 | | 0 |
| DUGS2_vs_3 | 18 | | 0 |
| DUGS2_vs_4 | 40 | | 0 |
| DUGS3_vs_4 | 41 | | 47 |
| DUGS1_vs_2 | 4 | Catechol oxidase activity | 6 |
| DUGS1_vs_3 | 0 | | 0 |
| DUGS1_vs_4 | 0 | | 12 |
| DUGS2_vs_3 | 0 | | 0 |
| DUGS2_vs_4 | 0 | | 44 |
| DUGS3_vs_4 | 0 | | 16 |
| DUGS1_vs_2 | 5 | Copper ion binding | 0 |
| DUGS1_vs_3 | 0 | | 0 |
| DUGS1_vs_4 | 0 | | 12 |
| DUGS2_vs_3 | 0 | | 81 |
| DUGS2_vs_4 | 0 | | 16 |
| DUGS3_vs_4 | 0 | | 0 |
| DUGS1_vs_2 | 0 | Cysteine-type endopeptidase activity | 6 |
| DUGS1_vs_3 | 0 | | 0 |
| DUGS1_vs_4 | 0 | | 43 |
| DUGS2_vs_3 | 0 | | 29 |
| DUGS2_vs_4 | 0 | | 0 |
| DUGS3_vs_4 | 30 | | 0 |
| DUGS1_vs_2 | 0 | Cytochrome-c oxidase activity | 6 |
| DUGS1_vs_3 | 18 | | 61 |
| DUGS1_vs_4 | 0 | | 12 |
| DUGS2_vs_3 | 29 | | 0 |
| DUGS2_vs_4 | 0 | | 0 |
| DUGS3_vs_4 | 0 | | 17 |
| DUGS1_vs_2 | 10 | DNA binding | 3 |
| DUGS1_vs_3 | 21 | | 17 |
| DUGS1_vs_4 | 100 | | 0 |
| DUGS2_vs_3 | 27 | | 0 |
| DUGS2_vs_4 | 76 | | 0 |
| DUGS3_vs_4 | 185 | | 0 |
| DUGS1_vs_2 | 0 | DNA-binding transcription factor activity | 0 |
| DUGS1_vs_3 | 17 | | 0 |
| DUGS1_vs_4 | 31 | | 0 |
| DUGS2_vs_3 | 0 | | 0 |
| DUGS2_vs_4 | 20 | | 0 |
| DUGS3_vs_4 | 40 | | 0 |
| DUGS1_vs_2 | 0 | Glutathione transferase activity | 4 |
| DUGS1_vs_3 | 14 | | 8 |
| DUGS1_vs_4 | 0 | | 0 |
| DUGS2_vs_3 | 14 | | 14 |
| DUGS2_vs_4 | 0 | | 21 |
| DUGS3_vs_4 | 0 | | 27 |
| DUGS1_vs_2 | 4 | Glycine hydroxymethyltransferase activity | 0 |
| DUGS1_vs_3 | 0 | | 0 |
| DUGS1_vs_4 | 0 | | 0 |
| DUGS2_vs_3 | 0 | | 18 |
| DUGS2_vs_4 | 0 | | 11 |
| DUGS3_vs_4 | 0 | | 0 |

| Conditions | Up | GO terms | Down |
|---|---|---|---|
| DUGS1_vs_2 | 0 | GTPase activator activity | 3 |
| DUGS1_vs_3 | 0 | | 11 |
| DUGS1_vs_4 | 22 | | 14 |
| DUGS2_vs_3 | 14 | | 0 |
| DUGS2_vs_4 | 13 | | 9 |
| DUGS3_vs_4 | 33 | | 0 |
| DUGS1_vs_2 | 4 | GTPase activity | 0 |
| DUGS1_vs_3 | 21 | | 0 |
| DUGS1_vs_4 | 72 | | 15 |
| DUGS2_vs_3 | 0 | | 0 |
| DUGS2_vs_4 | 48 | | 0 |
| DUGS3_vs_4 | 82 | | 16 |
| DUGS1_vs_2 | 0 | Guanyl-nucleotide exchange factor activity | 4 |
| DUGS1_vs_3 | 0 | | 0 |
| DUGS1_vs_4 | 23 | | 0 |
| DUGS2_vs_3 | 0 | | 14 |
| DUGS2_vs_4 | 13 | | 19 |
| DUGS3_vs_4 | 33 | | 14 |
| DUGS1_vs_2 | 6 | Hydrolase activity | 6 |
| DUGS1_vs_3 | 0 | | 9 |
| DUGS1_vs_4 | 0 | | 0 |
| DUGS2_vs_3 | 19 | | 23 |
| DUGS2_vs_4 | 20 | | 0 |
| DUGS3_vs_4 | 0 | | 0 |
| DUGS1_vs_2 | 5 | Hydrolase activity, hydrolyzing O-glycosyl compounds | 0 |
| DUGS1_vs_3 | 0 | | 11 |
| DUGS1_vs_4 | 0 | | 9 |
| DUGS2_vs_3 | 0 | | 0 |
| DUGS2_vs_4 | 0 | | 10 |
| DUGS3_vs_4 | 22 | | 0 |
| DUGS1_vs_2 | 0 | Inorganic phosphate transmembrane transporter activity | 0 |
| DUGS1_vs_3 | 0 | | 8 |
| DUGS1_vs_4 | 0 | | 10 |
| DUGS2_vs_3 | 0 | | 0 |
| DUGS2_vs_4 | 14 | | 0 |
| DUGS3_vs_4 | 0 | | 0 |
| DUGS1_vs_2 | 5 | Iron ion binding | 4 |
| DUGS1_vs_3 | 0 | | 10 |
| DUGS1_vs_4 | 0 | | 0 |
| DUGS2_vs_3 | 0 | | 12 |
| DUGS2_vs_4 | 0 | | 0 |
| DUGS3_vs_4 | 0 | | 0 |
| DUGS1_vs_2 | 0 | Kinase activity | 4 |
| DUGS1_vs_3 | 0 | | 8 |
| DUGS1_vs_4 | 0 | | 0 |
| DUGS2_vs_3 | 0 | | 16 |
| DUGS2_vs_4 | 15 | | 0 |
| DUGS3_vs_4 | 0 | | 22 |
| DUGS1_vs_2 | 0 | Lipid transporter activity | 0 |
| DUGS1_vs_3 | 0 | | 0 |
| DUGS1_vs_4 | 0 | | 0 |
| DUGS2_vs_3 | 0 | | 16 |
| DUGS2_vs_4 | 13 | | 23 |
| DUGS3_vs_4 | 0 | | 119 |
| DUGS1_vs_2 | 5 | Magnesium ion binding | 0 |
| DUGS1_vs_3 | 9 | | 13 |
| DUGS1_vs_4 | 27 | | 0 |
| DUGS2_vs_3 | 16 | | 32 |
| DUGS2_vs_4 | 23 | | 83 |
| DUGS3_vs_4 | 36 | | 0 |
| DUGS1_vs_2 | 0 | Metal ion binding | 0 |
| DUGS1_vs_3 | 15 | | 13 |
| DUGS1_vs_4 | 25 | | 0 |
| DUGS2_vs_3 | 14 | | 75 |
| DUGS2_vs_4 | 31 | | 0 |
| DUGS3_vs_4 | 36 | | 0 |
| DUGS1_vs_2 | 0 | Metalloendopeptidase activity | 0 |
| DUGS1_vs_3 | 0 | | 11 |
| DUGS1_vs_4 | 0 | | 73 |
| DUGS2_vs_3 | 0 | | 0 |
| DUGS2_vs_4 | 0 | | 0 |
| DUGS3_vs_4 | 24 | | 57 |

continued

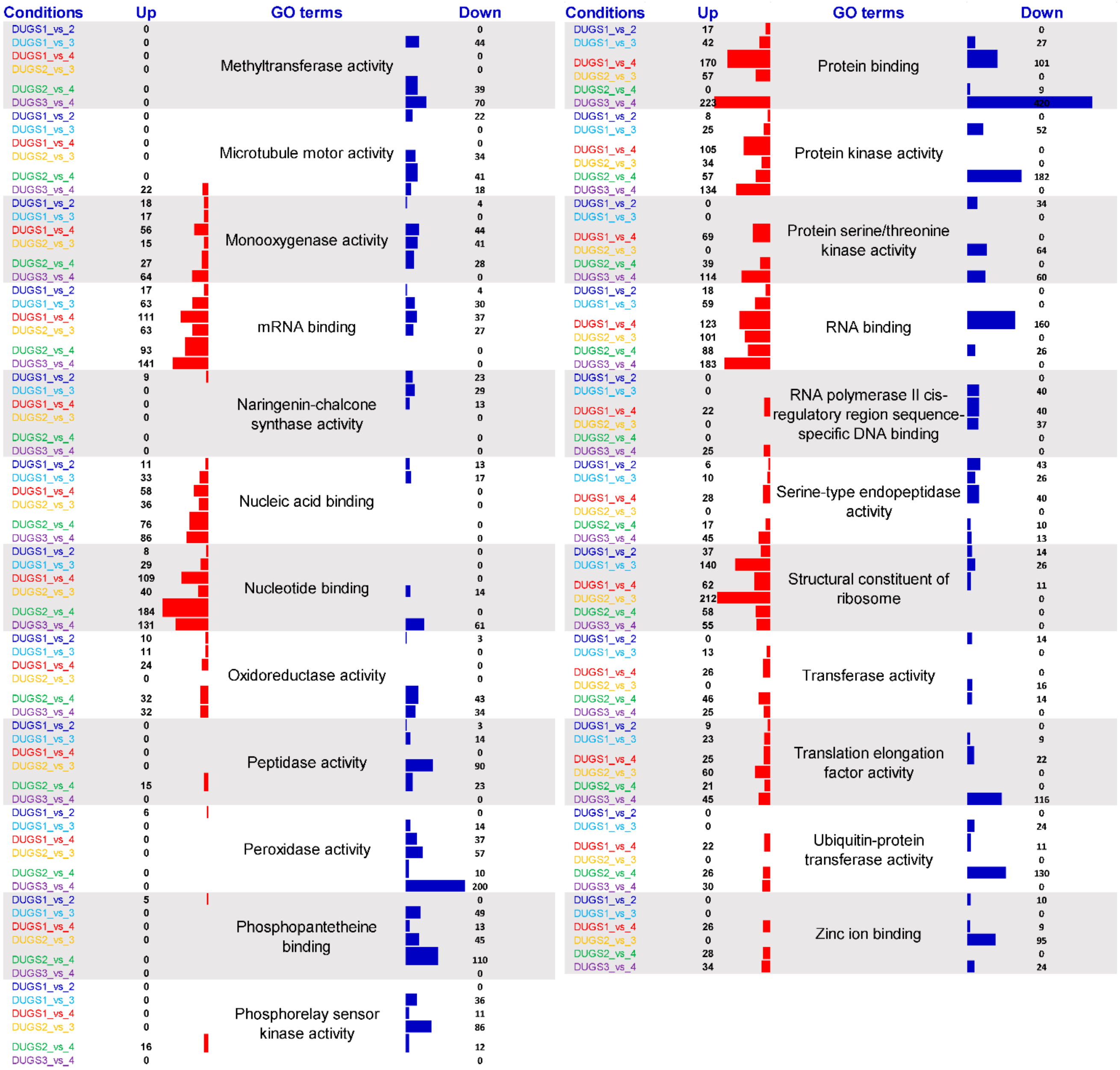


**Figure S6** Gene Ontology (GO) enrichment analysis of top 50 pathways of **(a)** biological processes (BP), **(b)** cellular components (CC), and **(c)** molecular functions (MF) for the differentially regulated genes of *D. hirsuta* in each of the six paired-wise comparisons of all four seasons (DUGS1_vs _DUGS2, DUGS1_vs_DUGS3, DUGS1 _vs _DUGS4, DUGS2_vs _DUGS3, DUGS2_vs _DUGS4 and DUGS3_vs_DUGS4). Values inside the parenthesis in front of each bars represents the enrichment counts. DU indicates *D. hirsuta*. GS1 is pre-monsoon, GS2 is monsoon, GS3 is post-monsoon, and GS4 is fruiting seasons.

## (a) BP

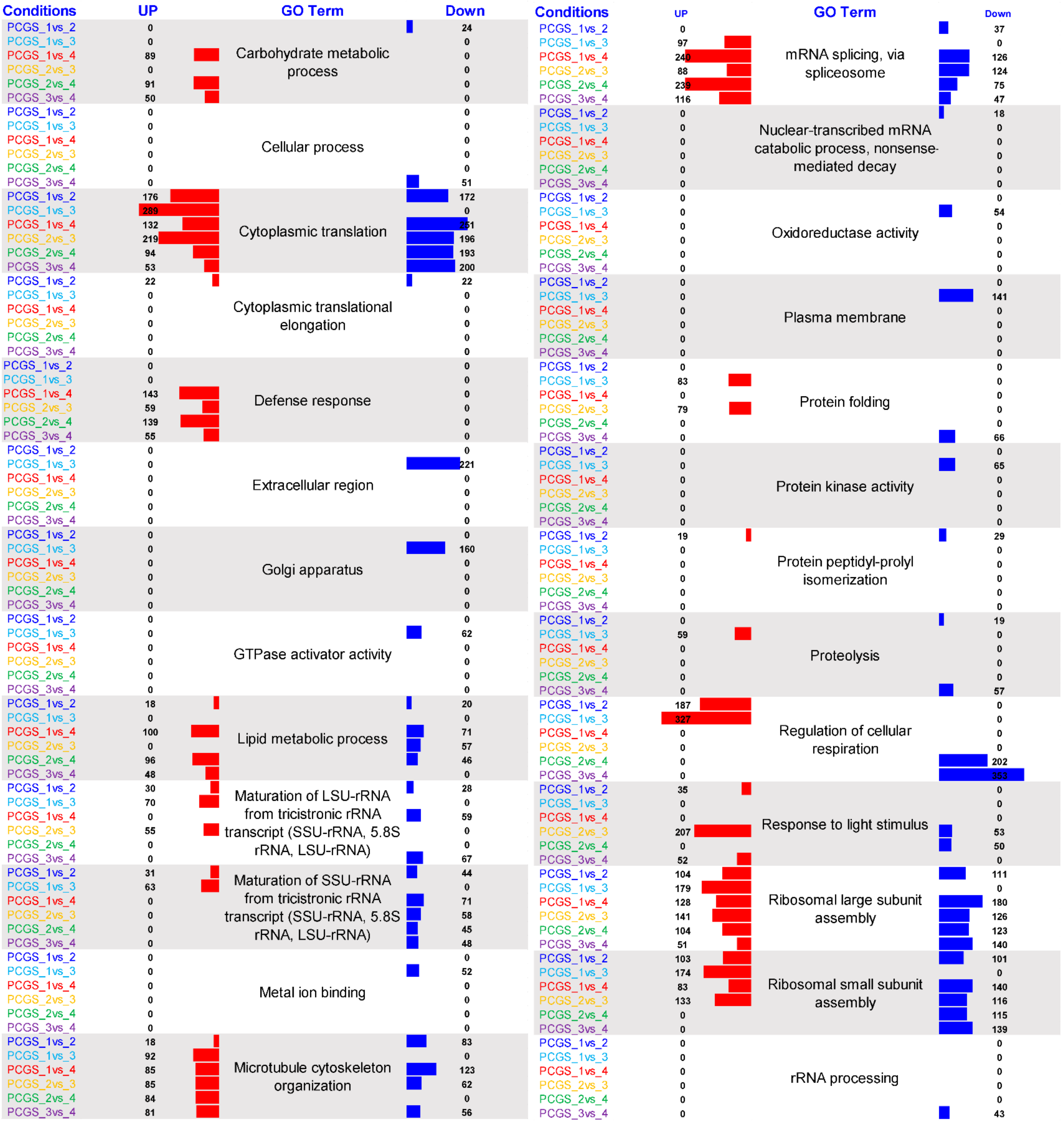

Conditions
UP
GO Term
Down
Carbohydrate metabolic process
PCGS_1vs_2 0 24
PCGS_1vs_3 0 0
PCGS_1vs_4 89 0
PCGS_2vs_3 0 0
PCGS_2vs_4 91 0
PCGS_3vs_4 50 0
Cellular process
PCGS_1vs_2 0 0
PCGS_1vs_3 0 0
PCGS_1vs_4 0 0
PCGS_2vs_3 0 0
PCGS_2vs_4 0 0
PCGS_3vs_4 0 51
Cytoplasmic translation
PCGS_1vs_2 176 172
PCGS_1vs_3 289 0
PCGS_1vs_4 132 251
PCGS_2vs_3 219 196
PCGS_2vs_4 94 193
PCGS_3vs_4 53 200
Cytoplasmic translational elongation
PCGS_1vs_2 22 22
PCGS_1vs_3 0 0
PCGS_1vs_4 0 0
PCGS_2vs_3 0 0
PCGS_2vs_4 0 0
PCGS_3vs_4 0 0
Defense response
PCGS_1vs_2 0 0
PCGS_1vs_3 0 0
PCGS_1vs_4 143 0
PCGS_2vs_3 59 0
PCGS_2vs_4 139 0
PCGS_3vs_4 55 0
Extracellular region
PCGS_1vs_2 0 0
PCGS_1vs_3 0 221
PCGS_1vs_4 0 0
PCGS_2vs_3 0 0
PCGS_2vs_4 0 0
PCGS_3vs_4 0 0
Golgi apparatus
PCGS_1vs_2 0 0
PCGS_1vs_3 0 160
PCGS_1vs_4 0 0
PCGS_2vs_3 0 0
PCGS_2vs_4 0 0
PCGS_3vs_4 0 0
GTPase activator activity
PCGS_1vs_2 0 0
PCGS_1vs_3 0 62
PCGS_1vs_4 0 0
PCGS_2vs_3 0 0
PCGS_2vs_4 0 0
PCGS_3vs_4 0 0
Lipid metabolic process
PCGS_1vs_2 18 20
PCGS_1vs_3 0 0
PCGS_1vs_4 100 71
PCGS_2vs_3 0 57
PCGS_2vs_4 96 46
PCGS_3vs_4 48 0
Maturation of LSU-rRNA from tricistronic rRNA transcript (SSU-rRNA, 5.8S rRNA, LSU-rRNA)
PCGS_1vs_2 30 28
PCGS_1vs_3 70 0
PCGS_1vs_4 0 59
PCGS_2vs_3 55 0
PCGS_2vs_4 0 0
PCGS_3vs_4 0 67
Maturation of SSU-rRNA from tricistronic rRNA transcript (SSU-rRNA, 5.8S rRNA, LSU-rRNA)
PCGS_1vs_2 31 44
PCGS_1vs_3 63 0
PCGS_1vs_4 0 71
PCGS_2vs_3 0 58
PCGS_2vs_4 0 45
PCGS_3vs_4 0 48
Metal ion binding
PCGS_1vs_2 0 0
PCGS_1vs_3 0 52
PCGS_1vs_4 0 0
PCGS_2vs_3 0 0
PCGS_2vs_4 0 0
PCGS_3vs_4 0 0
Microtubule cytoskeleton organization
PCGS_1vs_2 18 83
PCGS_1vs_3 92 0
PCGS_1vs_4 85 123
PCGS_2vs_3 85 62
PCGS_2vs_4 84 0
PCGS_3vs_4 81 56
Conditions
UP
GO Term
Down
mRNA splicing, via spliceosome
PCGS_1vs_2 0 37
PCGS_1vs_3 97 0
PCGS_1vs_4 240 126
PCGS_2vs_3 88 124
PCGS_2vs_4 239 75
PCGS_3vs_4 116 47
Nuclear-transcribed mRNA catabolic process, nonsense-mediated decay
PCGS_1vs_2 0 18
PCGS_1vs_3 0 0
PCGS_1vs_4 0 0
PCGS_2vs_3 0 0
PCGS_2vs_4 0 0
PCGS_3vs_4 0 0
Oxidoreductase activity
PCGS_1vs_2 0 0
PCGS_1vs_3 0 54
PCGS_1vs_4 0 0
PCGS_2vs_3 0 0
PCGS_2vs_4 0 0
PCGS_3vs_4 0 0
Plasma membrane
PCGS_1vs_2 0 0
PCGS_1vs_3 0 141
PCGS_1vs_4 0 0
PCGS_2vs_3 0 0
PCGS_2vs_4 0 0
PCGS_3vs_4 0 0
Protein folding
PCGS_1vs_2 0 0
PCGS_1vs_3 83 0
PCGS_1vs_4 0 0
PCGS_2vs_3 79 0
PCGS_2vs_4 0 0
PCGS_3vs_4 0 66
Protein kinase activity
PCGS_1vs_2 0 0
PCGS_1vs_3 0 65
PCGS_1vs_4 0 0
PCGS_2vs_3 0 0
PCGS_2vs_4 0 0
PCGS_3vs_4 0 0
Protein peptidyl-prolyl isomerization
PCGS_1vs_2 19 29
PCGS_1vs_3 0 0
PCGS_1vs_4 0 0
PCGS_2vs_3 0 0
PCGS_2vs_4 0 0
PCGS_3vs_4 0 0
Proteolysis
PCGS_1vs_2 0 19
PCGS_1vs_3 59 0
PCGS_1vs_4 0 0
PCGS_2vs_3 0 0
PCGS_2vs_4 0 0
PCGS_3vs_4 0 57
Regulation of cellular respiration
PCGS_1vs_2 187 0
PCGS_1vs_3 327 0
PCGS_1vs_4 0 0
PCGS_2vs_3 0 0
PCGS_2vs_4 0 202
PCGS_3vs_4 0 353
Response to light stimulus
PCGS_1vs_2 35 0
PCGS_1vs_3 0 0
PCGS_1vs_4 0 0
PCGS_2vs_3 207 53
PCGS_2vs_4 0 50
PCGS_3vs_4 52 0
Ribosomal large subunit assembly
PCGS_1vs_2 104 111
PCGS_1vs_3 179 0
PCGS_1vs_4 128 180
PCGS_2vs_3 141 126
PCGS_2vs_4 104 123
PCGS_3vs_4 51 140
Ribosomal small subunit assembly
PCGS_1vs_2 103 101
PCGS_1vs_3 174 0
PCGS_1vs_4 83 140
PCGS_2vs_3 133 116
PCGS_2vs_4 0 115
PCGS_3vs_4 0 139
rRNA processing
PCGS_1vs_2 0 0
PCGS_1vs_3 0 0
PCGS_1vs_4 0 0
PCGS_2vs_3 0 0
PCGS_2vs_4 0 0
PCGS_3vs_4 0 43

## (b) CC

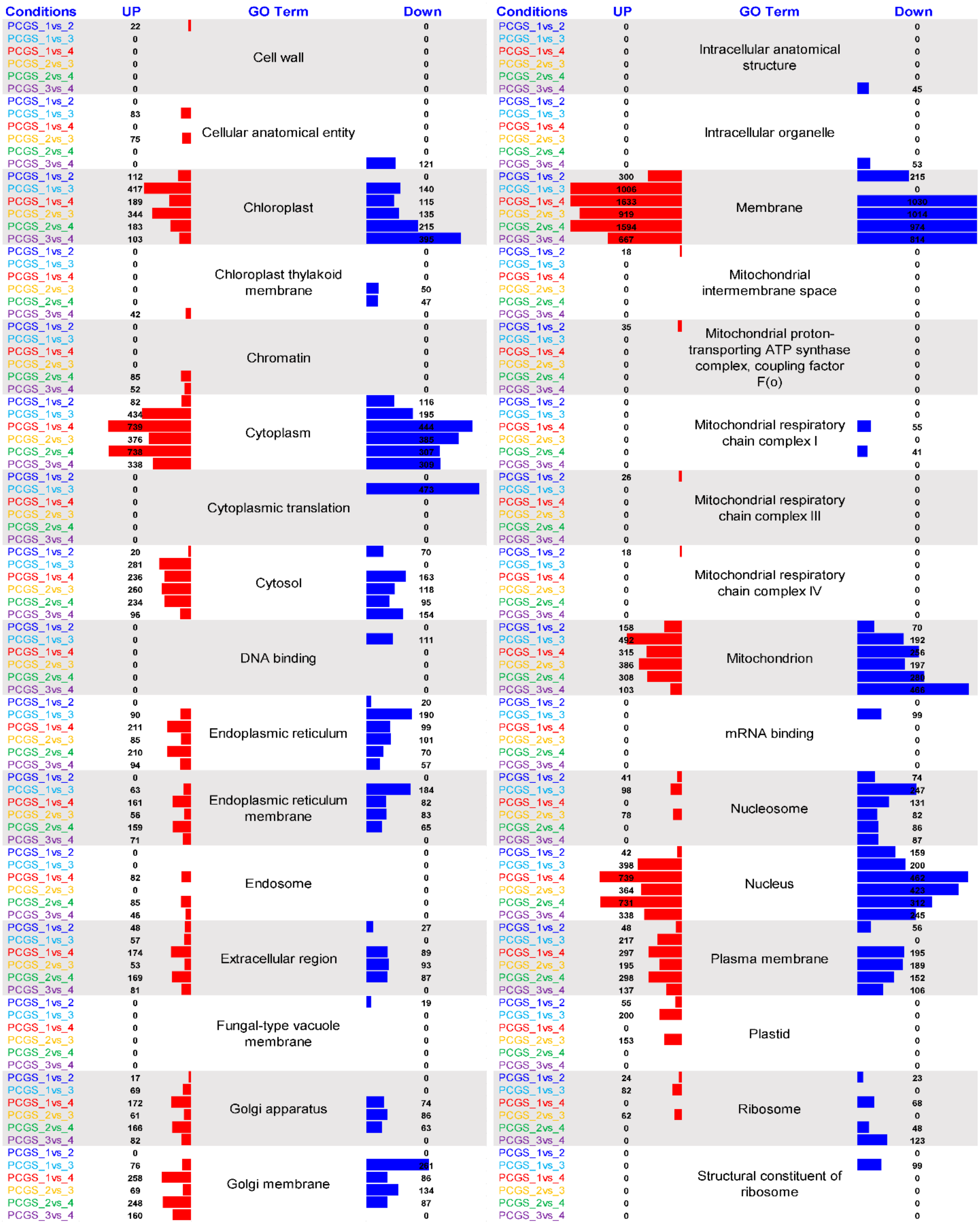
Conditions
UP
GO Term
Down
PCGS_1vs_2
PCGS_1vs_3
PCGS_1vs_4
PCGS_2vs_3
PCGS_2vs_4
PCGS_3vs_4
Cell wall
Cellular anatomical entity
Chloroplast
Chloroplast thylakoid membrane
Chromatin
Cytoplasm
Cytoplasmic translation
Cytosol
DNA binding
Endoplasmic reticulum
Endoplasmic reticulum membrane
Endosome
Extracellular region
Fungal-type vacuole membrane
Golgi apparatus
Golgi membrane
Intracellular anatomical structure
Intracellular organelle
Membrane
Mitochondrial intermembrane space
Mitochondrial proton-transporting ATP synthase complex, coupling factor F(o)
Mitochondrial respiratory chain complex I
Mitochondrial respiratory chain complex III
Mitochondrial respiratory chain complex IV
Mitochondrion
mRNA binding
Nucleosome
Nucleus
Plasma membrane
Plastid
Ribosome
Structural constituent of ribosome

**(c) MF**

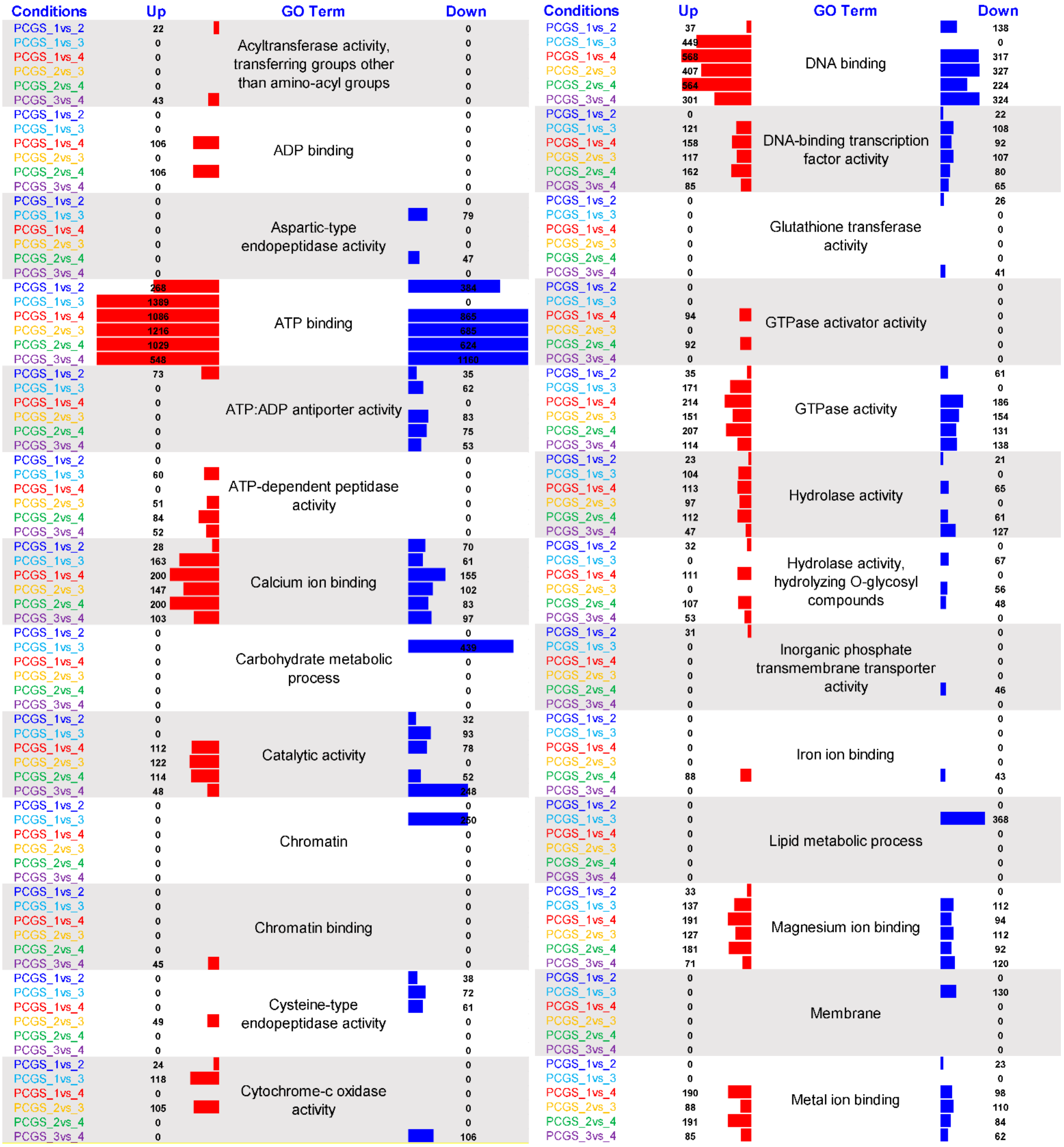


continued

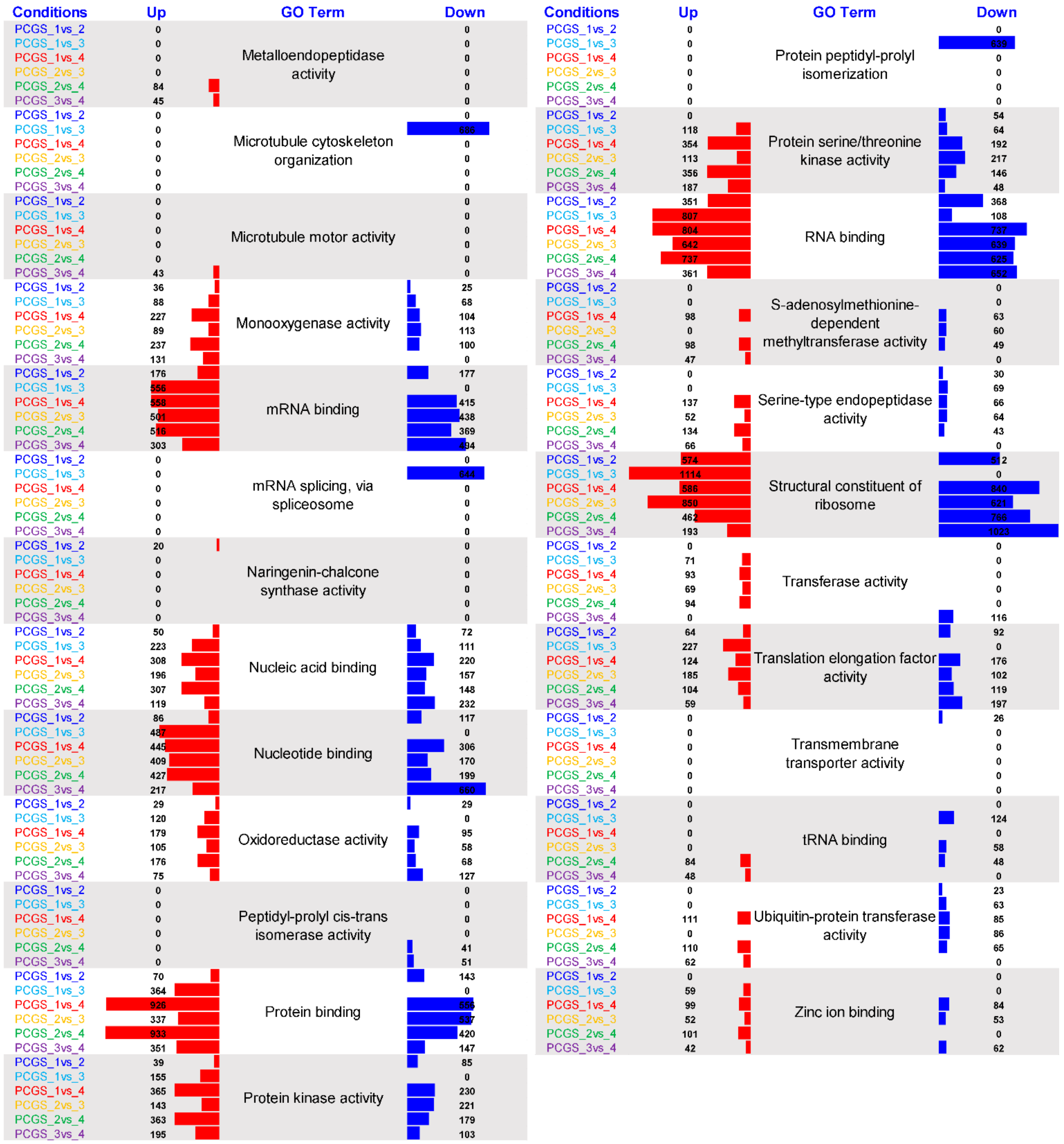


**Figure S7** Gene Ontology (GO) enrichment analysis of top ten pathways of **(a)** biological processes (BP), **(b)** cellular components (CC), and **(c)** molecular functions (MF) for the differentially regulated genes of *P. appendiculatum* in each of the six paired-wise comparisons of all four seasons (DUGS1_vs _DUGS2, DUGS1_vs_DUGS3, DUGS1 _vs _DUGS4, DUGS2_vs _DUGS3, DUGS2_vs _DUGS4 and DUGS3_vs_DUGS4). Values inside the parenthesis in front of each bars represents the enrichment counts. PC indicates *P. appendiculatum*. GS1 is pre-monsoon, GS2 is monsoon, GS3 is post-monsoon, and GS4 is fruiting seasons.

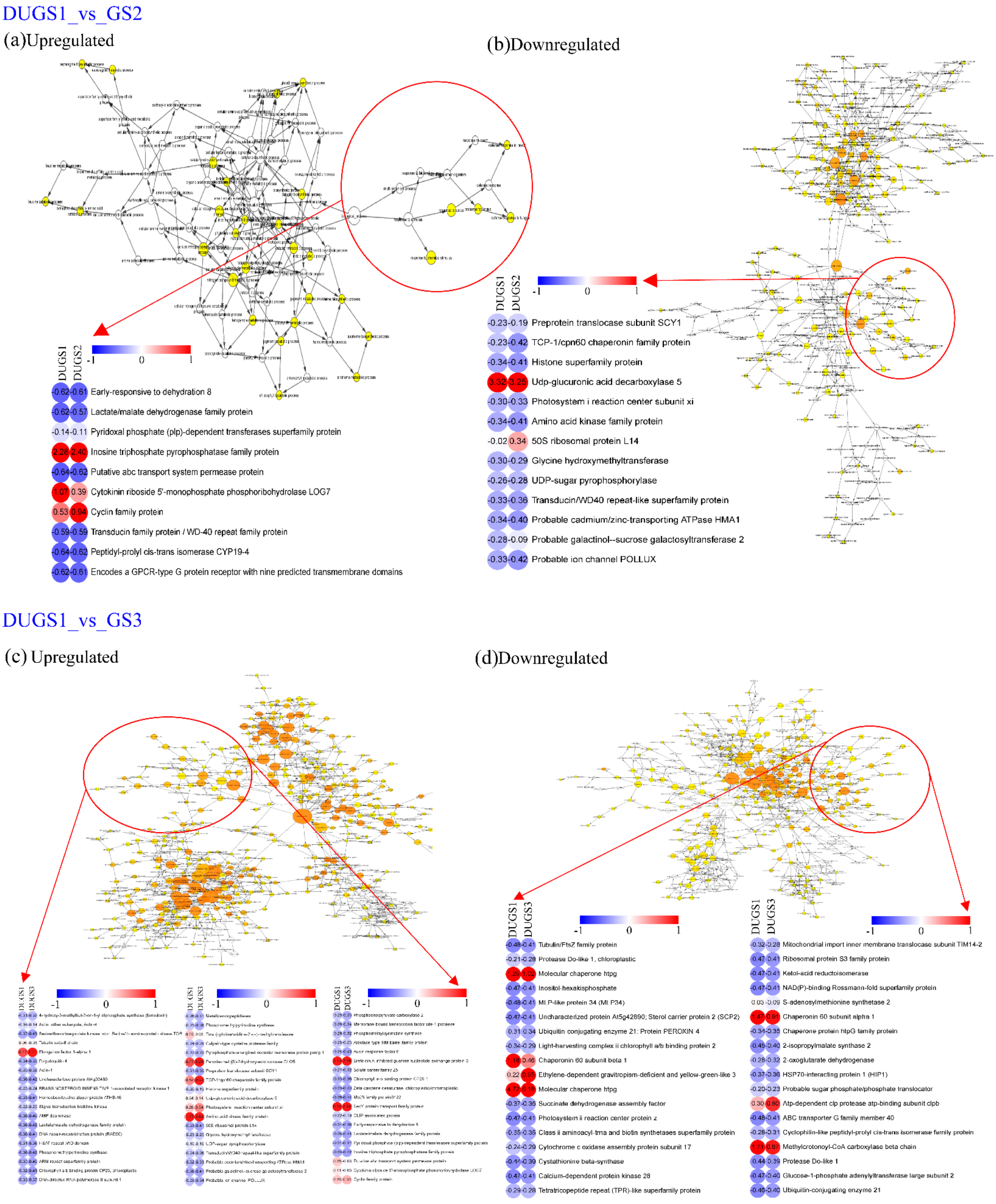
DUGS1_vs_GS2
(a)Upregulated
DUGS1
DUGS2
-1
0
1
-0.62 -0.61 Early-responsive to dehydration 8
-0.62 -0.57 Lactate/malate dehydrogenase family protein
-0.14 -0.11 Pyridoxal phosphate (plp)-dependent transferases superfamily protein
2.28 2.40 Inosine triphosphate pyrophosphatase family protein
-0.64 -0.62 Putative abc transport system permease protein
1.07 0.39 Cytokinin riboside 5'-monophosphate phosphoribohydrolase LOG7
0.53 0.94 Cyclin family protein
-0.59 -0.59 Transducin family protein / WD-40 repeat family protein
-0.64 -0.62 Peptidyl-prolyl cis-trans isomerase CYP19-4
-0.62 -0.61 Encodes a GPCR-type G protein receptor with nine predicted transmembrane domains
(b)Downregulated
DUGS1
DUGS2
-1
0
1
-0.23 -0.19 Preprotein translocase subunit SCY1
-0.23 -0.42 TCP-1/cpn60 chaperonin family protein
-0.34 -0.41 Histone superfamily protein
3.32 3.25 Udp-glucuronic acid decarboxylase 5
-0.30 -0.33 Photosystem i reaction center subunit xi
-0.34 -0.41 Amino acid kinase family protein
-0.02 0.34 50S ribosomal protein L14
-0.30 -0.29 Glycine hydroxymethyltransferase
-0.26 -0.28 UDP-sugar pyrophosphorylase
-0.33 -0.36 Transducin/WD40 repeat-like superfamily protein
-0.34 -0.40 Probable cadmium/zinc-transporting ATPase HMA1
-0.28 -0.09 Probable galactinol--sucrose galactosyltransferase 2
-0.33 -0.42 Probable ion channel POLLUX
DUGS1_vs_GS3
(c) Upregulated
(d)Downregulated
DUGS1
DUGS3
-1
0
1
Tubulin/FtsZ family protein
Protease Do-like 1, chloroplastic
Molecular chaperone htpg
Inositol-hexakisphosphate
MLP-like protein 34 (MLP34)
Uncharacterized protein At5g42890; Sterol carrier protein 2 (SCP2)
Ubiquitin conjugating enzyme 21; Protein PEROXIN 4
Light-harvesting complex ii chlorophyll a/b binding protein 2
Chaperonin 60 subunit beta 1
Ethylene-dependent gravitropism-deficient and yellow-green-like 3
Molecular chaperone htpg
Succinate dehydrogenase assembly factor
Photosystem ii reaction center protein z
Class ii aminoacyl-tRNA and biotin synthetases superfamily protein
Cytochrome c oxidase assembly protein subunit 17
Cystathionine beta-synthase
Calcium-dependent protein kinase 28
Tetratricopeptide repeat (TPR)-like superfamily protein
DUGS1
DUGS3
-1
0
1
Mitochondrial import inner membrane translocase subunit TIM14-2
Ribosomal protein S3 family protein
Ketol-acid reductoisomerase
NAD(P)-binding Rossmann-fold superfamily protein
S-adenosylmethionine synthetase 2
Chaperonin 60 subunit alpha 1
Chaperone protein htpG family protein
2-isopropylmalate synthase 2
2-oxoglutarate dehydrogenase
HSP70-interacting protein 1 (HIP1)
Probable sugar phosphate/phosphate translocator
Atp-dependent clp protease atp-binding subunit clpb
ABC transporter G family member 40
Cyclophilin-like peptidyl-prolyl cis-trans isomerase family protein
Methylcrotonoyl-CoA carboxylase beta chain
Protease Do-like 1
Glucose-1-phosphate adenylyltransferase large subunit 2
Ubiquitin-conjugating enzyme 21

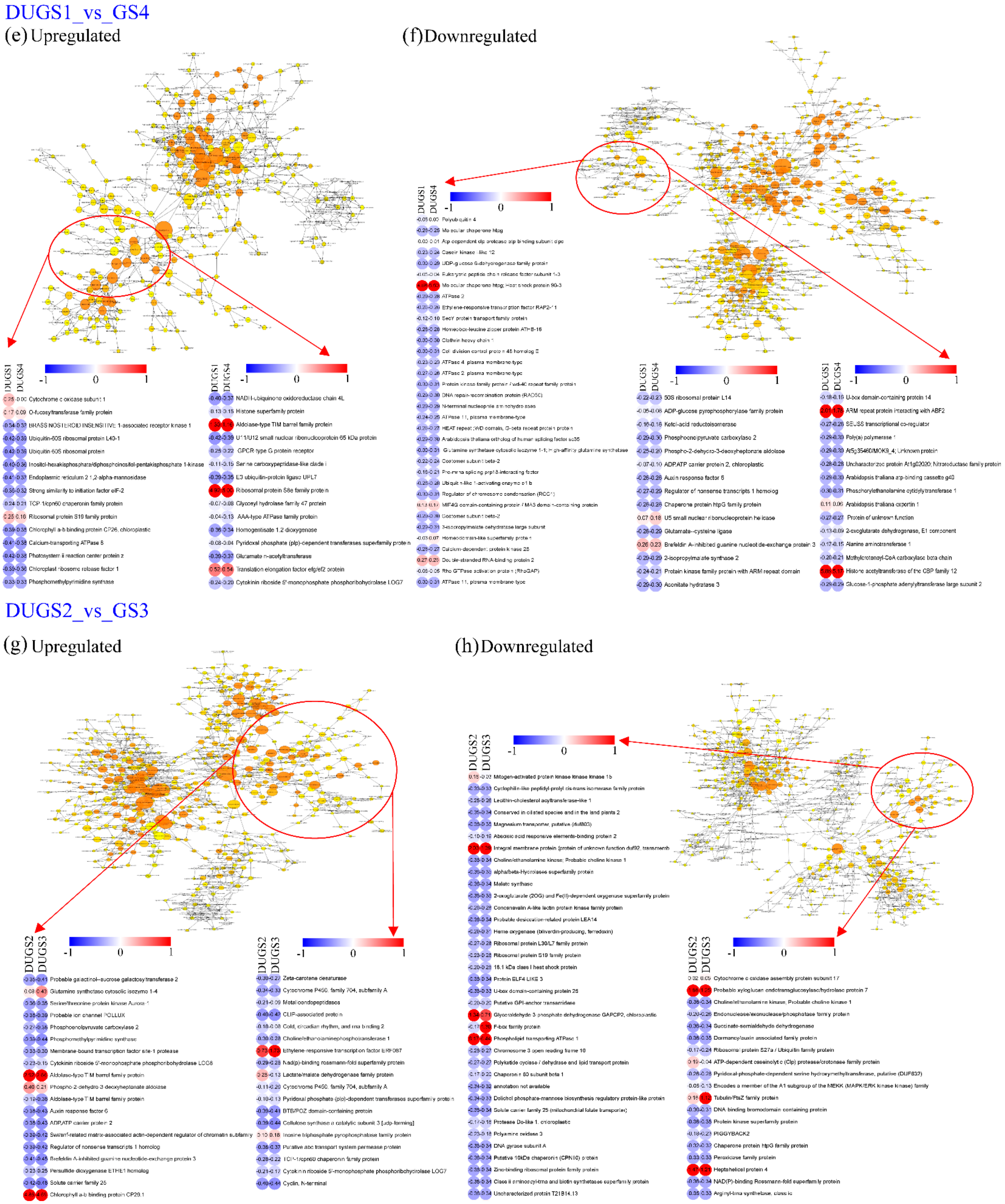
DUGS1_vs_GS4
(e) Upregulated
(f) Downregulated
DUGS2_vs_GS3
(g) Upregulated
(h) Downregulated

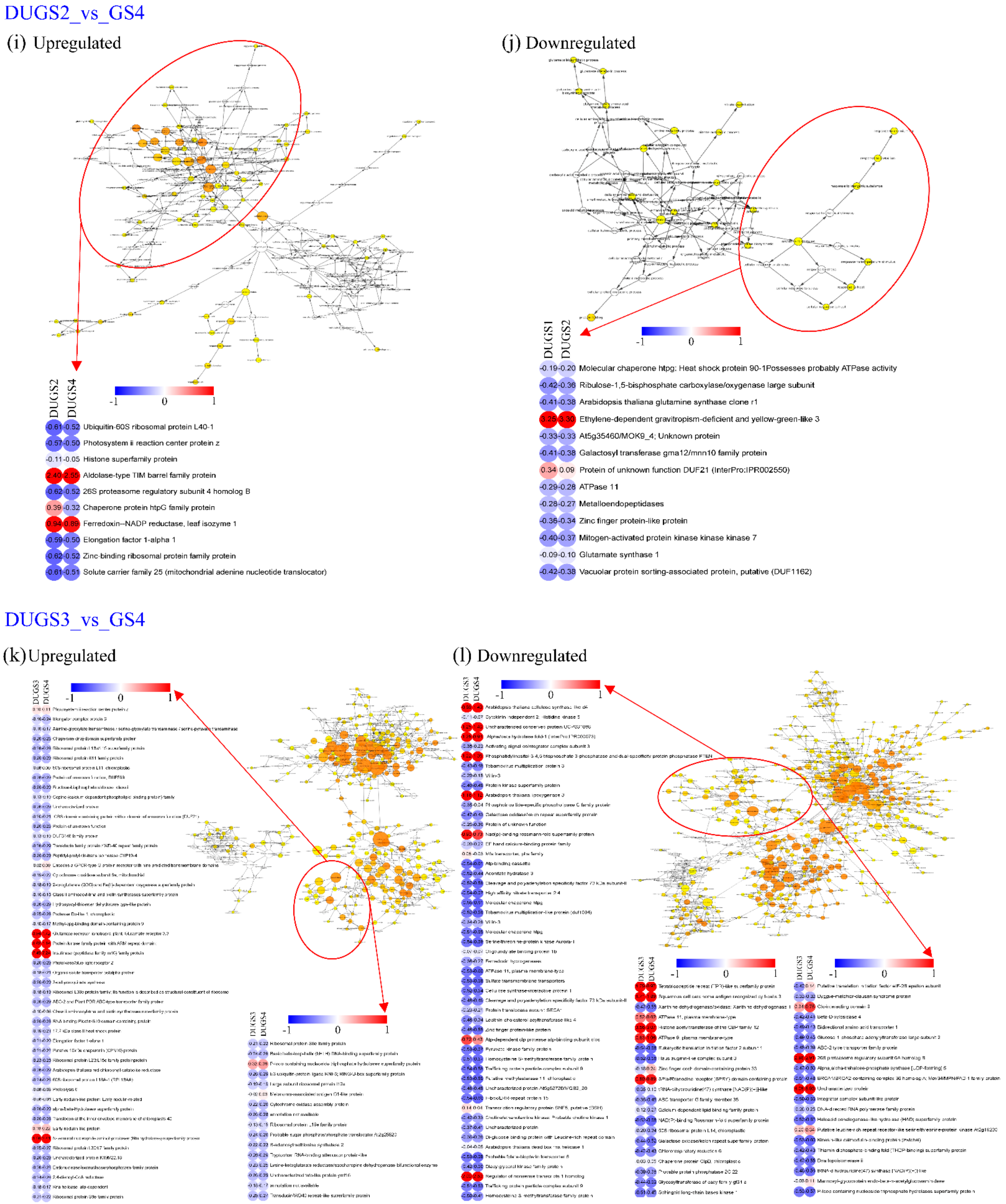
DUGS2_vs_GS4
(i) Upregulated
(j) Downregulated
DUGS2
DUGS4
-1
0
1
-0.61 -0.52 Ubiquitin-60S ribosomal protein L40-1
-0.57 -0.50 Photosystem ii reaction center protein z
-0.11 -0.05 Histone superfamily protein
2.40 2.55 Aldolase-type TIM barrel family protein
-0.62 -0.52 26S proteasome regulatory subunit 4 homolog B
0.39 -0.32 Chaperone protein htpG family protein
0.94 0.89 Ferredoxin--NADP reductase, leaf isozyme 1
-0.59 -0.50 Elongation factor 1-alpha 1
-0.62 -0.52 Zinc-binding ribosomal protein family protein
-0.61 -0.51 Solute carrier family 25 (mitochondrial adenine nucleotide translocator)
DUGS1
DUGS2
-0.19 -0.20 Molecular chaperone htpg: Heat shock protein 90-1Possesses probably ATPase activity
-0.42 -0.36 Ribulose-1,5-bisphosphate carboxylase/oxygenase large subunit
-0.41 -0.38 Arabidopsis thaliana glutamine synthase clone r1
3.25 3.30 Ethylene-dependent gravitropism-deficient and yellow-green-like 3
-0.33 -0.33 At5g35460/MOK9_4; Unknown protein
-0.41 -0.38 Galactosyl transferase gma12/mnn10 family protein
0.34 0.09 Protein of unknown function DUF21 (InterPro:IPR002550)
-0.29 -0.28 ATPase 11
-0.28 -0.27 Metalloendopeptidases
-0.36 -0.34 Zinc finger protein-like protein
-0.40 -0.37 Mitogen-activated protein kinase kinase kinase 7
-0.09 -0.10 Glutamate synthase 1
-0.42 -0.38 Vacuolar protein sorting-associated protein, putative (DUF1162)
DUGS3_vs_GS4
(k) Upregulated
(l) Downregulated
DUGS3
DUGS4

**Figure S8** Gene expression networks analysis of differentially expressed genes identified in each of the six paired-wise comparisons of all four seasons of *D. hirsuta*. Figures (a) to (l) depict the stress-related genes along with the complete gene set enrichment analysis (GSEA) network observed in the six inter-season comparison of DUGS1_vs _DUGS2, DUGS1_vs_DUGS3, DUGS1 _vs _DUGS4, DUGS2_vs _DUGS3, DUGS2_vs _DUGS4 and DUGS3_vs_DUGS4. The color scale (from white, blue to red) is based on (corrected) p-value (5% FDR, p = 0.05). Dark red categories are most significantly overrepresented, white nodes are not significantly overrepresented (NA); they are included to show other nodes in the context of the GO hierarchy.

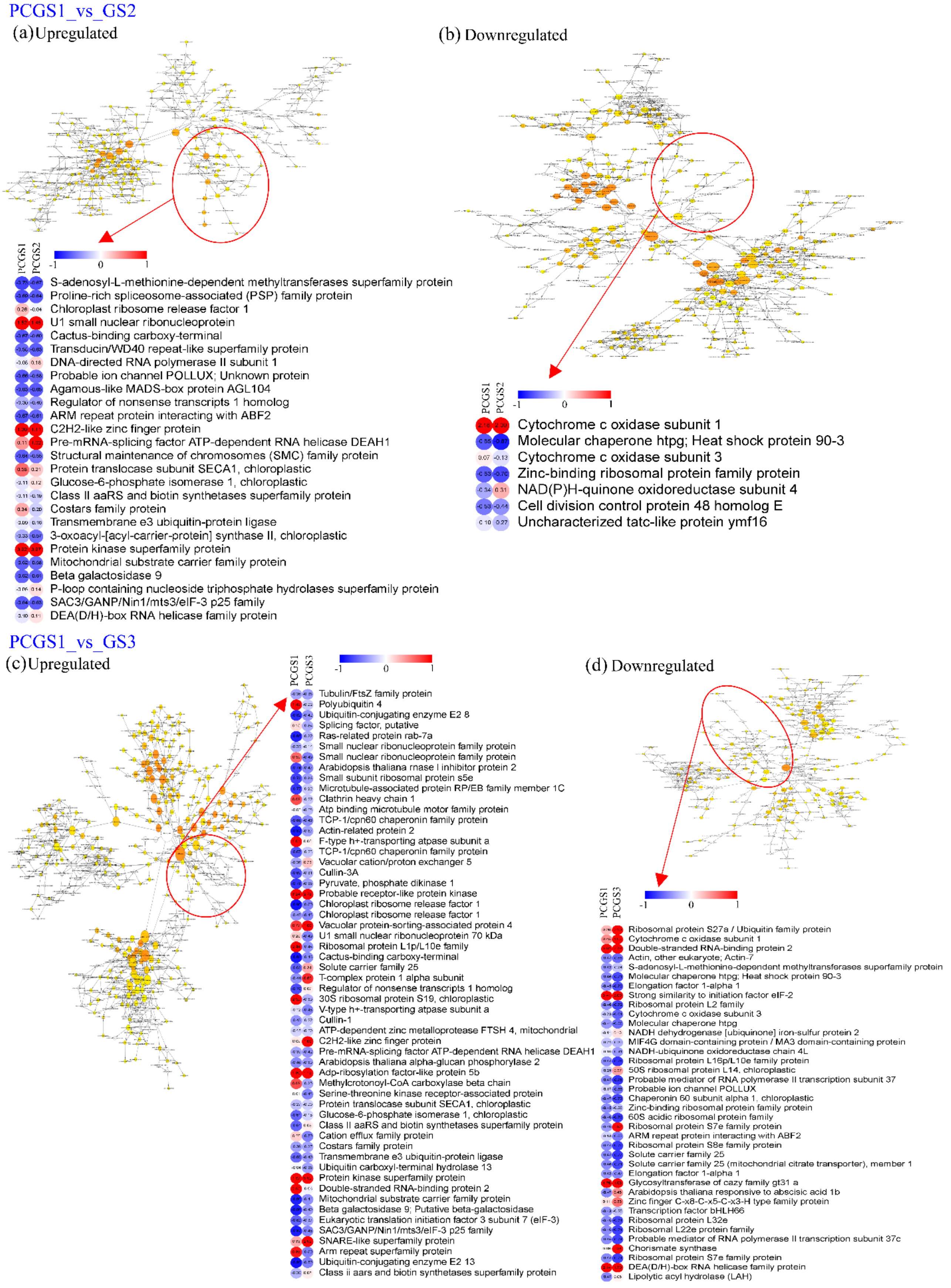
PCGS1_vs_GS2
(a)Upregulated
(b) Downregulated
PCGS1
PCGS2
-1
0
1
S-adenosyl-L-methionine-dependent methyltransferases superfamily protein
Proline-rich spliceosome-associated (PSP) family protein
Chloroplast ribosome release factor 1
U1 small nuclear ribonucleoprotein
Cactus-binding carboxy-terminal
Transducin/WD40 repeat-like superfamily protein
DNA-directed RNA polymerase II subunit 1
Probable ion channel POLLUX; Unknown protein
Agamous-like MADS-box protein AGL104
Regulator of nonsense transcripts 1 homolog
ARM repeat protein interacting with ABF2
C2H2-like zinc finger protein
Pre-mRNA-splicing factor ATP-dependent RNA helicase DEAH1
Structural maintenance of chromosomes (SMC) family protein
Protein translocase subunit SECA1, chloroplastic
Glucose-6-phosphate isomerase 1, chloroplastic
Class II aaRS and biotin synthetases superfamily protein
Costars family protein
Transmembrane e3 ubiquitin-protein ligase
3-oxoacyl-[acyl-carrier-protein] synthase II, chloroplastic
Protein kinase superfamily protein
Mitochondrial substrate carrier family protein
Beta galactosidase 9
P-loop containing nucleoside triphosphate hydrolases superfamily protein
SAC3/GANP/Nin1/mts3/eIF-3 p25 family
DEA(D/H)-box RNA helicase family protein
PCGS1
PCGS2
-1
0
1
Cytochrome c oxidase subunit 1
Molecular chaperone htpg; Heat shock protein 90-3
Cytochrome c oxidase subunit 3
Zinc-binding ribosomal protein family protein
NAD(P)H-quinone oxidoreductase subunit 4
Cell division control protein 48 homolog E
Uncharacterized tatc-like protein ymf16
PCGS1_vs_GS3
(c)Upregulated
(d) Downregulated
PCGS1
PCGS3
-1
0
1
Tubulin/FtsZ family protein
Polyubiquitin 4
Ubiquitin-conjugating enzyme E2 8
Splicing factor, putative
Ras-related protein rab-7a
Small nuclear ribonucleoprotein family protein
Small nuclear ribonucleoprotein family protein
Arabidopsis thaliana rnase l inhibitor protein 2
Small subunit ribosomal protein s5e
Microtubule-associated protein RP/EB family member 1C
Clathrin heavy chain 1
Atp binding microtubule motor family protein
TCP-1/cpn60 chaperonin family protein
Actin-related protein 2
F-type h+-transporting atpase subunit a
TCP-1/cpn60 chaperonin family protein
Vacuolar cation/proton exchanger 5
Cullin-3A
Pyruvate, phosphate dikinase 1
Probable receptor-like protein kinase
Chloroplast ribosome release factor 1
Chloroplast ribosome release factor 1
Vacuolar protein-sorting-associated protein 4
U1 small nuclear ribonucleoprotein 70 kDa
Ribosomal protein L1p/L10e family
Cactus-binding carboxy-terminal
Solute carrier family 25
T-complex protein 1 alpha subunit
Regulator of nonsense transcripts 1 homolog
30S ribosomal protein S19, chloroplastic
V-type h+-transporting atpase subunit a
Cullin-1
ATP-dependent zinc metalloprotease FTSH 4, mitochondrial
C2H2-like zinc finger protein
Pre-mRNA-splicing factor ATP-dependent RNA helicase DEAH1
Arabidopsis thaliana alpha-glucan phosphorylase 2
Adp-ribosylation factor-like protein 5b
Methylcrotonoyl-CoA carboxylase beta chain
Serine-threonine kinase receptor-associated protein
Protein translocase subunit SECA1, chloroplastic
Glucose-6-phosphate isomerase 1, chloroplastic
Class II aaRS and biotin synthetases superfamily protein
Cation efflux family protein
Costars family protein
Transmembrane e3 ubiquitin-protein ligase
Ubiquitin carboxyl-terminal hydrolase 13
Protein kinase superfamily protein
Double-stranded RNA-binding protein 2
Mitochondrial substrate carrier family protein
Beta galactosidase 9; Putative beta-galactosidase
Eukaryotic translation initiation factor 3 subunit 7 (eIF-3)
SAC3/GANP/Nin1/mts3/eIF-3 p25 family
SNARE-like superfamily protein
Arm repeat superfamily protein
Ubiquitin-conjugating enzyme E2 13
Class ii aars and biotin synthetases superfamily protein
PCGS1
PCGS3
-1
0
1
Ribosomal protein S27a / Ubiquitin family protein
Cytochrome c oxidase subunit 1
Double-stranded RNA-binding protein 2
Actin, other eukaryote; Actin-7
S-adenosyl-L-methionine-dependent methyltransferases superfamily protein
Molecular chaperone htpg; Heat shock protein 90-3
Elongation factor 1-alpha 1
Strong similarity to initiation factor eIF-2
Ribosomal protein L2 family
Cytochrome c oxidase subunit 3
Molecular chaperone htpg
NADH dehydrogenase [ubiquinone] iron-sulfur protein 2
MIF4G domain-containing protein / MA3 domain-containing protein
NADH-ubiquinone oxidoreductase chain 4L
Ribosomal protein L16p/L10e family protein
50S ribosomal protein L14, chloroplastic
Probable mediator of RNA polymerase II transcription subunit 37
Probable ion channel POLLUX
Chaperonin 60 subunit alpha 1, chloroplastic
Zinc-binding ribosomal protein family protein
60S acidic ribosomal protein family
Ribosomal protein S7e family protein
ARM repeat protein interacting with ABF2
Ribosomal protein S8e family protein
Solute carrier family 25
Solute carrier family 25 (mitochondrial citrate transporter), member 1
Elongation factor 1-alpha 1
Glycosyltransferase of cazy family gt31 a
Arabidopsis thaliana responsive to abscisic acid 1b
Zinc finger C-x8-C-x5-C-x3-H type family protein
Transcription factor bHLH66
Ribosomal protein L32e
Ribosomal L22e protein family
Probable mediator of RNA polymerase II transcription subunit 37c
Chorismate synthase
Ribosomal protein S7e family protein
DEA(D/H)-box RNA helicase family protein
Lipolytic acyl hydrolase (LAH)

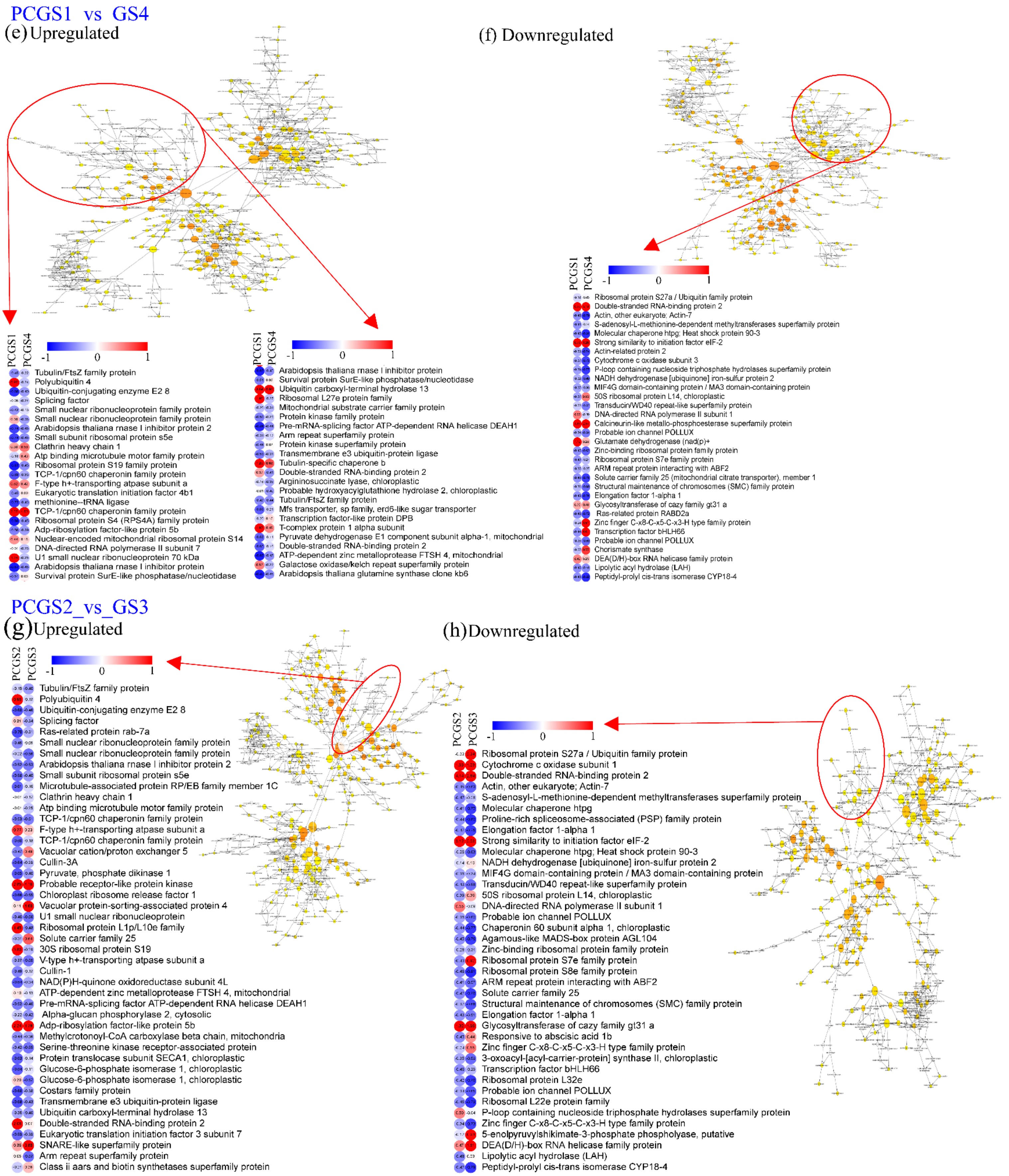
PCGS1 vs GS4
(e) Upregulated
(f) Downregulated
PCGS1
PCGS4
-1
0
1
Tubulin/FtsZ family protein
Polyubiquitin 4
Ubiquitin-conjugating enzyme E2 8
Splicing factor
Small nuclear ribonucleoprotein family protein
Small nuclear ribonucleoprotein family protein
Arabidopsis thaliana rnase l inhibitor protein 2
Small subunit ribosomal protein s5e
Clathrin heavy chain 1
Atp binding microtubule motor family protein
Ribosomal protein S19 family protein
TCP-1/cpn60 chaperonin family protein
F-type h+-transporting atpase subunit a
Eukaryotic translation initiation factor 4b1
methionine--tRNA ligase
TCP-1/cpn60 chaperonin family protein
Ribosomal protein S4 (RPS4A) family protein
Adp-ribosylation factor-like protein 5b
Nuclear-encoded mitochondrial ribosomal protein S14
DNA-directed RNA polymerase II subunit 7
U1 small nuclear ribonucleoprotein 70 kDa
Arabidopsis thaliana rnase l inhibitor protein
Survival protein SurE-like phosphatase/nucleotidase
Arabidopsis thaliana rnase l inhibitor protein
Survival protein SurE-like phosphatase/nucleotidase
Ubiquitin carboxyl-terminal hydrolase 13
Ribosomal L27e protein family
Mitochondrial substrate carrier family protein
Protein kinase family protein
Pre-mRNA-splicing factor ATP-dependent RNA helicase DEAH1
Arm repeat superfamily protein
Protein kinase superfamily protein
Transmembrane e3 ubiquitin-protein ligase
Tubulin-specific chaperone b
Double-stranded RNA-binding protein 2
Argininosuccinate lyase, chloroplastic
Probable hydroxyacylglutathione hydrolase 2, chloroplastic
Tubulin/FtsZ family protein
Mfs transporter, sp family, erd6-like sugar transporter
Transcription factor-like protein DPB
T-complex protein 1 alpha subunit
Pyruvate dehydrogenase E1 component subunit alpha-1, mitochondrial
Double-stranded RNA-binding protein 2
ATP-dependent zinc metalloprotease FTSH 4, mitochondrial
Galactose oxidase/kelch repeat superfamily protein
Arabidopsis thaliana glutamine synthase clone kb6
Ribosomal protein S27a / Ubiquitin family protein
Double-stranded RNA-binding protein 2
Actin, other eukaryote; Actin-7
S-adenosyl-L-methionine-dependent methyltransferases superfamily protein
Molecular chaperone htpg; Heat shock protein 90-3
Strong similarity to initiation factor eIF-2
Actin-related protein 2
Cytochrome c oxidase subunit 3
P-loop containing nucleoside triphosphate hydrolases superfamily protein
NADH dehydrogenase [ubiquinone] iron-sulfur protein 2
MIF4G domain-containing protein / MA3 domain-containing protein
50S ribosomal protein L14, chloroplastic
Transducin/WD40 repeat-like superfamily protein
DNA-directed RNA polymerase II subunit 1
Calcineurin-like metallo-phosphoesterase superfamily protein
Probable ion channel POLLUX
Glutamate dehydrogenase (nad(p)+
Zinc-binding ribosomal protein family protein
Ribosomal protein S7e family protein
ARM repeat protein interacting with ABF2
Solute carrier family 25 (mitochondrial citrate transporter), member 1
Structural maintenance of chromosomes (SMC) family protein
Elongation factor 1-alpha 1
Glycosyltransferase of cazy family gt31 a
Ras-related protein RABD2a
Zinc finger C-x8-C-x5-C-x3-H type family protein
Transcription factor bHLH66
Probable ion channel POLLUX
Chorismate synthase
DEA(D/H)-box RNA helicase family protein
Lipolytic acyl hydrolase (LAH)
Peptidyl-prolyl cis-trans isomerase CYP18-4
PCGS2_vs_GS3
(g)Upregulated
(h)Downregulated
PCGS2
PCGS3
Tubulin/FtsZ family protein
Polyubiquitin 4
Ubiquitin-conjugating enzyme E2 8
Splicing factor
Ras-related protein rab-7a
Small nuclear ribonucleoprotein family protein
Small nuclear ribonucleoprotein family protein
Arabidopsis thaliana rnase l inhibitor protein 2
Small subunit ribosomal protein s5e
Microtubule-associated protein RP/EB family member 1C
Clathrin heavy chain 1
Atp binding microtubule motor family protein
TCP-1/cpn60 chaperonin family protein
F-type h+-transporting atpase subunit a
TCP-1/cpn60 chaperonin family protein
Vacuolar cation/proton exchanger 5
Cullin-3A
Pyruvate, phosphate dikinase 1
Probable receptor-like protein kinase
Chloroplast ribosome release factor 1
Vacuolar protein-sorting-associated protein 4
U1 small nuclear ribonucleoprotein
Ribosomal protein L1p/L10e family
Solute carrier family 25
30S ribosomal protein S19
V-type h+-transporting atpase subunit a
Cullin-1
NAD(P)H-quinone oxidoreductase subunit 4L
ATP-dependent zinc metalloprotease FTSH 4, mitochondrial
Pre-mRNA-splicing factor ATP-dependent RNA helicase DEAH1
Alpha-glucan phosphorylase 2, cytosolic
Adp-ribosylation factor-like protein 5b
Methylcrotonoyl-CoA carboxylase beta chain, mitochondria
Serine-threonine kinase receptor-associated protein
Protein translocase subunit SECA1, chloroplastic
Glucose-6-phosphate isomerase 1, chloroplastic
Glucose-6-phosphate isomerase 1, chloroplastic
Costars family protein
Transmembrane e3 ubiquitin-protein ligase
Ubiquitin carboxyl-terminal hydrolase 13
Double-stranded RNA-binding protein 2
Eukaryotic translation initiation factor 3 subunit 7
SNARE-like superfamily protein
Arm repeat superfamily protein
Class ii aars and biotin synthetases superfamily protein
Ribosomal protein S27a / Ubiquitin family protein
Cytochrome c oxidase subunit 1
Double-stranded RNA-binding protein 2
Actin, other eukaryote; Actin-7
S-adenosyl-L-methionine-dependent methyltransferases superfamily protein
Molecular chaperone htpg
Proline-rich spliceosome-associated (PSP) family protein
Elongation factor 1-alpha 1
Strong similarity to initiation factor eIF-2
Molecular chaperone htpg; Heat shock protein 90-3
NADH dehydrogenase [ubiquinone] iron-sulfur protein 2
MIF4G domain-containing protein / MA3 domain-containing protein
Transducin/WD40 repeat-like superfamily protein
50S ribosomal protein L14, chloroplastic
DNA-directed RNA polymerase II subunit 1
Probable ion channel POLLUX
Chaperonin 60 subunit alpha 1, chloroplastic
Agamous-like MADS-box protein AGL104
Zinc-binding ribosomal protein family protein
Ribosomal protein S7e family protein
Ribosomal protein S8e family protein
ARM repeat protein interacting with ABF2
Solute carrier family 25
Structural maintenance of chromosomes (SMC) family protein
Elongation factor 1-alpha 1
Glycosyltransferase of cazy family gt31 a
Responsive to abscisic acid 1b
Zinc finger C-x8-C-x5-C-x3-H type family protein
3-oxoacyl-[acyl-carrier-protein] synthase II, chloroplastic
Transcription factor bHLH66
Ribosomal protein L32e
Probable ion channel POLLUX
Ribosomal L22e protein family
P-loop containing nucleoside triphosphate hydrolases superfamily protein
Zinc finger C-x8-C-x5-C-x3-H type family protein
5-enolpyruvylshikimate-3-phosphate phospholyase, putative
DEA(D/H)-box RNA helicase family protein
Lipolytic acyl hydrolase (LAH)
Peptidyl-prolyl cis-trans isomerase CYP18-4

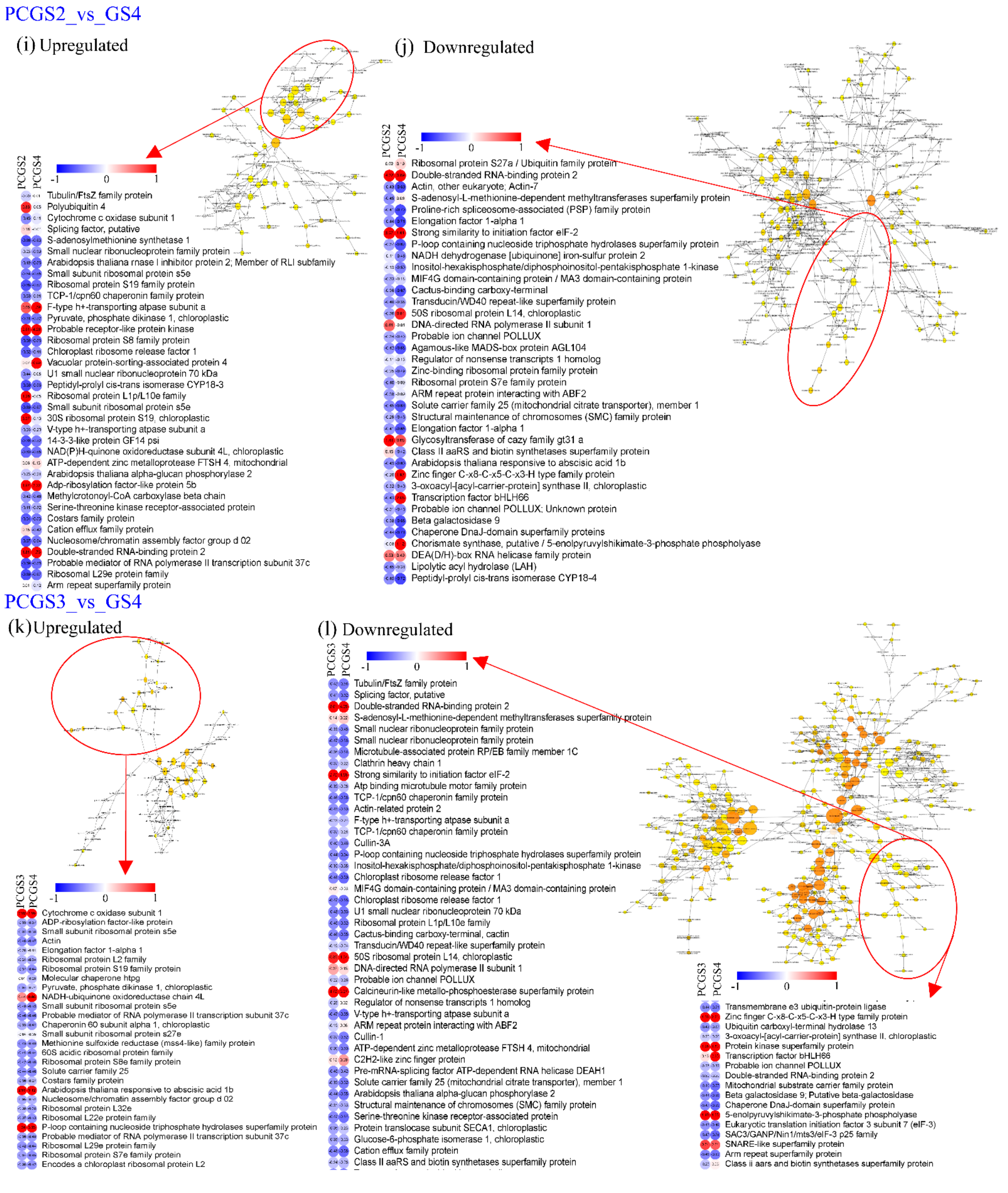
PCGS2_vs_GS4
(i) Upregulated
(j) Downregulated
PCGS2
PCGS4
-1
0
1
Tubulin/FtsZ family protein
Polyubiquitin 4
Cytochrome c oxidase subunit 1
Splicing factor, putative
S-adenosylmethionine synthetase 1
Small nuclear ribonucleoprotein family protein
Arabidopsis thaliana rnase l inhibitor protein 2; Member of RLI subfamily
Small subunit ribosomal protein s5e
Ribosomal protein S19 family protein
TCP-1/cpn60 chaperonin family protein
F-type h+-transporting atpase subunit a
Pyruvate, phosphate dikinase 1, chloroplastic
Probable receptor-like protein kinase
Ribosomal protein S8 family protein
Chloroplast ribosome release factor 1
Vacuolar protein-sorting-associated protein 4
U1 small nuclear ribonucleoprotein 70 kDa
Peptidyl-prolyl cis-trans isomerase CYP18-3
Ribosomal protein L1p/L10e family
Small subunit ribosomal protein s5e
30S ribosomal protein S19, chloroplastic
V-type h+-transporting atpase subunit a
14-3-3-like protein GF14 psi
NAD(P)H-quinone oxidoreductase subunit 4L, chloroplastic
ATP-dependent zinc metalloprotease FTSH 4, mitochondrial
Arabidopsis thaliana alpha-glucan phosphorylase 2
Adp-ribosylation factor-like protein 5b
Methylcrotonoyl-CoA carboxylase beta chain
Serine-threonine kinase receptor-associated protein
Costars family protein
Cation efflux family protein
Nucleosome/chromatin assembly factor group d 02
Double-stranded RNA-binding protein 2
Probable mediator of RNA polymerase II transcription subunit 37c
Ribosomal L29e protein family
Arm repeat superfamily protein
PCGS2
PCGS4
-1
0
1
Ribosomal protein S27a / Ubiquitin family protein
Double-stranded RNA-binding protein 2
Actin, other eukaryote; Actin-7
S-adenosyl-L-methionine-dependent methyltransferases superfamily protein
Proline-rich spliceosome-associated (PSP) family protein
Elongation factor 1-alpha 1
Strong similarity to initiation factor eIF-2
P-loop containing nucleoside triphosphate hydrolases superfamily protein
NADH dehydrogenase [ubiquinone] iron-sulfur protein 2
Inositol-hexakisphosphate/diphosphoinositol-pentakisphosphate 1-kinase
MIF4G domain-containing protein / MA3 domain-containing protein
Cactus-binding carboxy-terminal
Transducin/WD40 repeat-like superfamily protein
50S ribosomal protein L14, chloroplastic
DNA-directed RNA polymerase II subunit 1
Probable ion channel POLLUX
Agamous-like MADS-box protein AGL104
Regulator of nonsense transcripts 1 homolog
Zinc-binding ribosomal protein family protein
Ribosomal protein S7e family protein
ARM repeat protein interacting with ABF2
Solute carrier family 25 (mitochondrial citrate transporter), member 1
Structural maintenance of chromosomes (SMC) family protein
Elongation factor 1-alpha 1
Glycosyltransferase of cazy family gt31 a
Class II aaRS and biotin synthetases superfamily protein
Arabidopsis thaliana responsive to abscisic acid 1b
Zinc finger C-x8-C-x5-C-x3-H type family protein
3-oxoacyl-[acyl-carrier-protein] synthase II, chloroplastic
Transcription factor bHLH66
Probable ion channel POLLUX; Unknown protein
Beta galactosidase 9
Chaperone DnaJ-domain superfamily proteins
Chorismate synthase, putative / 5-enolpyruvylshikimate-3-phosphate phospholyase
DEA(D/H)-box RNA helicase family protein
Lipolytic acyl hydrolase (LAH)
Peptidyl-prolyl cis-trans isomerase CYP18-4
PCGS3_vs_GS4
(k) Upregulated
(l) Downregulated
PCGS3
PCGS4
-1
0
1
Cytochrome c oxidase subunit 1
ADP-ribosylation factor-like protein
Small subunit ribosomal protein s5e
Actin
Elongation factor 1-alpha 1
Ribosomal protein L2 family
Ribosomal protein S19 family protein
Molecular chaperone htpg
Pyruvate, phosphate dikinase 1, chloroplastic
NADH-ubiquinone oxidoreductase chain 4L
Small subunit ribosomal protein s5e
Probable mediator of RNA polymerase II transcription subunit 37c
Chaperonin 60 subunit alpha 1, chloroplastic
Small subunit ribosomal protein s27e
Methionine sulfoxide reductase (mss4-like) family protein
60S acidic ribosomal protein family
Ribosomal protein S8e family protein
Solute carrier family 25
Costars family protein
Arabidopsis thaliana responsive to abscisic acid 1b
Nucleosome/chromatin assembly factor group d 02
Ribosomal protein L32e
Ribosomal L22e protein family
P-loop containing nucleoside triphosphate hydrolases superfamily protein
Probable mediator of RNA polymerase II transcription subunit 37c
Ribosomal L29e protein family
Ribosomal protein S7e family protein
Encodes a chloroplast ribosomal protein L2
PCGS3
PCGS4
-1
0
1
Tubulin/FtsZ family protein
Splicing factor, putative
Double-stranded RNA-binding protein 2
S-adenosyl-L-methionine-dependent methyltransferases superfamily protein
Small nuclear ribonucleoprotein family protein
Small nuclear ribonucleoprotein family protein
Microtubule-associated protein RP/EB family member 1C
Clathrin heavy chain 1
Strong similarity to initiation factor eIF-2
Atp binding microtubule motor family protein
TCP-1/cpn60 chaperonin family protein
Actin-related protein 2
F-type h+-transporting atpase subunit a
TCP-1/cpn60 chaperonin family protein
Cullin-3A
P-loop containing nucleoside triphosphate hydrolases superfamily protein
Inositol-hexakisphosphate/diphosphoinositol-pentakisphosphate 1-kinase
Chloroplast ribosome release factor 1
MIF4G domain-containing protein / MA3 domain-containing protein
Chloroplast ribosome release factor 1
U1 small nuclear ribonucleoprotein 70 kDa
Ribosomal protein L1p/L10e family
Cactus-binding carboxy-terminal, cactin
Transducin/WD40 repeat-like superfamily protein
50S ribosomal protein L14, chloroplastic
DNA-directed RNA polymerase II subunit 1
Probable ion channel POLLUX
Calcineurin-like metallo-phosphoesterase superfamily protein
Regulator of nonsense transcripts 1 homolog
V-type h+-transporting atpase subunit a
ARM repeat protein interacting with ABF2
Cullin-1
ATP-dependent zinc metalloprotease FTSH 4, mitochondrial
C2H2-like zinc finger protein
Pre-mRNA-splicing factor ATP-dependent RNA helicase DEAH1
Solute carrier family 25 (mitochondrial citrate transporter), member 1
Arabidopsis thaliana alpha-glucan phosphorylase 2
Structural maintenance of chromosomes (SMC) family protein
Serine-threonine kinase receptor-associated protein
Protein translocase subunit SECA1, chloroplastic
Glucose-6-phosphate isomerase 1, chloroplastic
Cation efflux family protein
Class II aaRS and biotin synthetases superfamily protein
PCGS3
PCGS4
-1
0
1
Transmembrane e3 ubiquitin-protein ligase
Zinc finger C-x8-C-x5-C-x3-H type family protein
Ubiquitin carboxyl-terminal hydrolase 13
3-oxoacyl-[acyl-carrier-protein] synthase II, chloroplastic
Protein kinase superfamily protein
Transcription factor bHLH66
Probable ion channel POLLUX
Double-stranded RNA-binding protein 2
Mitochondrial substrate carrier family protein
Beta galactosidase 9; Putative beta-galactosidase
Chaperone DnaJ-domain superfamily protein
5-enolpyruvylshikimate-3-phosphate phospholyase
Eukaryotic translation initiation factor 3 subunit 7 (eIF-3)
SAC3/GANP/Nin1/mts3/eIF-3 p25 family
SNARE-like superfamily protein
Arm repeat superfamily protein
Class ii aars and biotin synthetases superfamily protein

**Fig. S9** Gene expression networks analysis of differentially expressed genes identified in each of the six paired-wise comparisons of all four seasons of *P. appendiculatum*. Figures (a) to (l) depict the stress-related genes along with the complete gene set enrichment analysis (GSEA) network observed in the six inter-season comparison of DUGS1_vs _DUGS2, DUGS1_vs_DUGS3, DUGS1 _vs _DUGS4, DUGS2_vs _DUGS3, DUGS2_vs _DUGS4 and DUGS3_vs_DUGS4. The color scale (from white, blue to red) is based on (corrected) p-value (5% FDR, p = 0.05). Dark red categories are most significantly overrepresented, white nodes are not significantly overrepresented (NA); they are included to show other nodes in the context of the GO hierarchy.

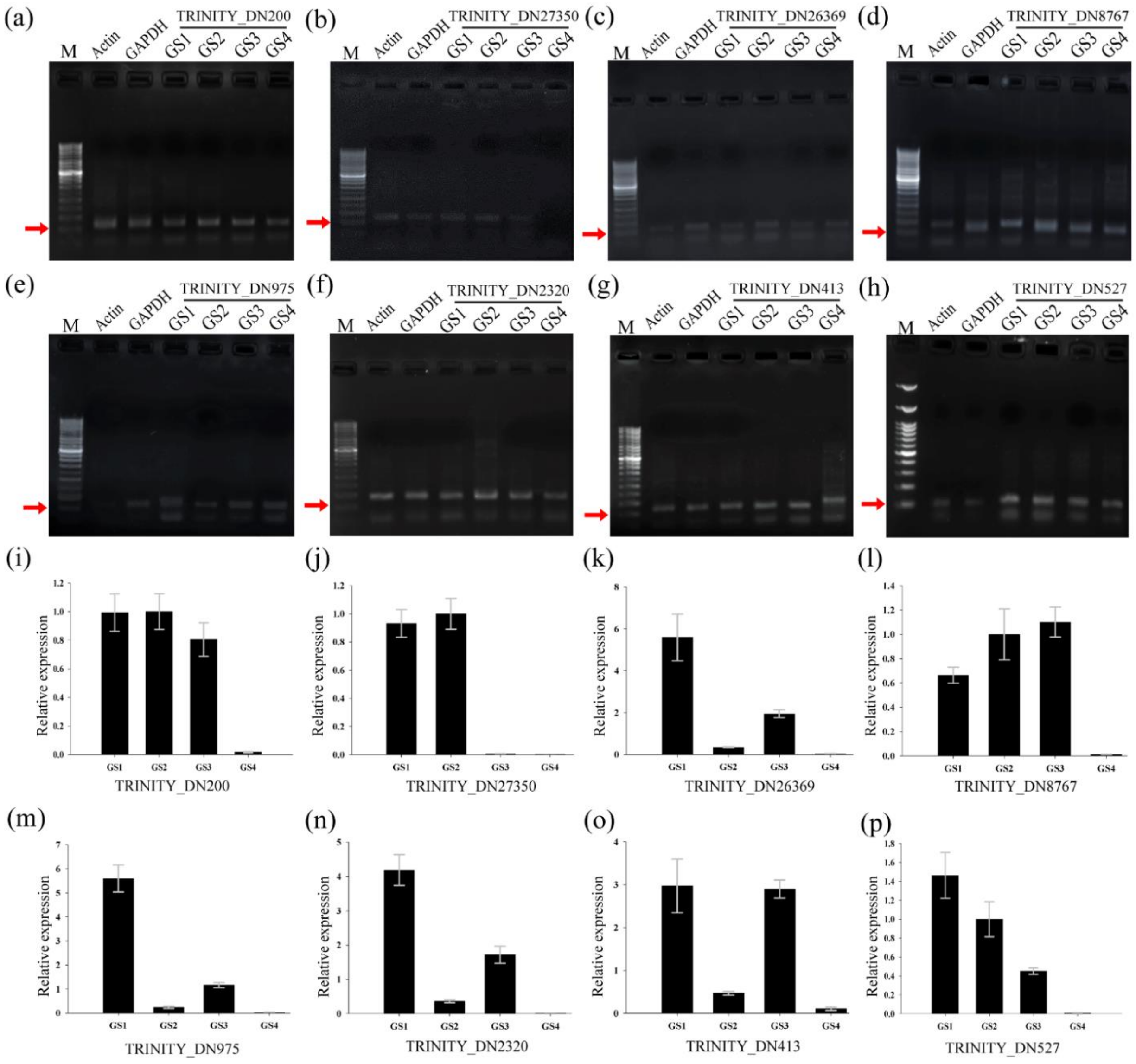


**Fig. S10** Validation of the RNAseq data of *D. hirsuta* by **(a-h)** semi-quantitative PCR (semi-qRT-PCR) and **(i-p)** quantitative real-time PCR (qRT-PCR) for eight selected genes. Expressions of all the transcripts in four different growing seasons (GS) can be observed in the agarose gels at approximately 120-130 bp region. In the qRT-PCR data, the bar graphs have been created on the normalized expression values against internal references. The fold difference of each amplified product in the samples was calculated using the $2^{-\Delta\Delta Ct}$ method. GS1 is pre-monsoon, GS2 is monsoon, GS3 is post-monsoon, and GS4 is fruiting seasons.

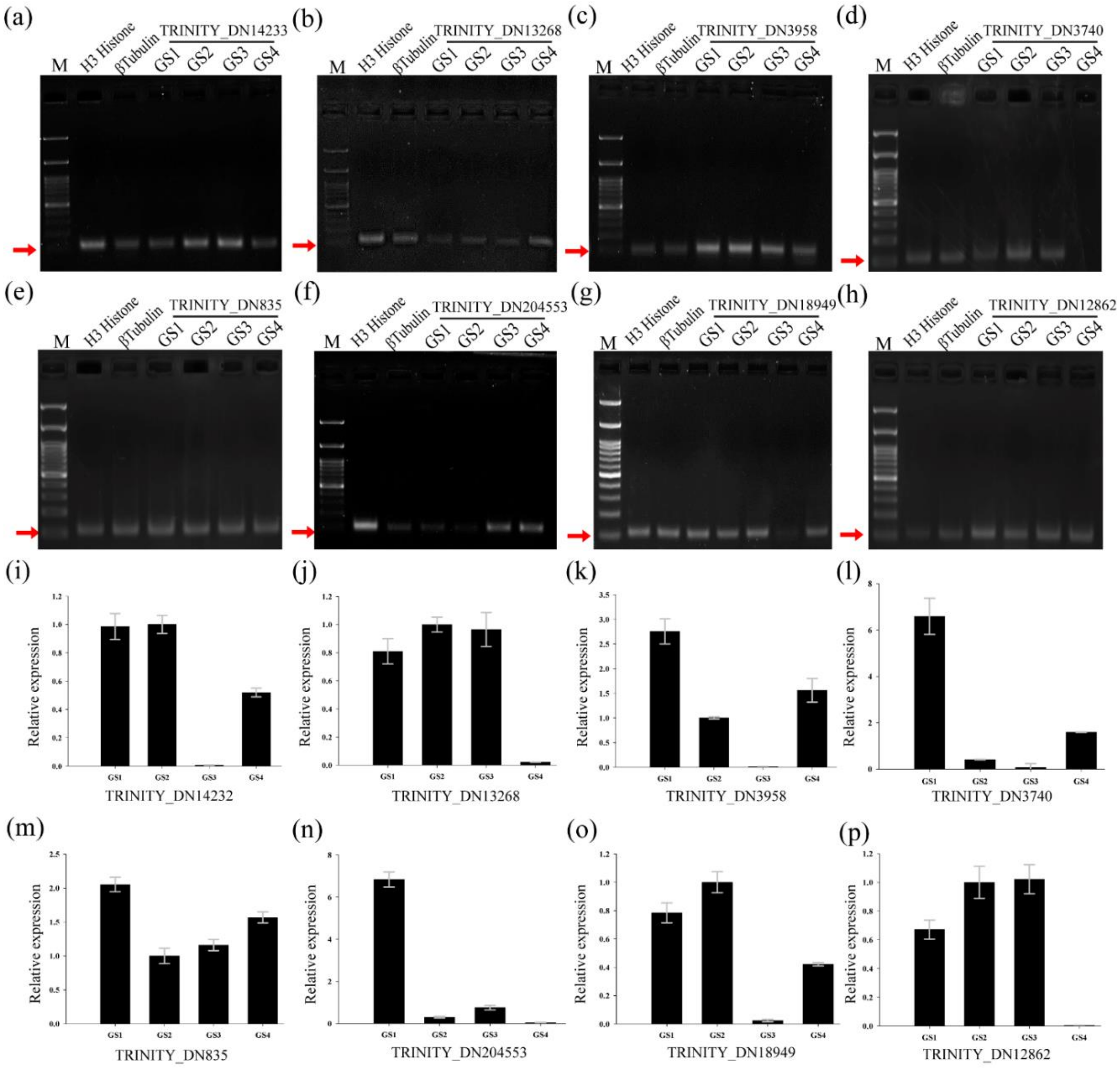


**Fig. S11** Validation of the RNAseq data of *P. appendiculatum* by **(a-h)** semi-quantitative PCR (semi-qRT-PCR) and **(i-p)** quantitative real-time PCR (qRT-PCR) for eight selected genes. Expressions of all the transcripts in four different growing seasons (GS) can be observed in the agarose gels at approximately 120-130 bp region. In the qRT-PCR data, the bar graphs have been created on the normalized expression values against internal references. The fold difference of each amplified product in the samples was calculated using the $2^{-\Delta\Delta Ct}$ method. GS1 is pre-monsoon, GS2 is monsoon, GS3 is post-monsoon, and GS4 is fruiting seasons.

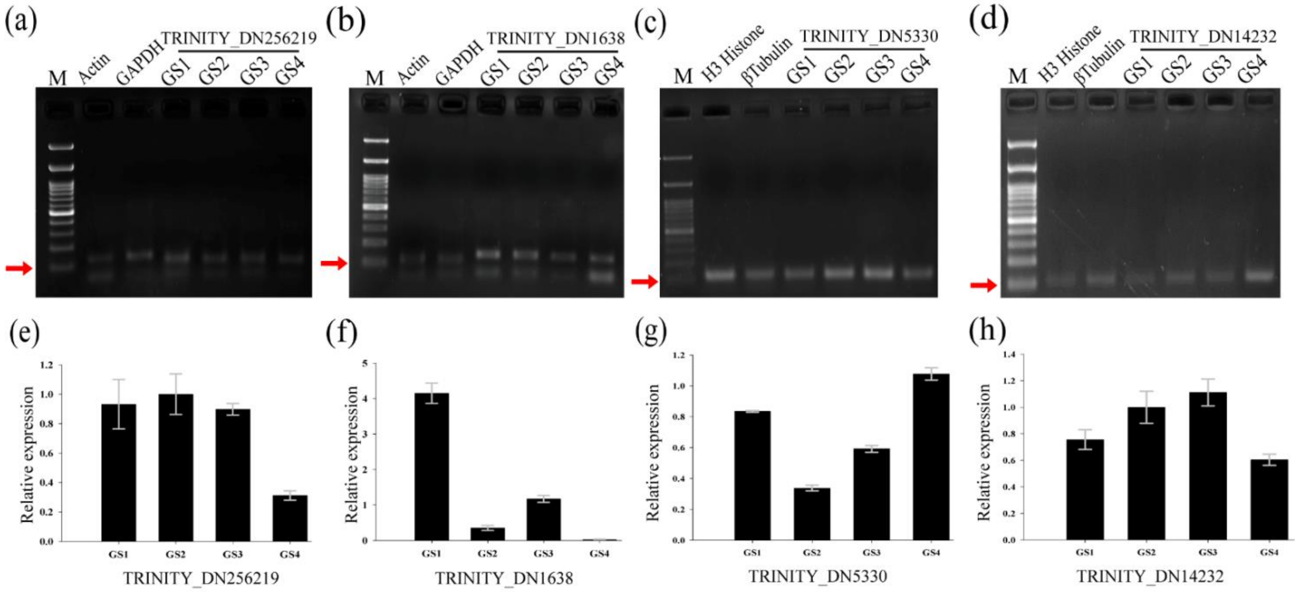


**Fig. S12** Validation of the RNAseq data of **(a**,**b**,**e**,**f)** *D. hirsuta* and **(c**,**d**,**g**,**h)** *P. appendiculatum* by **(a-d)** semi-quantitative PCR (semi-qRT-PCR) and **(e-h)** quantitative real-time PCR (qRT-PCR) for selected transcription factors. The trinity IDs are given above/bellow on gels/bars. Expressions of all the transcripts in four different growing seasons (GS) can be observed in the agarose gels at approximately 120-130 bp region. In the qRT-PCR data, the bar graphs have been created on the normalized expression values against internal references. The fold difference of each amplified product in the samples was calculated using the $2^{-\Delta\Delta Ct}$ method. GS1 is pre-monsoon, GS2 is monsoon, GS3 is post-monsoon, and GS4 is fruiting seasons.
